# Sequence-defined oligophosphoesters for selective inhibition of the KRAS G12D/RAF1 interaction

**DOI:** 10.1101/2024.03.12.584553

**Authors:** Bini Claringbold, Steven Vance, Alexandra R. Paul, Michelle D. Garrett, Christopher J. Serpell

## Abstract

Rat Sarcoma (RAS) genes are the most frequently mutated genes in cancer, with KRAS being the most predominant oncogene, yet they have proved extremely difficult to drug because they operate primarily through protein-protein interactions (PPIs) which lack an obvious pocket for small molecules. Sequence-defined synthetic oligomers could combine the precision and customisability of synthetic molecules with the size requirements to address entire protein-protein interaction surfaces. We have adapted the phosphoramidite chemistry of oligonucleotide synthesis to produce a library of nearly one million non-nucleosidic oligophosphoester sequences – phosphoestamers - and used a fluorescent-activated bead sorting (FABS) process to select oligomers that inhibit the interaction between KRAS^G12D^ (the most prevalent, and undrugged, mutant) and RAF, a downstream effector of RAS whose activation results in cell proliferation. Hits were identified using tandem mass spectrometry, and validation showed effective inhibition with IC_50_ values as low as 25 nM, and excellent selectivity for the mutant over the wild type form. These findings could lead to new drugs against cancers driven by mutant RAS, and provided proof-of-principle for the phosphoestamer platform against PPIs in general.

## Introduction

RAS proteins are small GTPases with a GTP-bound “active” state (RAS-GTP) and a GDP-bound “inactive” state (RAS-GDP)^1^ which they regularly cycle between. When in the active RAS-GTP conformation, RAS interacts with several downstream effector pathways, such as the RAF-MEK-ERK, RalGDS and PI3K-AKT-mTOR pathways.^2,3^ Oncogenic RAS is locked in the GTP conformation, which encourages tumourigenesis.^4,5^ Kirsten Rat Sarcoma (KRAS) is the most frequently mutated of the RAS family of proteins, accounting for approximately 75% of RAS mutations.^6^ Within KRAS, 98% of the mutations are seen at the G12, G13, or Q61 positions,^7^ but the G12D mutation is the most common overall.^8^ KRAS^G12D^ is commonly found in pancreatic,^9^ colorectal,^10^ lung,^11^ and breast cancers;^12^ these cancers have the highest incidence of death^12,13^ and are associated with poor prognosis in patients.^14^

The difficulty in drug discovery for RAS is that the only obvious pocket for a small molecule is occupied by GDP/GTP which are both strongly bound,^15^ and present at high cellular concentrations,^16^ making its replacement inaccessible. The rest of RAS’s activity occurs through protein-protein interactions (PPIs) – these involve large surfaces which are relatively flat and featureless compared with the clefts that medicinal chemists target classically.^17^ In the case of RAS, the interaction surfaces lack even a well-defined interaction surface which could be addressed with compounds such as α-helix mimics.^18^ Nonetheless, small molecule inhibitors have been found which exploit the nucleophilicity of cysteine in the G12C mutant, combined with a less-obvious binding site, which have been approved for non-small cell lung cancer, but this mutant is only present in 12% of such cancers, and resistance has been observed to develop rapidly.^19,20^ While there has been some progress with small molecule G12D inhibitors, such as Mirati Therapeutics MRTX1133,^21^ there is still a great need for more research in this area.

Larger molecules could be used to inhibit PPIs, and indeed there are advances based upon natural sequenced polymers/oligomers – antibodies,^22^ peptides,^23^ and aptamers^24^ – which have been discovered through selection methodologies. The disadvantages of using biomolecular chemistry are that chemical diversity is fundamentally limited, and that it is recognised by endogenous processes, which can result in degradation^25^ and/or immune response.^26^ Synthetic foldamers which can display a programmable set of functional groups can circumnavigate these problems, but are currently best suited as mimics of secondary structures with prior knowledge of which groups should be displayed.^27,28^

Our approach is to create larger synthetic sequence-defined molecules which could cover a significant amount of protein surface area, without bias towards any particular protein substructure. To ensure that uniform macromolecules (as opposed to disperse polymers) can be obtained, we are adapting the automated phosphoramidite chemistry used in oligonucleotide synthesis^29,30^ which is capable of >150 couplings,^31^ but employing non-nucleosidic monomers, to obtain phosphoestamers: abiotic, uniform oligo- or polyphosphoesters.^32–34^

We herein report the synthesis of a phosphoestamer library, and identification of oligomers through selection which disrupt the interaction between KRAS^G12D^ in its GTP form and RAF1-RBD with IC_50_ values as low as 25 nM. The stringent selection process means the three oligomers are selective and do not bind either the equivalent wild-type KRAS, nor the GDP-hosting form. These results demonstrate that phosphoestamers can be effective at blocking medically important protein-protein interactions and could lead to transformative new drug development programmes.

## Results

### General Overview of the strategy

Our route to selection of phosphoestamers for PPI inhibition (Fig. 1) has five key steps: (1) choice and synthesis of phosphoramidite monomers; (2) synthesis of the one-bead-one-sequence library; (3) multiple rounds of fluorescence-activated bead sorting (FABS) for the selection of oligomers that disrupt PPIs; (4) Sequencing of selected oligomers by LC-MS/MS; and (5) resynthesis and validation of selected oligomers.

**Figure 1.**
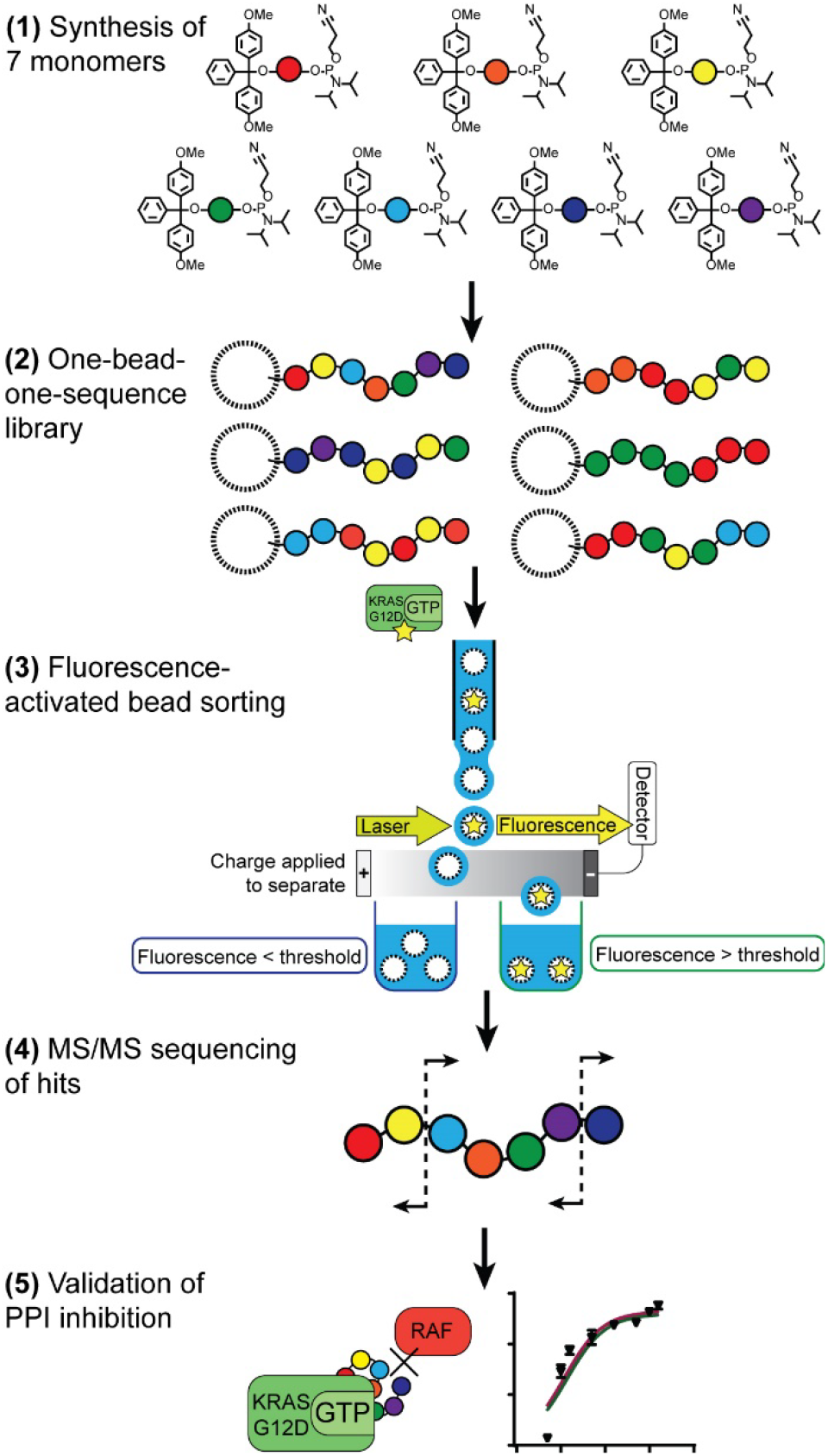
Overview of oligomer synthesis strategy.

### Monomer synthesis

Seven phosphoramidite monomers (Fig. 2) were selected for use in the oligomer library. The synthesis requires monomers to be based upon diols which are then protected at one hydroxyl with a dimethoxytrityl (or trityl if phenolic) group, followed by activation at the second using 2-cyanoethyl *N,N*-diisopropylchlorophosphoramidite (full procedures and data, Supporting Information Section 2.2). This yield monomers which can be linked using standard automated oligonucleotide synthesis chemistry. **BPA** (based upon the diol bisphenol A) and **C12** (dodecanediol) provide hydrophobic regions within the oligomer, with **BPA** being rigid while **C12** is flexible. **HEG** (hexaethylene glycol) is hydrophilic. Patterns of **C12** and **HEG** have been shown to direct supramolecular chemistry in polyphosphoesters.^33,35,36^ **cSS** (cyclic di-serine) and **cYY** (cyclic di-tyrosine) are diketopiperazines based upon amino acids which form a rigid structure and are able to act as both hydrogen bond donors and acceptors.^37,38^ **NDI** (naphthalene diimide) and **DAN** (dialkoxynaphthalene) are capable of π-π interactions, and in particular form a donor (DAN)/acceptor (NDI) pair which also enables folding,^39^ including in oligophosphoesters.^34,40^ All phosphoramidite monomers were successfully synthesised, except **C12** and **HEG** which were available commercially.

**Figure 2.**
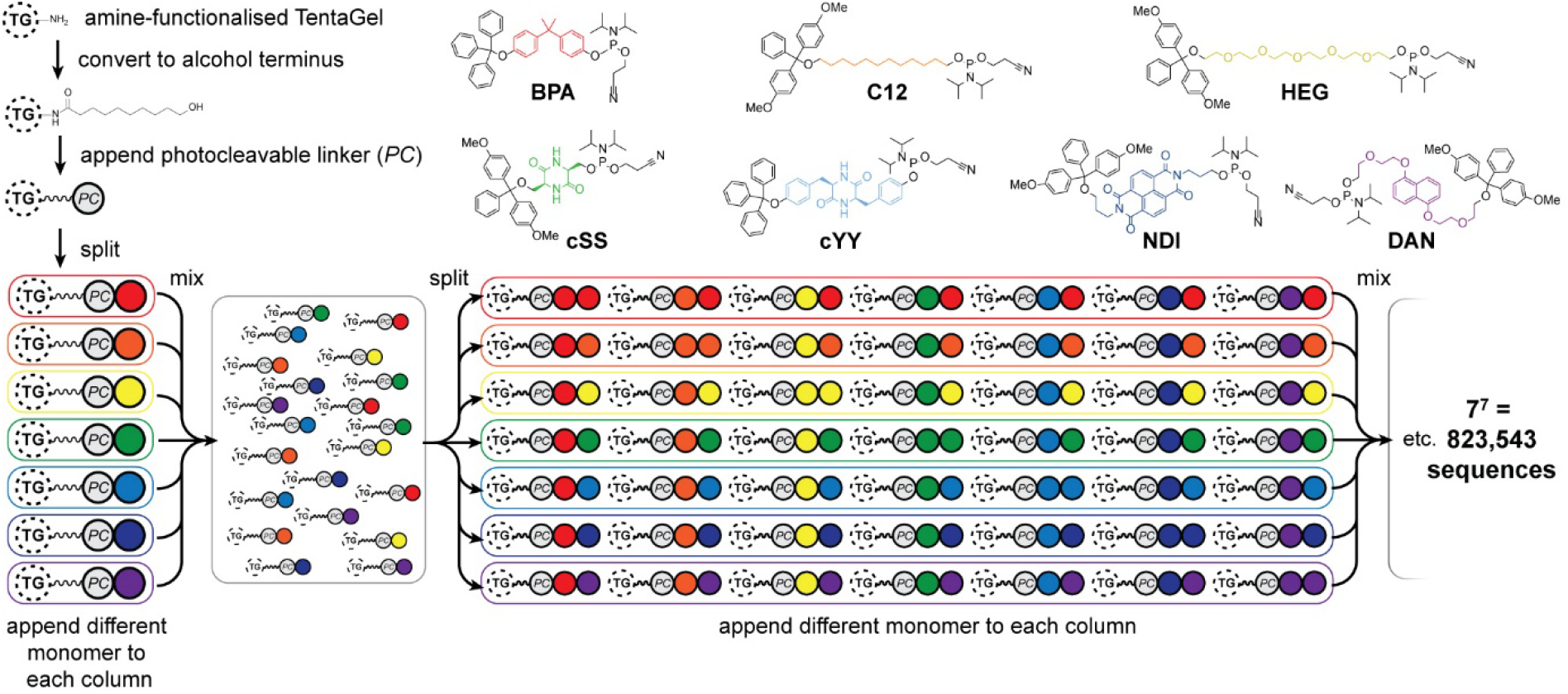
Phosphoramidites used for the oligomer library, and preparation of the library on TentaGel® (TG) beads with a photocleavable (PC) linker by split-and-mix synthesis.

### Synthesis of Library

The one-bead-one-sequence phosphoestamer library was constructed using split-and-mix techniques^41^ (Fig. 2, Supporting Information Sections 2.3-2.4). Creating 7mers of all the combinations of the monomers produced 7^7^ = 823,543 unique full-length sequences, plus any sequences that did not go to completion.^42^ Synthesis of the library was completed using automated phosphoramidite synthesis on TentaGel® M NH_2_ Monosized Amino TentaGel Microspheres (TG-beads); TG-beads have a polystyrene backbone with a PEG spacer and are chemically inert, making them suitable to phosphoramidite addition.^43^ The TG-beads were modified with 10-hydroxydecanoic acid to create hydroxy TG-beads and were swelled in dichloromethane before a photocleavable linker was attached to allow for photocleavage from the TG-beads after fluorescent selection.^44^ The beads were then split for the first round of monomer addition. After each monomer is added to an individual pool, the library was mixed and split out again for the second monomer addition, creating 49 different combinations in the second step, before being mixed again as the cycle continues. The resultant oligomer library contained over 200 million individual TG-beads, giving on average 268 beads per sequence, each displaying 10^11^ copies of that specific phosphoestamer sequence (Fig. 2). The trityl monitor was used to monitor the efficiency of each coupling.

### Fluorescent-activated bead sorting (FABS)

Fluorescent-activated bead sorting (FABS) is a methodology that allows for the selection of the best binding oligomers using a flow cytometer. Flow cytometry has previously been used for the selection and optimisation of aptamers^42,45,46^ and inhibitors of small GTPases such as Rho and Rab.^47^ We used several selection steps to identify phosphoestamers that bind to KRAS^G12D^-GMPPnP and disrupt any interaction between KRAS^G12D^-GMPPnP (a non-hydrolysable analogue of GTP) and RAF1-Ras binding domain (RBD). Protein production details are in Supporting Information Section 3. Using fluorescently labelled proteins, measured bead fluorescence correlates with the binding affinity of the strands on that bead for the protein being tested. Gating can then be used to separate beads above or below any chosen fluorescent intensity, indicating higher or lower binding by the sequence on that bead. The proteins used in the FABS selection were expressed with a biotin tag, which was then used as a linker to fluorophore-labelled streptavidin (STV). KRAS^G12D^-GMPPnP and KRAS^G12D^-GDP were tagged with fluorescein-STV, and RAF1-RBD with rhodamine Red™-X-STV.

Selection of the phosphoestamer library for KRAS^G12D^-GTP/RAF1-RBD PPI inhibition employed four rounds of separations (full data and analysis, Supporting Information Section 4). In Round 1 the oligomer library was incubated with enough fluorescein-tagged KRAS^G12D^-GMPPnP to cover 4% of the oligomer library; only oligomers with a high affinity for KRAS^G12D^-GMPPnP would therefore acquire fluorescence that can be detected, and thus and retained for round 2 (Fig. 3a). After the first selection round, 48,169 beads (of the original 2 × 10^8^) were retained, and the KRAS^G12D^-GMPPnP was removed from the selected portion of the library by washing. The pool was then incubated with fluorescein-tagged KRAS^G12D^-GDP. Beads which display a strong fluorescent signal would be bound to KRAS^G12D^-GDP, and were therefore removed (since KRAS-GDP does not interact with RAF) in this round of FABS (Fig. 3b), leaving 12,111 library beads. The KRAS^G12D^-GDP was removed by washing, and the remaining library was incubated with fluorescein-tagged KRAS^G12D^-GMPPnP and a 3-fold excess of RAF1-RBD (rhodamine Red™-X tagged). Here, any oligomers from the library that had a high fluorescent signal using the 585/29 and 600 nm bandpass filters either had an affinity for RAF1-RBD itself, or – more likely given prior rounds – bound KRAS^G12D^-GTP in a manner which did not prevent the GTPase from also binding RAF1-RBD. Conversely, those beads which did not acquire rhodamine fluorescence must inhibit the PPI since their binding of KRAS was selected for in the first round (Fig. 3c); this was validated through checking fluorescein fluorescence, which gave high readings (Fig. S44, Supporting Information). 676 library beads remained for the final round; still far too many to practically sequence. A fourth round was used to select only those with highest affinity: the library was washed again to remove any remaining proteins and incubated with enough fluorescein-tagged KRAS^G12D^-GMPPnP to cover 50% of the remaining library. For the final selection, 200 beads of the highest fluorescence were sorted such that each individual bead was placed in an individual well of a well plate (Fig. 3d). The oligomers were then cleaved from the TG-beads *via* the photocleavable linker.

**Figure 3.**
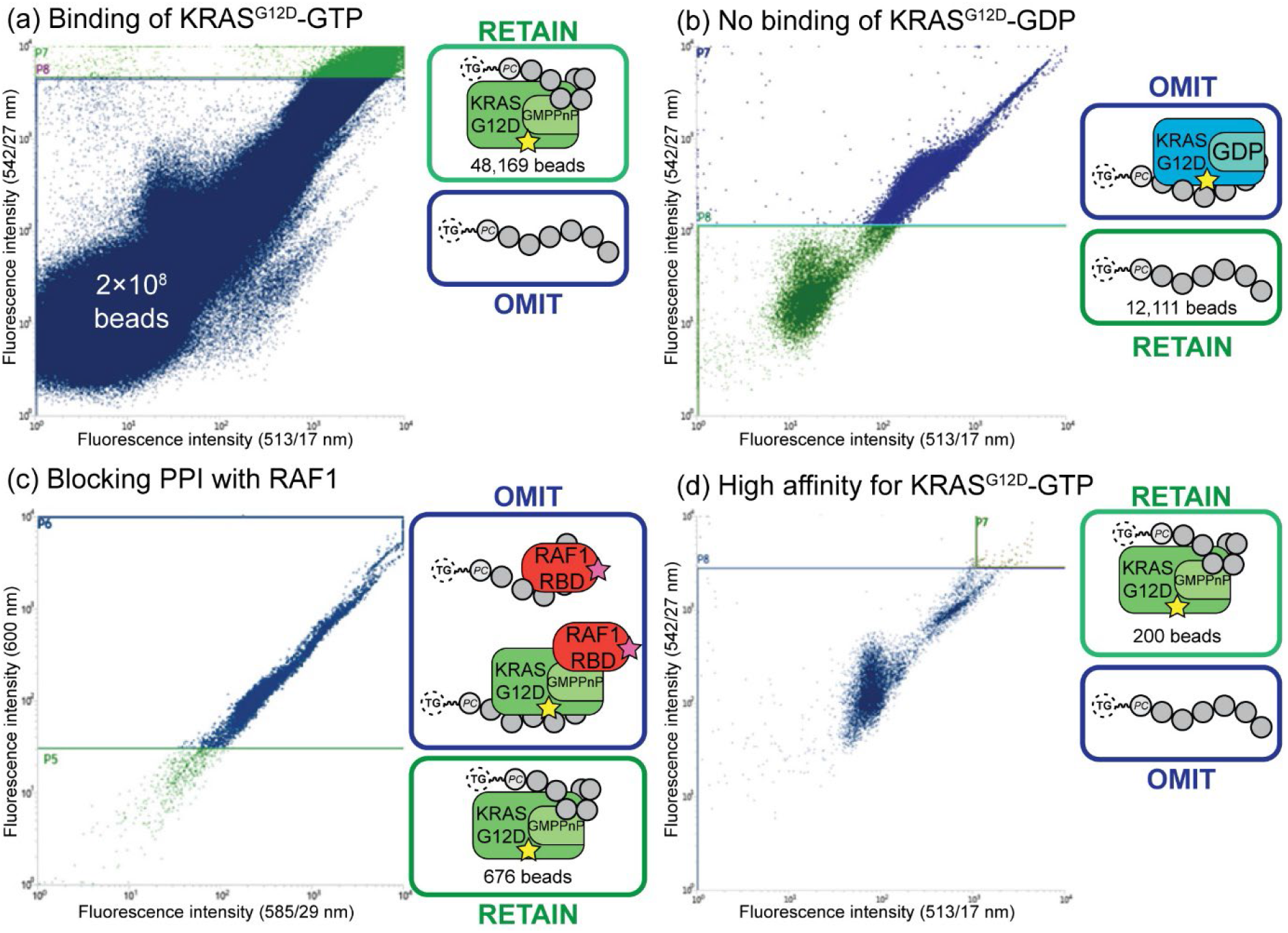
FABS selection of top binding oligomers. Yellow star = fluorescein label; pink star = rhodamine label. (a) Initial selection step, selecting for oligomers that bind to KRAS^G12D^-GMPPnP. (b) Second selection step, selecting for oligomers that do not bind to KRAS^G12D^-GDP. (c) Third selection step, selecting for oligomers that do not bind to RAF1-RBD. (d) Final selection step, selecting top oligomers that bind to KRAS^G12D^-GMPPnP.

### Sequencing of hit phosphoestamers by mass spectrometry

Oligomers were sequenced and identified with a Q-TOF nanospray LC-MS/MS method. A commercially purchased 7-base DNA oligomer was used to identify the limit of detection and observe patterns in how oligomers of this length could fragment. These results showed the oligomers were most likely to be detected as [M-2H]^2-^ parent ions, and MS/MS identified c- and y-ions^48^ as the most predominant. Of the 200 top oligomers selected from FABS, 21 (according to capacity) were prepared for LC-MS/MS analysis and 6 oligomers (**O1-O6**) produced viable data which could be interpreted (Table 1, full data analysis Supporting Information Section 5).

**Table 1.** Oligomer masses detected *via* LC-MS/MS.

| Oligomer | m/z detected | Neutral molecular mass (Da) |
| --- | --- | --- |
| **O1** | 854.978 $[M-2H]^{2-}$<br>1710.963 $[M-H]^-$ | 1,711.970 |
| **O2** | 992.314 $[M-2H]^{2-}$ | 1,986.642 |
| **O3** | 911.342 $[M-2H]^{2-}$ | 1,824.698 |
| **O4** | 971.321 $[M-2H]^{2-}$ | 1,944.656 |
| **O5** | 971.288 $[M-2H]^{2-}$ | 1,944.590 |
| **O6** | 700.120 $[M-2H]^{2-}$ | 1,402.240 |

MS/MS data revealed molecular ions which fell within the expected oligomer library range (1669.08 – 3125.43 Da), with the exception of **O6** (M = 1402.24) which represents a truncation. Data from **O1** showed not only the common [M-2H]^2-^ parent ion, but also a smaller [M-H]^-^ ion at 1710.693 m/z; this provided two separate sets of MS/MS data that could be analysed and compared when identifying the sequence, assisting in validation of the workflow. Sequencing was performed using RoboOligo, a programme designed for the analysis of tandem mass spectrometry data of oligonucleotides.^49^

Figure 4 shows the RoboOligo analysis of **O2**, which was identified as **NDI-C12-C12-C12-NDI-BPA**. Examining all the sequences selected (Figure 5), there were some common patterns identified, such as the multiple adjacent monomers of both **C12** in **O2** and **O4**, and of **HEG** in **O1** and **O3**. Every initial monomer used was found in at least one phosphoestamer sequence, except **cSS** which was not seen in any top binder analysed. Of the sequences identified only one (**O4**) was a full-length 7-mer, with **O1**-**O5** being 5/6mer oligomers and **O6** being a tetramer. Given that monitoring trityl groups during synthesis showed that the couplings were successful to the end, and having used a redundancy of 268, at least one instance of each full-length sequence should be present. It is therefore likely that these smaller oligomers are better binders compared to the 7mers. We have observed the selection of optimal sequences arising from synthetic inefficiencies previously.^42^

**Figure 4.**
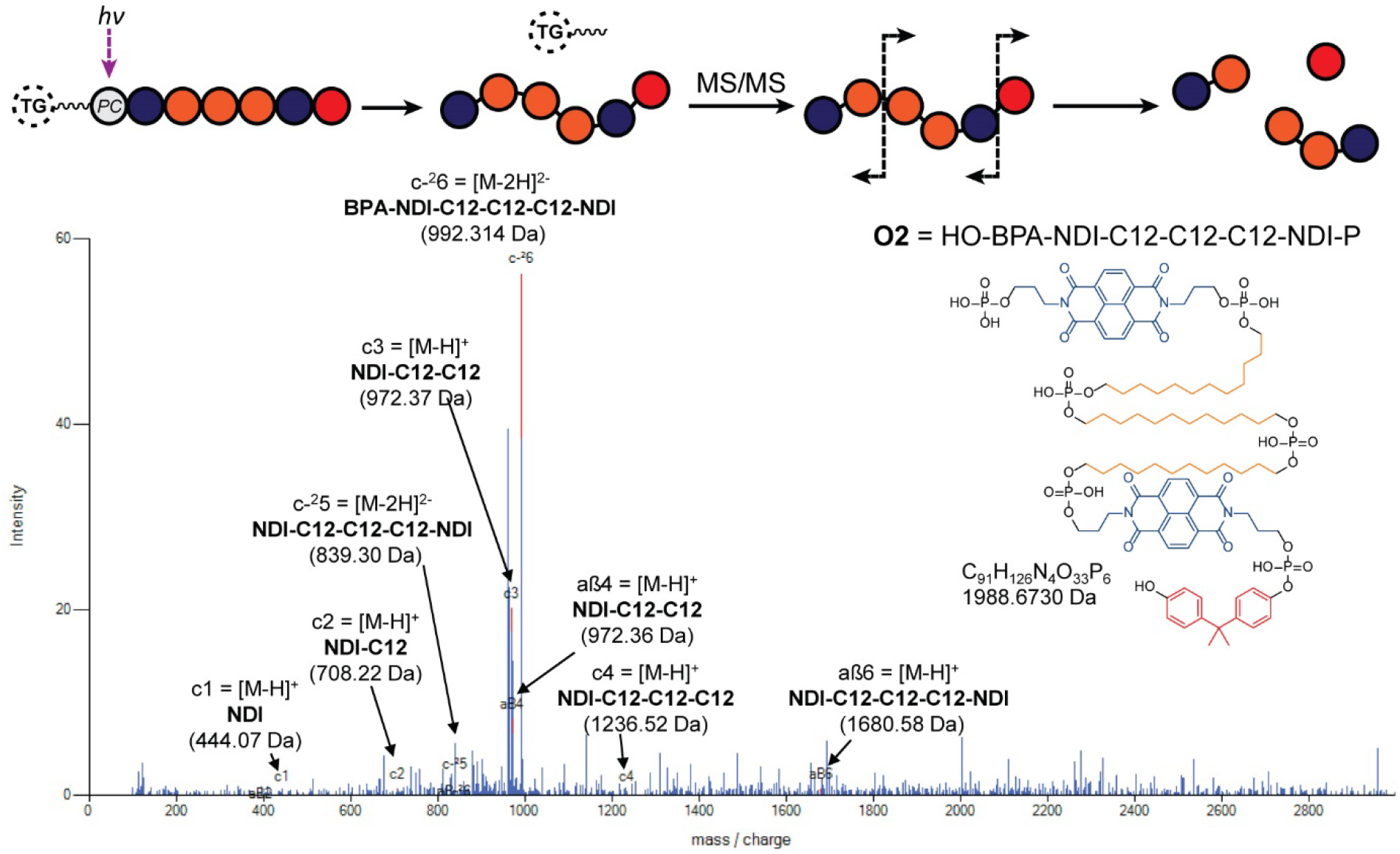
RoboOligo analysis of **O2**. Other major peaks also indexed on alternative fragments (see Supporting Information Section 5)

**Figure 5.**
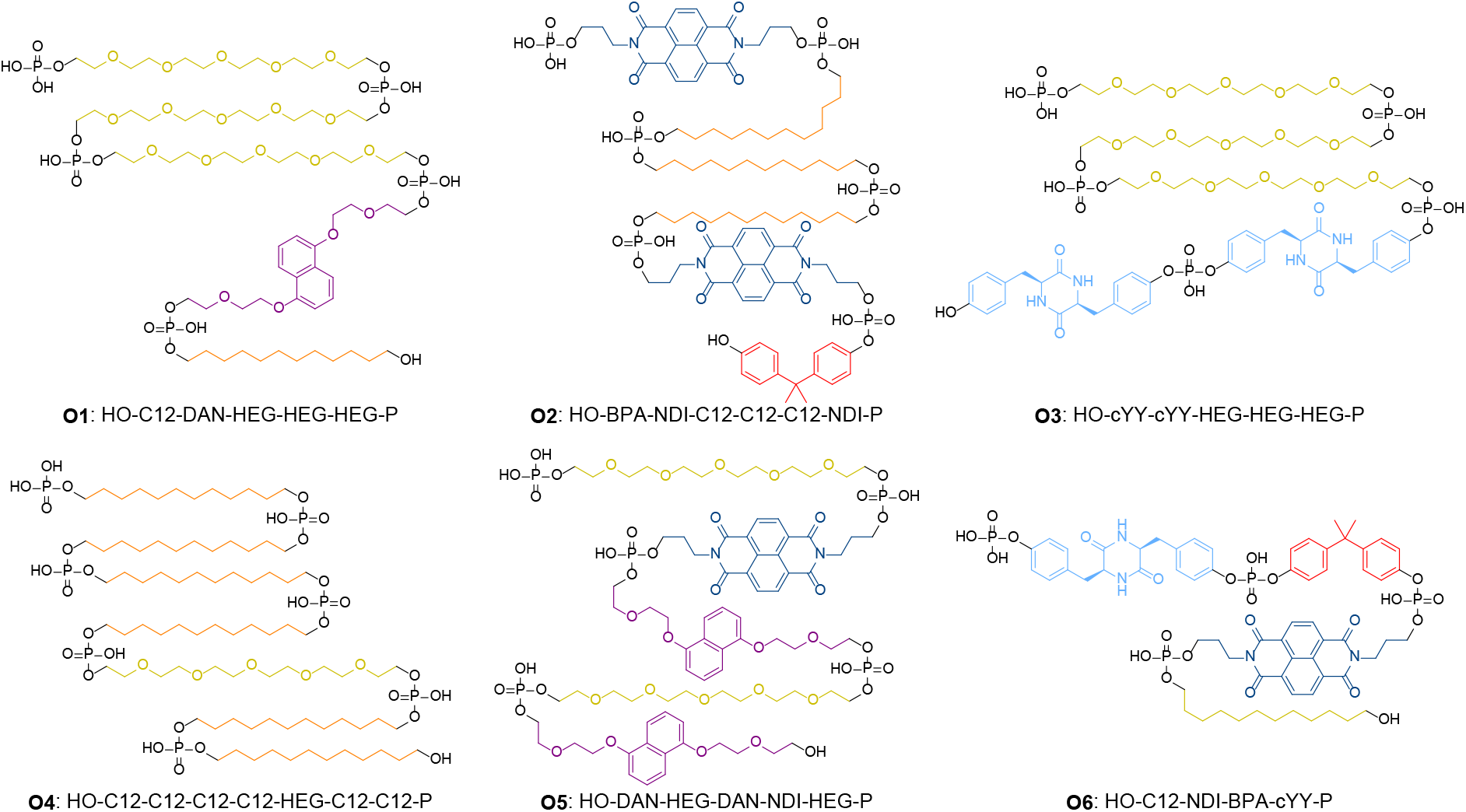
Sequences of top binding phosphoestamers. Sequences are given by analogy with nucleic acid conventions (5’ to 3’) meaning that the first monomer listed is the last one added during chemical synthesis. This is evident from the location of the terminal phosphate which exists in the MS/MS spectra as a result of the photocleavage reaction, leading to the HO- and -P (phosphate) termini.

### Validation of PPI inhibition by phosphoestamers

Oligomers **O1** – **O6** were remade using the DNA synthesiser and the yields were determined by manually cleaving the final DMT protecting group on the oligomers and quantifying the DMT cation by UV-visible spectroscopy (Supporting Information Section 2.4). The achievement of desired length and purity was confirmed by polyacrylamide gel electrophoresis (Fig. 6a). The remade oligomers were purified of small molecules using C18 spin tips. To ensure the assay was viable, a positive control **Ch-3** (Fig. 6a) known to disrupt KRAS^G12D^ interactions^50^ was synthesised. An assay was developed (Supporting Information Section 6) in which polystyrene well plates were coated by overnight incubation with KRAS^G12D^-GMPPnP. The well plate was washed with a blocking solution before incubation with oligomers and RAF1-RBD-GFP. Any KRAS sites not blocked by the oligomers would interact with the RAF1-GFP and result in a fluorescent signal which would be detected. In this assay, we first established that both KRAS-GMPPnP and RAF1-RBD-GFP were required to give a fluorescence signal. The assay was then conducted at varying concentrations of **Ch-3**, giving a resultant IC_50_ of 6.35 ± 0.20 μM (Fig. 6b), consistent with reported cell viability assays performed with the same compound.^50^

**Figure 6.**
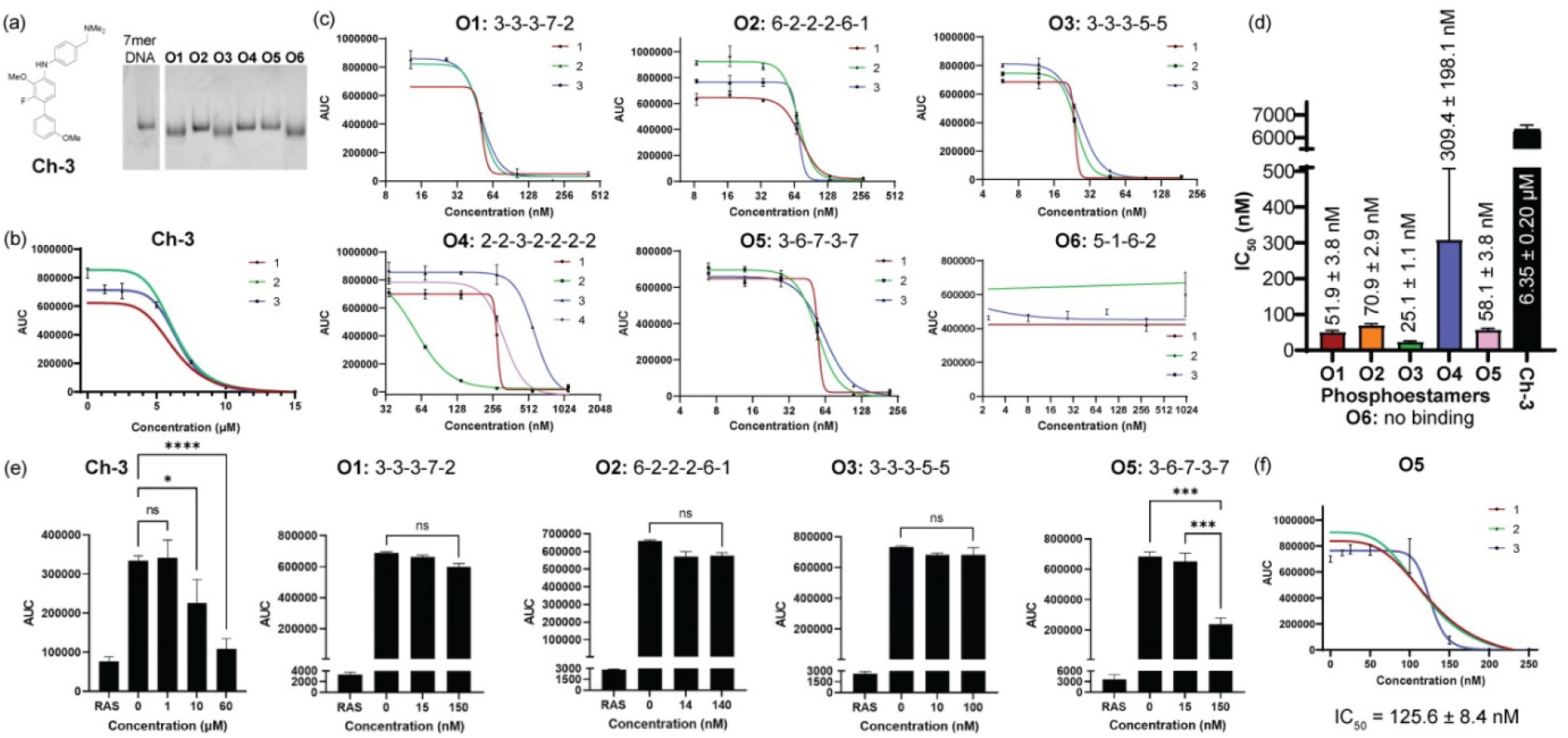
(a) Positive control compound **Ch-3**, and pure phosphoestamers **O1**-**O6** characterised by polyacrylamide gel electrophoresis. (b) KRAS^G12D^/RAF1-RBD interaction assay results for **Ch-3**. Data collected using area under the curve (AUC) of GFP emission spectra between 490 and 540 nm. Numbers 1-3 indicate biological repeats. (c) KRAS^G12D^/RAF1-RBD interaction assays results for **O1**-**O6**. (d) IC_50_ values for **Ch-3** and **O1**-**O6** calculated from assays. (e) Effect upon KRAS^WT^/RAF1-RBD interaction of **Ch-3** and top scoring phosphoestamers screened at concentrations up to 3 × IC_50_ for KRAS^G12D^/RAF1-RBD, and (f) full measurement of IC_50_ for **O5**, the only phosphoestamer which showed an effect.

Conducting the same assays with the phosphoestamers (Fig. 6c, d) showed that **O1, O2, O3** and **O5** had IC_50_ values data at the nanomolar level: 20 – 250 times smaller than the positive control **8. O6** showed no change in fluorescent signal across three repeats, and so is unlikely to have any effect on the PPIs between KRAS and RAF. **O6** is the smallest oligomer, and it is possible that the MS/MS has only detected a fragment of a whole chain that does not successfully disrupt the PPIs on its own, or that some multivalency effect which was in operation on the beads but cannot work with isolated strands.

**O3** had the lowest IC_50_ value, at 25.14 ± 1.06 nM; suggesting the strongest affinity for KRAS^G12D^-GMPPnP. **O1** had a similar structural motif to **O3** and had the second lowest IC_50_ at 51.94 ± 3.75 nM. The **3-3-3** sequence could be key to improving binding to KRAS^G12D^-GMPPnP compared to the other oligomers. **O5** had a similar IC_50_ to **O1**, 58.08 ± 3.78 nM, but the only similarity between these two is a **3-7** subsequence within the oligomers. **O2** was the only 6-mer and had a higher IC_50_, 70.87 ± 2.88 nM, which could suggest that 5-mer oligomers do have a better affinity to KRAS proteins compared to the longer chains. The only full length (7-mer) oligomer was **O4**, and like **O2** had a much larger IC_50_, 309.38 ± 198.09 nM; the standard deviation was large meaning this oligomer potentially does not bind or disrupt interactions consistently – this is unsurprising given its very flexible nature, which may only bind when multivalency is provided on a bead surface.

Since four oligomers were determined to have consistent dissociation activity between KRAS^G12D^-GMPPnP and RAF1-GFP, we then used the same assay to determine whether these oligomers would have any activity against KRAS^WT^-GMPPnP or whether they would be selective for the mutant form (Fig. 5e). Testing ‘high’ and ‘low’ concentrations of **O1, O2, O3**, and **O5** (approximated as IC_50_ ÷ 3 and IC50 × 3 respectively), it was found that only **O5** showed a decrease in fluorescent signal as the phosphoestamer was added, indicating disruption of the KRAS^WT^-GMPPnP/RAF1-GFP interaction, and weaker selectivity. An IC_50_ was determined for **O5** at 125.61 ± 8.45 nM (Fig. 5f), more than twice that of the value for KRAS^G12D^-GMPPnP, meaning that even **O5** has some selectivity for KRAS^G12D^-GMPPnP over the WT.

Overall, the FABS selection process was successful in providing potential inhibitors of KRAS^G12D^ PPIs, with these oligomers having a much stronger binding affinity compared to the positive control used. Additionally, three of these oligomers are selective between KRAS^WT^ and KRAS^G12D^.

## Discussion

We have combined several different techniques – phosphoramidite synthesis, one-bead-one-compound library synthesis, fluorescence-activated sorting, and tandem mass spectrometry – to create a unique methodology for the selection of novel oligomers that selectively inhibit protein-protein interactions between mutant KRAS^G12D^ and RAF1. The top three final targets (**O1, O2** and **O3**) were inactive against KRAS^WT^ but inhibited KRAS^G12D^ between 25 and 58 nM.

Mutations in RAS cause overactive cell signalling, driving 30% of cancers including ~95% of pancreatic, 32% of lung adenocarcinoma, and 45% of colorectal cancers,^51^ and it stands as an extremely important target in cancer therapy.^52^ Current examples into KRAS^G12D^-GTP inhibitors work at between 180 nM and 6 μM^50,53,54^, so our methodology has produced highly active compounds. MRTX1133, a highly optimised G12D drug in clinical trials has an IC_50_ of 5 nM, but binds to the inactive GDP-form of the protein.^21^ We have not undertaken any chemical optimisation of phosphoestamers, but nonetheless have obtained a number of strong inhibitors. Our mechanistic aim differs in that our selection was set up not for binding to a particular site, but for blocking of a specific PPI – in this case one which only the GTP form participates in. This is important because it means that in principle, our method could be used to generate phosphoestamers addressing any other PPI of interest – opening up access to modulating mechanisms in diseases as diverse as neurodegeneration and viral infection.

Compared to other selection methods, once our protocols were developed, the entire selection process was quick and efficient. With monomers in hand, the oligomer library can be synthesised by a well-trained bench scientist in one week (or quicker if synthesising smaller oligomers), the FABS selection completed in one week, and the analysis of selected hits takes two-to-three weeks. Within one to two months the entire selection process can be completed, whereas methods like HTS and SELEX can take months of analysis and validation.^55–57^

There are some limitations within this framework however, as the method is heavily dependent upon the makeup of the library. Our library contained over 800,000 sequences, and there is the potential to synthesise over 10 million unique combinations using the same system. However, a larger pool of monomers or the synthesis of larger oligomer chains can result in an exponential increase in the materials needed for synthesis and the overall analysis time. Another challenge is the sensitivity of the mass spectrometry – at such small concentrations the instrument available to us was working close to the limit of detection with our analysis, and as such some hits analysed could have been below the detectable limit. More advanced mass spectrometry could enable much deeper sequencing of hits would then lead to better understanding of sequence/property relationships. It must be recognised that the molecules selected are not classically ‘drug-like’ in their size, polarity, or rigidity, and this may limit their use as drugs in their current form. This is particularly a question for RAS targets, since the proteins are located on the interior of the cell membrane. However, the drug discovery community is increasingly aware of the value of drugs outside of Lipinski’s rule of 5, such as peptides, macrocycles, and degraders.^58^ Indeed, phosphoestamers are most physicochemically reminiscent of oligonucleotides, which can enter cells through simple chemical modifications^59^ or a variety of cell entry strategies,^60^ making nucleic acid therapeutics an area of intense research. Such strategies could be employed here.

## Conclusion

We have synthesised and screened a library of phosphoestamers for inhibition of the undrugged mutant KRAS^G12D^-GTP/RAF interaction, through fluorescence-activated bead sorting, identifying six novel oligomers through tandem mass spectrometry analysis. Validation assays showed that three of these oligomers show both an affinity to KRAS^G12D^ that disrupts the interaction with RAF1, and does not affect the equivalent PPI in the wild type protein. This affinity is an improvement upon previously synthesised inhibitors and as well as providing leads for development of new types of drugs, and proves this method could be used to identify inhibitors against other difficult protein-protein interactions.

## Acknowledgements

This work was funded by the Rosetrees Trust PhD Project Grant M743 and PhDPlus Project PhD2022\100050.

## Notes

### Competing Interest Statement

The authors have declared no competing interest.

### Summary of Updates

ORCIDs have been added for remaining authors in the BioRxiv system. No change to the manuscript itself.

## References

1. F. McCormick, K-Ras protein as a drug target, J. Mol. Med., 2016, 94, 253–258.

2. L. Huang, F. Hofer, G. S. Martin and S. H. Kim, Structural basis for the interaction of Ras with RalGDS, Nat. Struct. Biol., 1998, 5, 422–426.

3. H. Feng, Y. Zhang, P. H. Bos, J. M. Chambers, M. M. Dupont and B. R. Stockwell, K-RasG12D Has a Potential Allosteric Small Molecule Binding Site, Biochemistry, 2019, 58, 2542–2554.

4. L. Huang, Z. Guo, F. Wang and L. Fu, KRAS mutation: from undruggable to druggable in cancer, Signal Transduct. Target. Ther., 2021, 6, 1–20.

5. A. M. Waters and C. J. Der, KRAS: The Critical Driver and Therapeutic Target for Pancreatic Cancer, Cold Spring Harb. Perspect. Med., 2018, 8, a031435.

6. E. Sheffels and R. L. Kortum, The Role of Wild-Type RAS in Oncogenic RAS Transformation, Genes, 2021, 12, 1–18.

7. S. Lu, H. Jang, R. Nussinov and J. Zhang, The Structural Basis of Oncogenic Mutations G12, G13 and Q61 in Small GTPase K-Ras4B, Sci. Rep., 2016, 6, 21949.

8. P. D. Chatani and J. C. Yang, Mutated RAS: Targeting the “untargetable” with T cells, Clin. Cancer Res., 2020, 26, 537–544.

9. S. Lanfredini, A. Thapa and E. O. Neill, RAS in pancreatic cancer, Biochem. Soc. Trans., 2019, 1–12.

10. R. A. Burge and G. A. Hobbs, in RAS: Past, Present and Future, eds. J. P. O’Bryan and G. A. Piazza, Academic Press, 1st edn., 2022, pp. 29–63.

11. K. Wood, T. Hensing, R. Malik and R. Salgia, Prognostic and predictive value in KRAS in non-small-cell lung cancer, JAMA Oncol., 2016, 2, 805–812.

12. N. Zeng and J. Xiang, Detection of KRAS G12D point mutation level by anchor-like DNA electrochemical biosensor, Talanta, 2019, 198, 111–117.

13. B. Papke and C. J. Der, Drugging RAS: Know the enemy, Science, 2017, 355, 1158–1163.

14. Q. He, Z. Liu and J. Wang, Targeting KRAS in PDAC: A New Way to Cure It?, Cancers, 2022, 14, 1–16.

15. J. John, R. Sohmen, J. Feuerstein, R. Linke, A. Wittinghofer and R. S. Goody, Kinetics of interaction of nucleotides with nucleotide-free H-ras p21, Biochemistry, 1990, 29, 6058–6065.

16. T. W. Traut, Physiological concentrations of purines and pyrimidines, Mol. Cell. Biochem., 1994, 140, 1–22.

17. M. R. Arkin, Y. Tang and J. A. Wells, Small-Molecule Inhibitors of Protein-Protein Interactions: Progressing toward the Reality, Chem. Biol., 2014, 21, 1102–1114.

18. J. M. Davis, L. K. Tsou and A. D. Hamilton, Synthetic non-peptide mimetics of [small alpha]-helices, Chem. Soc. Rev., 2007, 36, 326–334.

19. M. M. Awad, S. Liu, I. I. Rybkin, K. C. Arbour, J. Dilly, V. W. Zhu, M. L. Johnson, R. S. Heist, T. Patil, G. J. Riely, J. O. Jacobson, X. Yang, N. S. Persky, D. E. Root, K. E. Lowder, H. Feng, S. S. Zhang, K. M. Haigis, Y. P. Hung, L. M. Sholl, B. M. Wolpin, J. Wiese, J. Christiansen, J. Lee, A. B. Schrock, L. P. Lim, K. Garg, M. Li, L. D. Engstrom, L. Waters, J. D. Lawson, P. Olson, P. Lito, S.-H. I. Ou, J. G. Christensen, P. A. Jänne and A. J. Aguirre, Acquired Resistance to KRASG12C Inhibition in Cancer, N. Engl. J. Med., 2021, 384, 2382–2393.

20. N. Tanaka, J. J. Lin, C. Li, M. B. Ryan, J. Zhang, L. A. Kiedrowski, A. G. Michel, M. U. Syed, K. A. Fella, Sakhi, I. Baiev, D. Juric, J. F. Gainor, S. J. Klempner, J. K. Lennerz, G. Siravegna, L. Bar-Peled, A. Hata, R. S. Heist and R. B. Corcoran, Clinical Acquired Resistance to KRASG12C Inhibition through a Novel KRAS Switch-II Pocket Mutation and Polyclonal Alterations Converging on RAS– MAPK Reactivation, Cancer Discov., 2021, 11, 1913–1922.

21. X. Wang, S. Allen, J. F. Blake, V. Bowcut, D. M. Briere, A. Calinisan, J. R. Dahlke, J. B. Fell, J. P. Fischer, R. J. Gunn, J. Hallin, J. Laguer, J. D. Lawson, J. Medwid, B. Newhouse, P. Nguyen, J. M. O’Leary, P. Olson, S. Pajk, L. Rahbaek, M. Rodriguez, C. R. Smith, T. P. Tang, N. C. Thomas, D. Vanderpool, G. P. Vigers, J. G. Christensen and M. A. Marx, Identification of MRTX1133, a Noncovalent, Potent, and Selective KRASG12DInhibitor, J. Med. Chem., 2022, 65, 3123–3133.

22. T. R. Gadek and D. A. Ockey, Inhibitors of protein-protein interactions, Expert Opin. Ther. Pat., 2002, 12, 393–400.

23. H. Wang, R. S. Dawber, P. Zhang, M. Walko, A. J. Wilson and X. Wang, Peptide-based inhibitors of protein–protein interactions: biophysical, structural and cellular consequences of introducing a constraint, Chem. Sci., 2021, 12, 5977–5993.

24. A. P. Silwal, R. Jahan, S. K. S. Thennakoon, S. P. Arya, R. M. Postema, E. C. V. Ark and X. Tan, A universal DNA aptamer as an efficient inhibitor against spike-protein/hACE2 interactions, Chem. Commun., 2022, 58, 8049–8052.

25. L. Otvos and J. D. Wade, Current challenges in peptide-based drug discovery, Front. Chem.,, DOI:10.3389/fchem.2014.00062.

26. L. Fernandez, R. H. Bustos, C. Zapata, J. Garcia, E. Jauregui and G. M. Ashraf, Immunogenicity in Protein and Peptide Based-Therapeutics: An Overview, Curr. Protein Pept. Sci., 2018, 19, 958–971.

27. V. Azzarito, K. Long, N. S. Murphy and A. J. Wilson, Inhibition of α-helix-mediated protein–protein interactions using designed molecules, Nat. Chem., 2013, 5, 161–173.

28. E. Lenci and A. Trabocchi, Peptidomimetic toolbox for drug discovery, Chem. Soc. Rev., 2020, 49, 3262–3277.

29. M. H. Caruthers, Chemical Synthesis of DNA and DNA Analogues, Acc. Chem. Res., 1991, 24, 278–284.

30. M. H. Caruthers, Robert Letsinger: The father of synthetic DNA chemistry, Proc. Natl. Acad. Sci., 2014, 111, 18098–18099.

31. E. M. LeProust, B. J. Peck, K. Spirin, H. B. McCuen, B. Moore, E. Namsaraev and M. H. Caruthers, Synthesis of high-quality libraries of long (150mer) oligonucleotides by a novel depurination controlled process, Nucleic Acids Res., 2010, 38, 2522–2540.

32. N. Appukutti and C. J. Serpell, High definition polyphosphoesters: between nucleic acids and plastics, Polym. Chem., 2018, 9, 2210–2226.

33. N. Appukutti, J. R. Jones and C. J. Serpell, Sequence isomerism in uniform polyphosphoesters programmes self-assembly and folding, Chem. Commun., 2020, 56, 5307–5310.

34. N. Appukutti, A. H. de Vries, P. G. Gudeangadi, B. R. Claringbold, M. D. Garrett, M. R. Reithofer and C. J. Serpell, Sequence-complementarity dependent co-assembly of phosphodiester-linked aromatic donor-acceptor trimers, Chem. Commun., 2022, 58, 12200–12203.

35. T. G. W. Edwardson, K. M. M. Carneiro, C. J. Serpell and H. F. Sleiman, An efficient and modular route to sequence-defined polymers appended to DNA, Angew. Chem. - Int. Ed., 2014, 53, 4567–4571.

36. P. Chidchob, T. G. W. Edwardson, C. J. Serpell and H. F. Sleiman, Synergy of Two Assembly Languages in DNA Nanostructures: Self-Assembly of Sequence-Defined Polymers on DNA Cages, J. Am. Chem. Soc., 2016, 138, 4416–4425.

37. C. F. Lin and L. E. Webb, Crystal Structures and Conformations of the Cyclic Dipeptides cyclo-(Glycyl-l-tyrosyl) and cyclo-(l-Seryl-l-tyrosyl) Monohydrate, J. Am. Chem. Soc., 1973, 95, 6803–6811.

38. K. Yoshizawa, K. Hirata, S. I. Ishiuchi, M. Fujii and A. Zehnacker, Do Stereochemical Effects Overcome a Charge-Induced Perturbation in Isolated Protonated Cyclo(Tyr-Tyr)?, J. Phys. Chem. A, 2022, 126, 6387–6394.

39. M. Al Kobaisi, S. V. Bhosale, K. Latham, A. M. Raynor and S. V. Bhosale, Functional Naphthalene Diimides: Synthesis, Properties, and Applications, Chem. Rev., 2016, 116, 11685–11796.

40. K. P. de Carvasal, N. Aissaoui, G. Vergoten, G. Bellot, J.-J. Vasseur, M. Smietana and F. Morvan, Folding of phosphodiester-linked donor-acceptor oligomers into supramolecular nanotubes in water, Chem. Commun., 2021, 57, 4130–4133.

41. K. S. Lam, M. Lebl and V. Krchnák, The ‘One-Bead-One-Compound’ Combinatorial Library Method., Chem. Rev., 1997, 97, 411–448.

42. A. Paul, M. Falsaperna, H. Lavender, M. D. Garrett and C. J. Serpell, Selection of Optimised Ligands by Fluorescence-Activated Bead Sorting, Chem. Sci., 2023, 14, 9517–9525.

43. W. Rapp, in Combinatorial Peptide and Nonpeptide Libraries: A Handbook, ed. G. Jung, VCH, 1996, pp. 425–464.

44. M. J. Hansen, W. A. Velema, M. M. Lerch, W. Szymanski and B. L. Feringa, Wavelength-selective cleavage of photoprotecting groups: Strategies and applications in dynamic systems, Chem. Soc. Rev., 2015, 44, 3358–3377.

45. X. Fang, W. Li, T. Gao, Q. U. Ain Zahra, Z. Luo and R. Pei, Rapid screening of aptamers for fluorescent targets by integrated digital PCR and flow cytometry, Talanta, 2022, 242, 123302.

46. L. A. Fraser, A. B. Kinghorn, M. S. L. Tang, Y. W. Cheung, B. Lim, S. Liang, R. M. Dirkzwager, J. A. Tanner and A. O. A. Miller, Oligonucleotide functionalised microbeads: Indispensable tools for high-throughput aptamer selection, Molecules, 2015, 20, 21298–21312.

47. Z. Surviladze, A. Waller, Y. Wu, E. Romero, B. S. Edwards, A. Wandinger-Ness and L. A. Sklar, Identification of a small GTPase inhibitor using a high-throughput flow cytometry bead-based multiplex assay, J. Biomol. Screen., 2010, 15, 10–20.

48. J. F. J. Todd, Recommendations for nomenclature and symbolism for mass spectroscopy, Int. J. Mass Spectrom. Ion Process., 1995, 142, 209–240.

49. P. J. Sample, K. W. Gaston, J. D. Alfonzo and P. A. Limbach, RoboOligo: Software for mass spectrometry data to support manual and de novo sequencing of post-transcriptionally modified ribonucleic acids, Nucleic Acids Res., 2015, 43, 1–13.

50. A. Cruz-Migoni, P. Canning, C. E. Quevedo, C. J. R. Bataille, N. Bery, A. Miller, A. J. Russell, S. E. V. Phillips, S. B. Carr and T. H. Rabbitts, Structure-based development of new RAS-effector inhibitors from a combination of active and inactive RAS-binding compounds, Proc. Natl. Acad. Sci. U. S. A., 2019, 116, 2545–2550.

51. G. A. Hobbs, C. J. Der and K. L. Rossman, RAS isoforms and mutations in cancer at a glance, J. Cell Sci., 129, 1287–1292.

52. S. R. Punekar, V. Velcheti, B. G. Neel and K.-K. Wong, The current state of the art and future trends in RAS-targeted cancer therapies, Nat. Rev. Clin. Oncol., 2022, 19, 637–655.

53. D. Kessler, M. Gmachl, A. Mantoulidis, L. J. Martin, A. Zoephel, M. Mayer, A. Gollner, D. Covini, S. Fischer, T. Gerstberger, T. Gmaschitz, C. Goodwin, P. Greb, D. Häring, W. Hela, J. Hoffmann, J. Karolyi-Oezguer, P. Knesl, S. Kornigg, M. Koegl, R. Kousek, L. Lamarre, F. Moser, S. Munico-Martinez, C. Peinsipp, J. Phan, J. Rinnenthal, J. Sai, C. Salamon, Y. Scherbantin, K. Schipany, R. Schnitzer, A. Schrenk, B. Sharps, G. Siszler, Q. Sun, A. Waterson, B. Wolkerstorfer, M. Zeeb, M. Pearson, S. W. Fesik and D. B. McConnell, Drugging an undruggable pocket on KRAS, Proc. Natl. Acad. Sci., 2019, 116, 15823–15829.

54. M. E. Welsch, A. Kaplan, J. M. Chambers, M. E. Stokes, P. H. Bos, A. Zask, Y. Zhang, M. Sanchez-Martin, M. A. Badgley, C. S. Huang, T. H. Tran, H. Akkiraju, L. M. Brown, R. Nandakumar, S. Cremers, W. S. Yang, L. Tong, K. P. Olive, A. Ferrando and B. R. Stockwell, Multivalent Small-Molecule Pan-RAS Inhibitors, Cell, 2017, 168, 878-889.e29.

55. R. Sahu, S. Yadav, S. Nath, J. Banjeree and A. R. Kapdi, DNA-Encoded Libraries Via Late-Stage Functionalization Strategies: A Review, Chem. Commun.,, DOI:10.1039/D3CC01075A.

56. L. Mannocci, M. Leimbacher, M. Wichert, J. Scheuermann and D. Neri, 20 years of DNA-encoded chemical libraries, Chem. Commun., 2011, 47, 12747–12753.

57. R. Stoltenburg, C. Reinemann and B. Strehlitz, SELEX-A (r)evolutionary method to generate high-affinity nucleic acid ligands, Biomol. Eng., 2007, 24, 381–403.

58. G. Caron, J. Kihlberg, G. Goetz, E. Ratkova, V. Poongavanam and G. Ermondi, Steering New Drug Discovery Campaigns: Permeability, Solubility, and Physicochemical Properties in the bRo5 Chemical Space, ACS Med. Chem. Lett., 2021, 12, 13–23.

59. Q. Laurent, R. Martinent, D. Moreau, N. Winssinger, N. Sakai and S. Matile, Oligonucleotide Phosphorothioates Enter Cells by Thiol-Mediated Uptake, Angew. Chem. Int. Ed., 2021, 60, 19102–19106.

60. J. A. Kulkarni, D. Witzigmann, S. B. Thomson, S. Chen, B. R. Leavitt, P. R. Cullis and R. van der Meel, The current landscape of nucleic acid therapeutics, Nat. Nanotechnol., 2021, 16, 630–643.

