## Supporting information for "Sequence-defined oligophosphoesters for selective inhibition of the KRAS G12D/RAF1 interaction"

### 1 General considerations

#### 1.1 Materials

TentaGel® M $NH_2$ Monosized Amino TentaGel Microspheres were purchased from Rapp Polymere. 3-amino-1-propanol, caesium carbonate, dichloromethane (dried over molecular sieves), ethylene glycol, pyridine (dried over molecular sieves), sodium hydroxide, N, N, N', N'-tetramethylethylenediamine (TEMED) and trichloroacetic acid were all purchased from Acros Organics. 4, 4'-dimethoxytrityl chloride and bisphenol A were purchased from Alfa Aesar. L-tyrosine was purchased from Apollo Scientific. Sheath fluid, Sphero™ Rainbow Calibration Beads (8-peak) 3.0 – 3.4 µM, and BD FACS™ Accudrop Beads were all purchased from BD Biosciences. 30% Acrylamide/Bis solution (29:1) was purchased from Bio-Rad Laboratories. 2-(2-chloroethoxy)ethanol, 3-bromoanisole, 4-amino-N, N-dimethylbenzylamine, 4-chloro-2-fluoro-3-methoxyphenyl boronic acid, 4-dimethylamino pyridine (DMAP), L-hydroxyproline, L-serine, L-threonine, N, N'-diisopropylcarbomide (DIC), N, N-diisopropylethylamine (DIPEA), palladium (II) acetate, Pd(dppf)$Cl_2$, rhodamine B, triethylamine (anhydrous), trityl chloride, and XPhos were all purchased from Fluorochem. 2-cyanoethyl N, N-diisopropyl chlorophosphoramidite was purchased from LGC Biosearch™ Technologies. 1, 5-dihydroxynaphthalene, naphthalenetetracarboxylic dianhydride, 5(6)-carboxyfluorescein, bovine serum albumin, dithiothreitol (DTT), glycine, methanol hyper grade for LC-MS LiChrosolv®, water for chromatography (LC-MS Grade) LiChrosolv®, triethylammonium bicarbonate (TEAB) and tween-20 were all purchased from Sigma-Aldrich. Acetonitrile, acetonitrile (dried over molecular sieves), ammonia solution (35%), acetic acid (glacial 99%), dichloromethane, dimethylformamide (DMF) diethyl ether, Dulbecco's phosphate buffered saline (PBS) solution, ethanol, ethyl acetate, glycerol, hexane, magnesium chloride, magnesium sulphate, methanol, n-butanol, ninhydrin, phenol, potassium cyanide (KCN), pyridine, sodium carbonate, sodium hydrogen carbonate, trifluoroacetic acid, tris, sodium chloride, sodium dodecyl sulfate (SDS), ammonium persulfate, Nunc™ 96-well plates (clear), PageRuler™ Prestained Protein Ruler, potassium carbonate, Pierce™ Silver Stain Kit, Pierce™ C18 Spin Tips and Columns, streptavidin (fluorescein conjugate), and streptavidin (Rhodamine Red™-X conjugate) were all purchased from Thermo Fisher Scientific. All NMR samples were run in either chloroform ($CDCl_3$) or dimethylsulfoxide (DMSO-$d_6$), purchased from Goss Scientific. Mass spectrometry columns nanoEase™ m/z symmetry C18, 180 µM x 20 mm column (Trap Column) and nanoEase™ m/z HSS C18T3, 100 Å, 1.8 µM, 75 µM x 150 mm column were purchased from Waters. Universal Unylinker Support Beads were purchased from ChemGenes Corporation. Cap Mix A (THF/Pyridine/Acetic Anhydride 8:1:1), Cap Mix B (10% methylimidazole in THF), 2-cyanoethylN, N-diisopropylchlorophosphoramidite, ETT Activator (0.25 M) (5-ethylthio-1H-tetrazole in Acetonitrile), Oxidiser (0.02 M Iodine in 20% Pyridine), PC Spacer ([4-(4, 4'-Dimethoxytrityloxy) butyramidomethyl)-1-(2-nitrophenyl)-ethyl]-2-cyanoethyl- (N, N-diisopropyl)-phosphoramidite), monomer **2** (Spacer CE-Phosphoramidite C12) and monomer **3** (Spacer CE-Phosphoramidite 18) were all purchased from LGC Biosearch™ Technologies.

The custom made 7 base DNA standard, leucine enkephalin for LC-MS, LiChrosolv® methanol (hypergrade for LC-MS), LiChrosolv® water (hypergrade for LC-MS), and triethylammonium bicarbonate (TEAB) were all purchased from Sigma Aldrich. Acetonitrile, Pierce™ C18 Spin tips and columns, and trifluoroacetic acid (TFA) were all purchased from Thermo Fisher Scientific. The trap column (nanoEase™ *m/z* symmetry C18, 180 µM x 20 mm column) and UHPLC column (nanoEase™ *m/z* HSS C18T3, 100 Å, 1.8 µM, 75 µM x 150 mm column) were purchased from Waters.

#### 1.2 Buffers

**KRAS protein buffer:** 10 mM Tris (pH 7.74), 50 mM NaCl, 2 mM $MgCl_2$ made up to 100 mL in milliQ water.

**RAF protein buffer:** 10 mM Tris (pH 7.74), 50 mM NaCl made up to 100 mL in milliQ water.

**Tris-glycine running buffer:** 25 mM Tris (pH 7.42), 192 mM Glycine, made up to 1 L in milliQ water.

**Tris-glycine sample buffer (4x):** 62.5 mM Tris (pH 6.85), 25% glycerol (v/v), 1% bromophenol blue, made up to 10 mL in milliQ water.

**Tris-glycine SDS buffer:** 25 mM Tris base (pH 7.42), 200 mM Glycine, 0.1% SDS (w/v) made up to 1 L in milliQ water.

**Tris-glycine SDS sample buffer (5x):** 10% w/v SDS, 10 mM DTT, 20% v/v glycerol, 0.2 M Tris (pH 6.82), 0.05% bromophenol blue.

**Coating buffer:** 10 mM $Na_2CO_3$, 30 mM $NaHCO_3$ (pH 9) made up to 1 L in milliQ water.

**Blocking buffer:** 20 mM Tris (pH 7.6), 150 mM NaCl, 2 mM $MgCl_2$, 1 mM DTT, 0.5% BSA (w/v), 0.05% Tween-20 (v/v) made up to 1 L in milliQ water.

**PBS Buffer:** 20 mM Tris (pH 7.6), 150 mM NaCl, 2 mM $MgCl_2$, 1 mM DTT, 0.05% Tween (v/v), made up to 1 L in PBS.

**MS Buffer A:** 8 mM TEAB in LC-MS Grade LiChrosolv® water.

**MS Buffer B:** 8 mM TEAB in LC-MS Grade LiChrosolv® water and LC-MS Grade LiChrosolv® methanol (1:1).

#### 1.3 Instrumentation

The oligomers were synthesised on an Expedite™ 8909 Nucleic Acid Synthesiser system provided by Biolytic. Sequences for the oligomer library were created using Validate XP software (5.4.15). All phosphoramidites were dissolved in dry dichloromethane (10 mL per 0.5 g of sample) and oligomers were synthesised at 1 µM.

Automated column purification was carried out using a Biotage Isolera One system. Specific column and solvent systems are detailed for each compound. Columns for purification were purchased from Biotage.

NMR spectra were obtained on a Bruker AVII 400 MHz spectrometer, 400 MHz for $^{1}H$ NMR spectra and 100 MHz for $^{13}C$ spectra and calibrated to the centre of the set deuterated solvent peak. Chemical shifts were reported in parts per million (ppm) and J coupling values were reported in hertz (Hz). All NMR data was processed using ACD-NMR processor software (Academic Version 12.01).

Low resolution mass spectrometry data was obtained using a Thermo MSQPlus instrument fitted with a Zorbax SB-C18 5 µm 3.0 x 150 mm column. The mobile phases used were $H_2O$ + 0.1% formic acid and methanol + 0.1% formic acid in positive mode or $H_2O$ + 0.1% ammonia and acetonitrile + 0.1% ammonia in negative mode. The gradient in both positive and negative mode was as shown in **Table S 1**. The flow rate was 1 mL/min, and the injection volume was 10 µL. Data was analysed using Chromeleon™ Chromatography Data System (CDS) Software.

**Table S 1.** LCMS gradients.

| % Solvent A | % Solvent B | Time (minutes) |
|---|---|---|
| **70** | 30 | 0.0 |
| **70** | 30 | 2.0 |
| **5** | 95 | 7.0 |
| **5** | 95 | 10.0 |
| **70** | 30 | 10.2 |
| **70** | 30 | 13.0 (End) |

A Bruker micrOTOF-Q LCMS system was used to obtain high-resolution mass spectrometry data for small molecules. Samples were dissolved in HPLC-grade methanol and injected using direct injection mode with a mobile phase system of 50:50 methanol and $H_2O$. Data was processed using Bruker Compass Hystar software.

Flow cytometry data was collected on a BD FACSJazz™ Cell Sorter by Becton Dickson. Data collected from flow cytometry was analysed using BD™ FACS Sortware software.

Fluorescent gels were imaged using GeneSys Image Capture Software (version 1.5.2.0) with either fluorescein (Automatic Exposure Blue LED Module filter 525 nm) or rhodamine (Automatic Exposure Green LED Module filter 605 nm) filters.

All 96 well plates were analysed using the ClarioStar Plus plate reader, analysing the emission spectrum of fluorescein (500 – 600 nm) and GFP (490 – 560 nm). Data analysis (area under the curve calculations) was performed on GraphPad Prism (version 9.5.1).

A NanoDrop One UV-Vis Spectrophotometer (Thermo Fisher Scientific) was used to collect absorption data at 500 nm.

### 2 Chemical synthesis and characterisation

#### *2.1 Synthesis of phosphoramidite monomers*

**BPA-trityl**

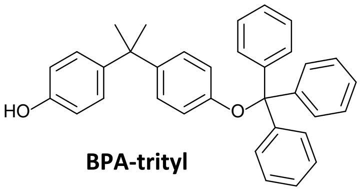

**BPA-trityl**

Bisphenol A (0.500 g, 2.19 mmol) and dimethylaminopyridine (0.011 g, 0.09 mmol) were dissolved in dichloromethane (45 mL). Triethylamine (305 µL, 2.19 mmol) was added dropwise, after which the reaction was cooled to 0 °C. Once cool, trityl chloride (0.488 g, 1.75 mmol) dissolved in anhydrous dichloromethane (5 mL) was added dropwise. The reaction was stirred at 0 °C for a further 90 minutes then left to return to room temperature. The reaction was stirred for 24 hours at room temperature under nitrogen. After stirring, the mixture was washed with water (50 mL). The organic layer was separated, dried over magnesium sulphate, after which the solvent was removed *in vacuo*. The crude residue was purified with flash chromatography using the Biotage Isolera One, running a hexane and ethyl acetate gradient. The final product (**BPA-trityl**) was a clear oil (0.326 g, 39.5%).

$^{1}$H NMR (400 MHz, DMSO-d$_6$) δ (ppm): 9.17 (1H, s, O**H**), 7.40 (5H, m, trityl), 7.31 (6H, m, trityl), 7.23 (4H, m, trityl), 6.84 (4H, m, BPA Ar-**H**), 6.60 (2H, m, BPA Ar-**H**), 6.53 (2H, m, BPA Ar-**H**), 1.44 (6H, s, C**H$_3$**)

$^{13}$C NMR (100 MHz, DMSO-d$_6$) δ (ppm): 155.48, 155.42, 144.30, 144.03, 141.43, 128.88, 128.31, 127.73, 127.64, 126.93, 120.24, 115.01, 89.96, 62.97, 52.43, 31.35, 31.09, 7.65

MS: m/z calculated ($C_{34}H_{30}O_2$): 469.2251 [M-H]$^-$ Found: 469.2295 [M-H]$^-$

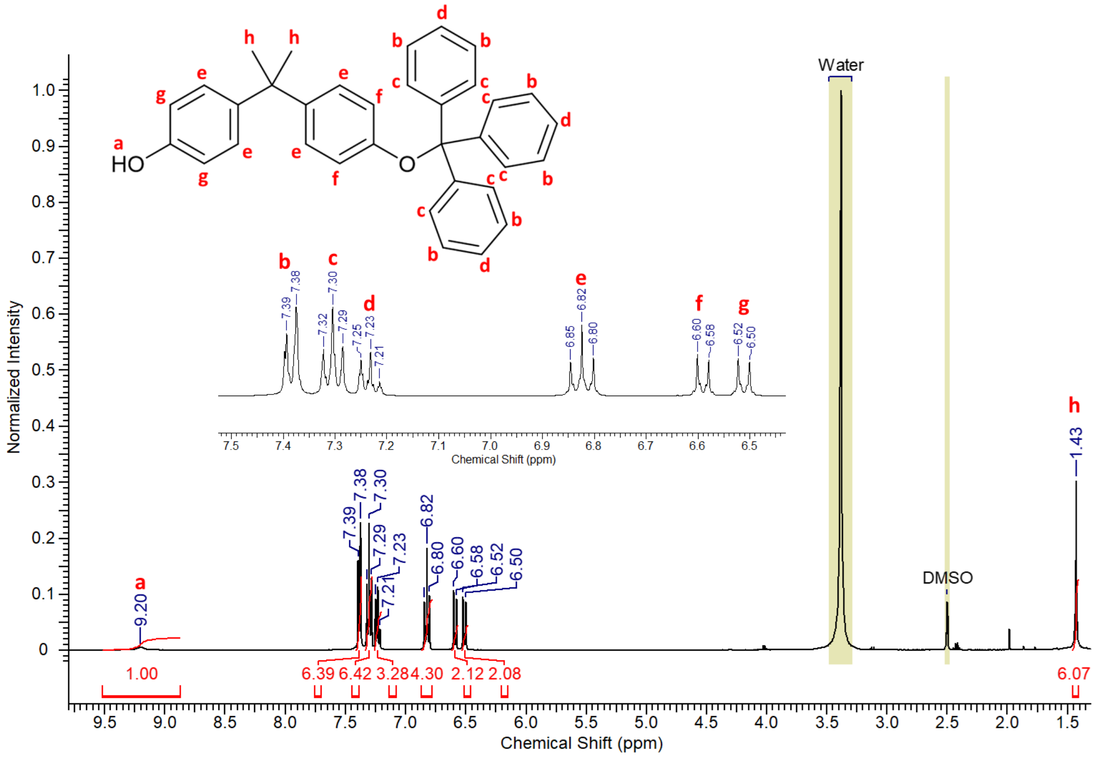

**Figure S 1.** $^{1}$H NMR spectrum of **BPA-trityl**.

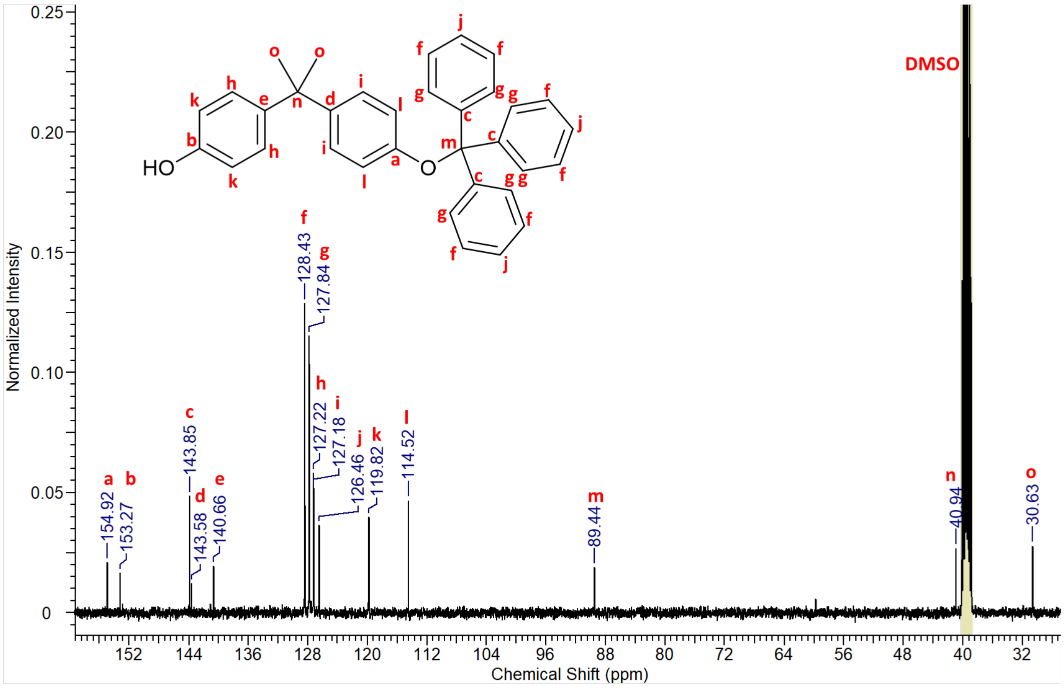

**Figure S 2.** $^{31}$C NMR spectrum of **BPA-trityl**.

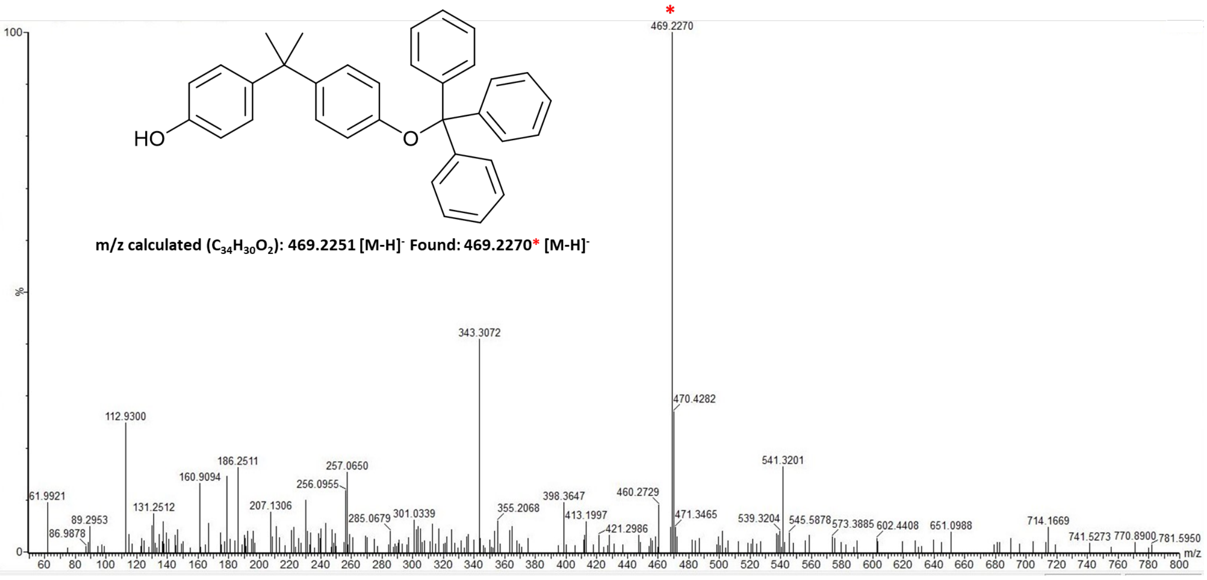

**Figure S 3.** Mass spectrum of **BPA-trityl**.

**BPA phosphoramidite monomer**

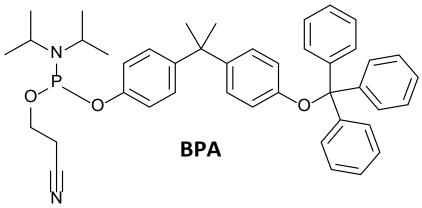

**BPA-trityl** (0.200 g, 0.425 mmol), dimethylaminopyridine (0.005 g, 0.043 mmol), and diisopropylethylamine (370 µL, 2.125 mmol) were all dissolved in anhydrous dichloromethane (25 mL) and stirred under nitrogen. 2-cyanoethyl N, N-diisopropylchlorophosphoramidite (284 µL, 1.275 mmol) was then added, and the reaction mixture was stirred for 2 hours under nitrogen. The dichloromethane was removed *in vacuo*, leaving a yellow oil (0.172 g, 60.3%). $^{31}$P NMR was run on the product, to prove the addition of the phosphoramidite group. No other analysis was run on it because of the air sensitive nature of the product.

$^{31}$P NMR (160 MHz, DMSO-$d_6$ capillary in DCM) δ (ppm): 148.75, 145.62, 138.55, 138.32

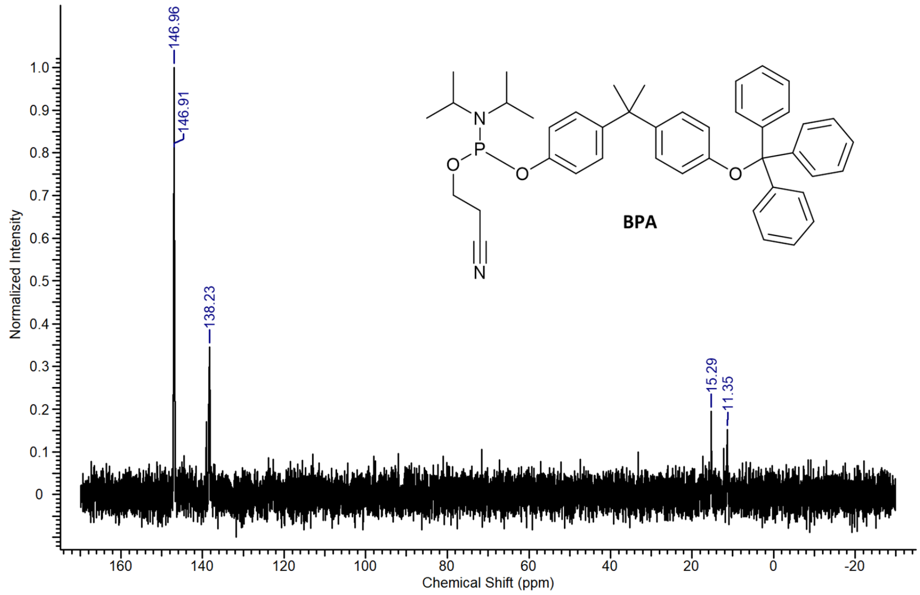

**Figure S 4.** $^{31}$P NMR spectrum of **BPA**.

**cSS-diol**

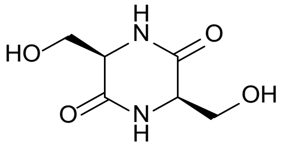

**cSS-diol**

L-serine (5 g, 0.047 mol) was added to ethylene glycol (20 mL, 0.358 mol) and stirred under an atmosphere of nitrogen. After stirring the mixture was refluxed (220 °C) for 24 hours under nitrogen. The reaction mixture was left to cool to room temperature before being cooled further to 4 °C for 24 hours, causing a white solid to precipitate. The solid was filtered, washed with ethanol (10 mL) and dried, giving the final product of a beige powder (**cSS-diol**) (2.316 g, 28.0%).

$^{1}$H NMR (400 MHz, DMSO-$d_6$) δ (ppm): 7.88 (2H, s, N**H**), 4.97 (2H, t, J = 5.3 Hz, O**H**), 3.72 (4H, m, C**H$_2$**-OH), 3.57 (2H, m, C**H**-NH)

$^{13}C$ NMR (100 MHz, DMSO-$d_6$) δ (ppm): 172.73, 62.95, 60.59, 56.80

MS: m/z calculated ($C_6H_{10}N_2O_4$): 197.06 $[M+Na]^+$ Found: 197.0 $[M + Na]^+$

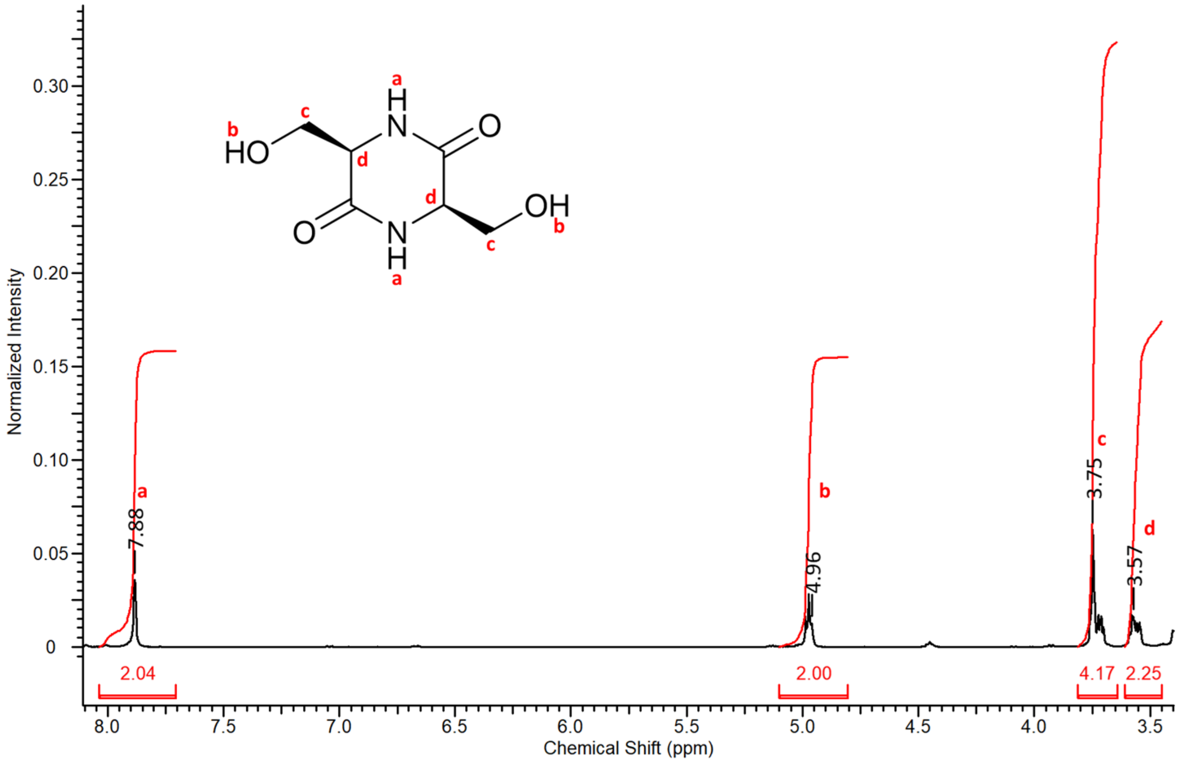

**Figure S 5.** $^1H$ NMR spectrum of **cSS-diol**.

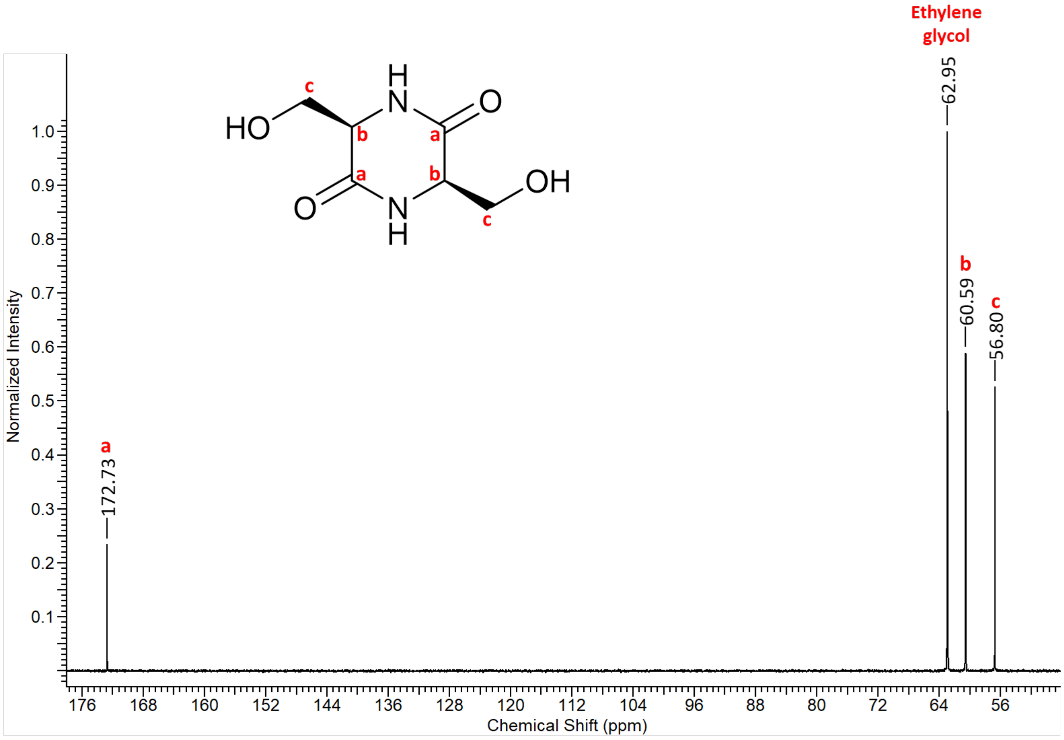

**Figure S 6.** $^{13}C$ NMR spectrum of **cSS-diol**.

**cSS-DMT**

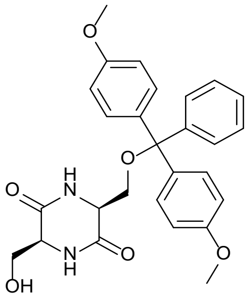

**cSS-DMT**

**cSS-diol** (1 g, 5.74 mmol) was dissolved in pyridine (45 mL), and dimethylaminopyridine (0.071 g, 0.58 mmol) and diisopropylethylamine (1.20 mL, 6.89 mmol) was added after and the solution was stirred under nitrogen. Dimethoxytrityl chloride (972 mg, 2.87 mmol) dissolved in pyridine (5 mL) was added dropwise, and the reaction was left stirring overnight under nitrogen. After stirring, the pyridine was removed *in vacuo*. The crude residue was purified through flash chromatography, using a 10 g SNAP KP-Sil column (Biotage), running a gradient of dichloromethane (0.1% TEA) and methanol (0.1% TEA) (0% to 100% methanol (0.1% TEA)). The final product was a yellow oil (**cSS-DMT**) (0.602 g, 22.0%).
$^1$H NMR (400 MHz, $CDCl_3$) δ (ppm): 7.36 (2H, d, J = 7.2 Hz, N**H**), 7.25 (4H, d, J = 9.0 Hz, trityl Ar-**H**), 7.21 (4H, m, trityl Ar-**H**), 7.10 (1H, d, J = 9.0 Hz, trityl Ar-**H**), 6.75 (4H, d, J = 8.9 Hz, trityl Ar-**H**), 3.72 (2H, s, CH-C**H$_2$**-ODMT), 3.71 (6H, s, C**H$_3$**), 3.41 (1H, s, C**H**-$CH_2$-ODMT), 2.97 (3H, s, C**H$_2$**-C**H**-OH)
$^{13}$C NMR (100 MHz, $CDCl_3$) δ (ppm): 157.94, 136.45, 130.29, 129.31, 128.26, 113.63, 55.25, 55.15, 29.73
MS: m/z calculated ($C_{27}H_{28}N_2O_6$): 475.1953 [M-H]$^-$. Found: 475.2891 [M-H]$^-$

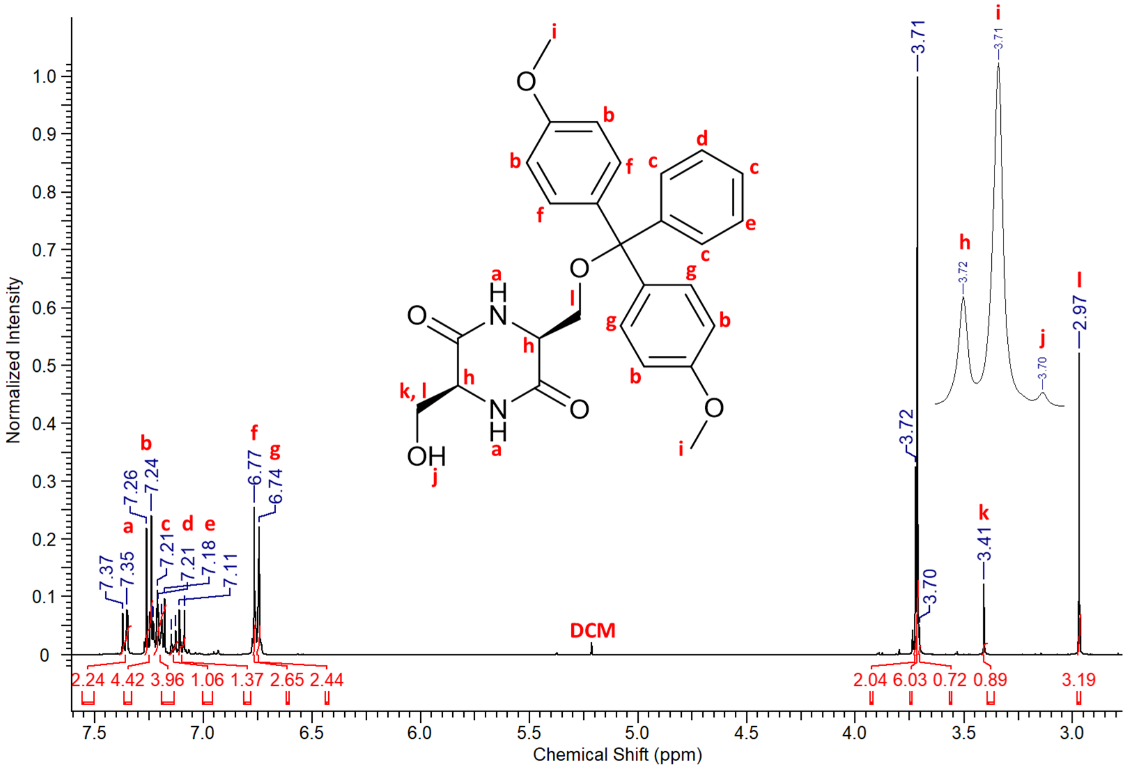

**Figure S 7**. $^1$H NMR spectrum of **cSS-DMT**.

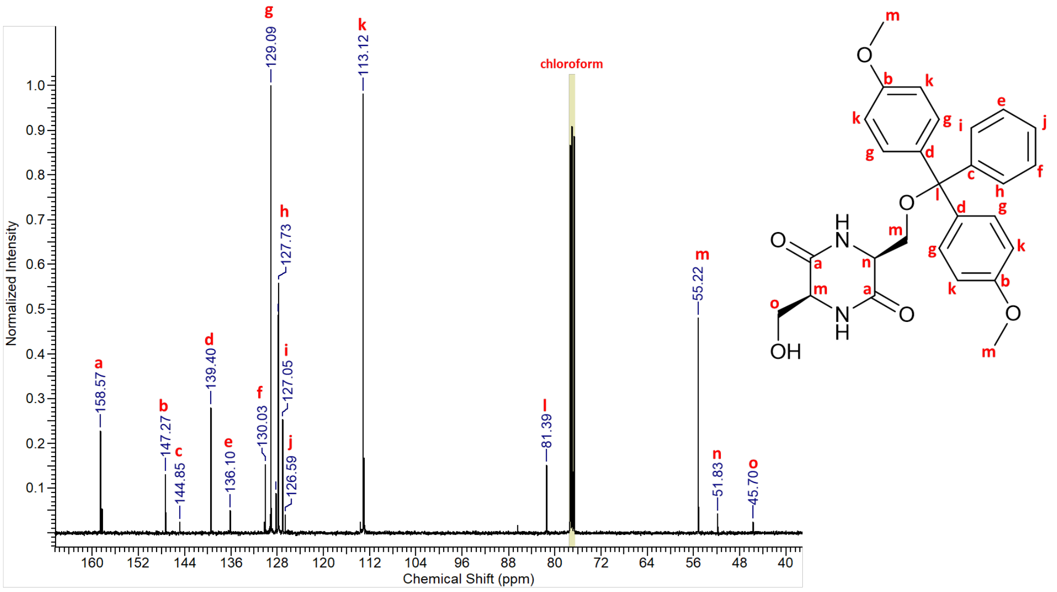

**Figure S 8.** $^{13}C$ NMR spectrum of **cSS-DMT**.

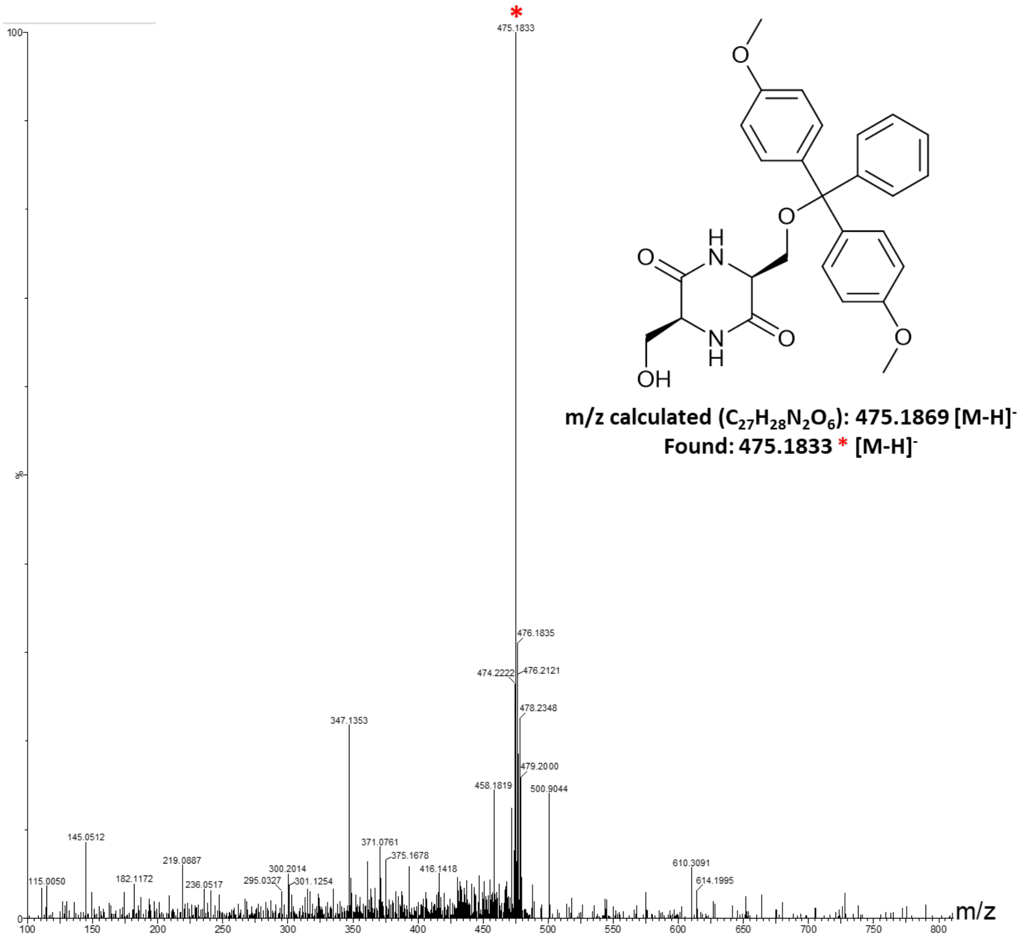

**Figure S 9.** Mass spectrum of **cSS-DMT**.

**cSS phosphoramidite monomer**

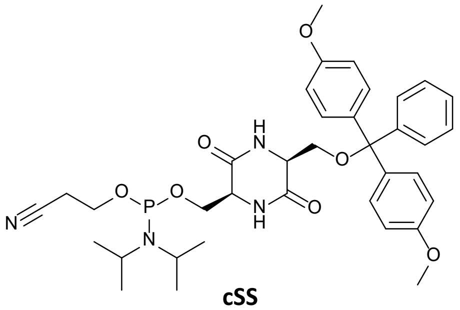

**cSS**

**cSS-DMT** (0.345 g, 0.724 mmol), dimethylaminopyridine (0.009 g, 0.072 mmol), and diisopropylethylamine (630 µL, 3.620 mmol), were all dissolved in anhydrous dichloromethane (25 mL) and stirred under nitrogen. 2-cyanoethyl N, N-diisopropyl chlorophosphoramidite (484 µL, 2.171 mmol) was added after, and the reaction was left to stir for 2 hours at room temperature. After stirring the dichloromethane was removed *in vacuo*. The reaction mixture was a brown solid (0.120 g, 88.6%) $^{31}P$ NMR was run on the product, to prove the addition of the phosphoramidite group. No other analysis was run on it because of the air sensitive nature of the product.
$^{31}P$ NMR (160 MHz, DMSO-$d_6$ capillary in DCM) δ (ppm): 148.04, 139.07

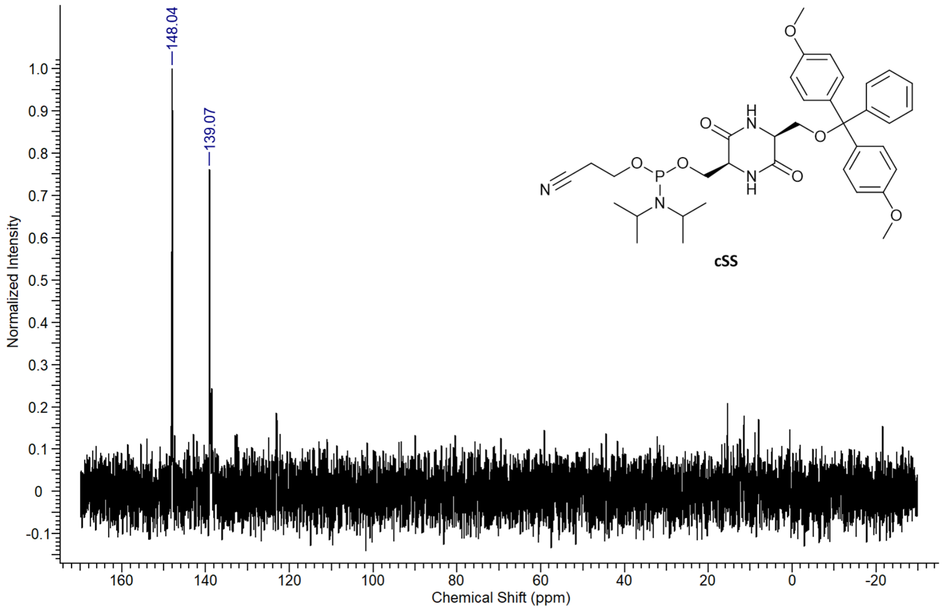

**Figure S 10**. $^{31}P$ NMR spectrum of **4**.

**cYY-diol**

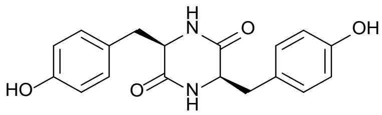

**cYY**

L-tyrosine (5 g, 0.028 mol) was added to ethylene glycol (20 mL, 0.358 mol) and stirred under an atmosphere of nitrogen for 15 minutes. After stirring the mixture was refluxed (220 °C) for 24 hours under nitrogen. The reaction mixture was cooled to room temperature, before pouring into a minimal amount of ice, causing an orange solid to precipitate. The solid was filtered, washed with cold water (10 mL) and dried, giving the final product of light orange powder (**cYY-diol**) (7.142 g, 79.3%).

$^{1}$H NMR (400 MHz, DMSO-$d_6$) δ (ppm): 9.27 (2H, s, O**H**), 7.92 (1H, s, N**H**), 7.78 (1H, d, J = 2.3 Hz, N**H**), 6.91 (2H, d, J = 8.5 Hz, $CH_2$-Ar-**H**), 6.84 (2H, d, J = 8.5 Hz, $CH_2$-Ar-**H**), 6.68 (2H, d, J = 8.4 Hz, HO-Ar-**H**), 6.62 (2H, d, J = 8.5 Hz, HO-Ar-**H**), 4.50 (1H, s, CO-C**H**-NH), 3.86 (1H, t, J = 6.5 Hz, CO-C**H**-NH), 2.90 (1H, dd, J = 13.7, 3.4 Hz, C**H₂**-Ar), 2.60 (1H, dd, J = 13.8, 4.7 Hz, C**H₂**-Ar), 2.54 (1H, d, J = 4.5 Hz, C**H₂**-Ar), 2.10 (1H, dd, J = 13.6, 6.6 Hz, C**H₂**-Ar)

$^{13}$C NMR (100 MHz, DMSO-$d_6$) δ (ppm): 167.65, 166.88, 156.50, 131.55, 131.25, 126.94, 126.23, 115.53, 115.28, 56.19, 55.27, 37.44

MS: m/z calculated ($C_{18}H_{18}N_2O_4$): 327.13 [M+H]$^+$, 349.12 [M+Na]$^+$ Found: 327.1 [M+H]$^+$, 349.1 [M+Na]$^+$

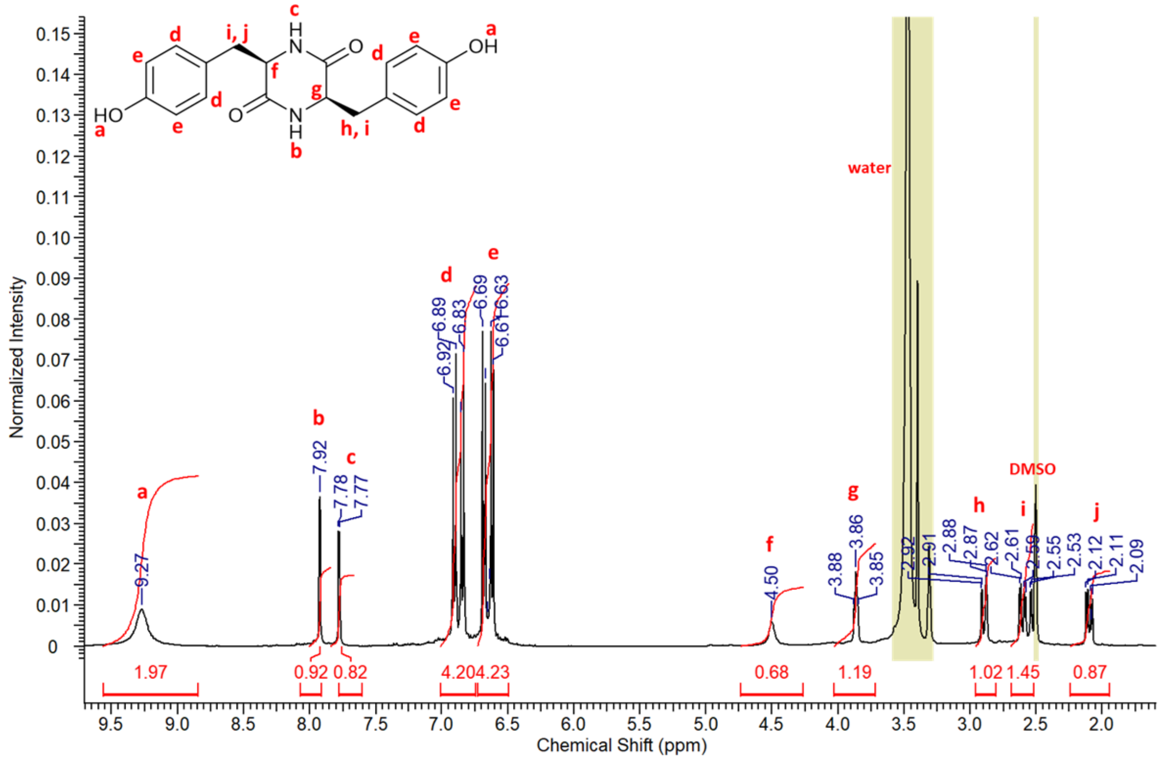

**Figure S 11.** $^{1}$H NMR spectrum of **cYY-diol**.

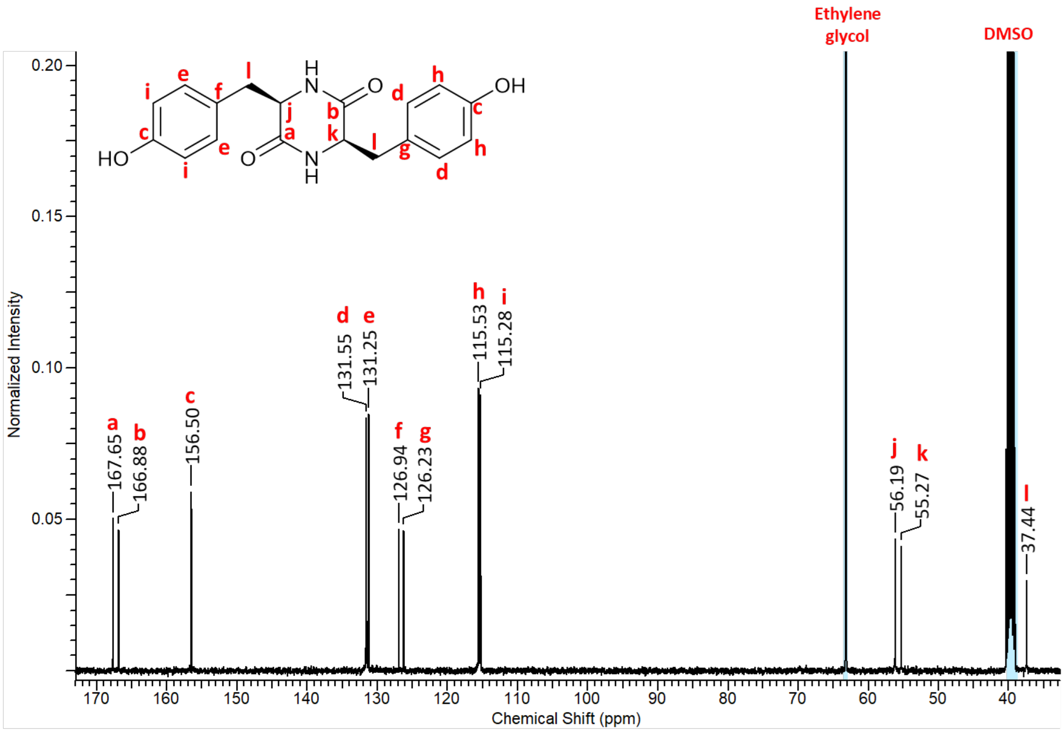

**Figure S 12.** $^{13}C$ NMR spectrum of **cYY-diol**.

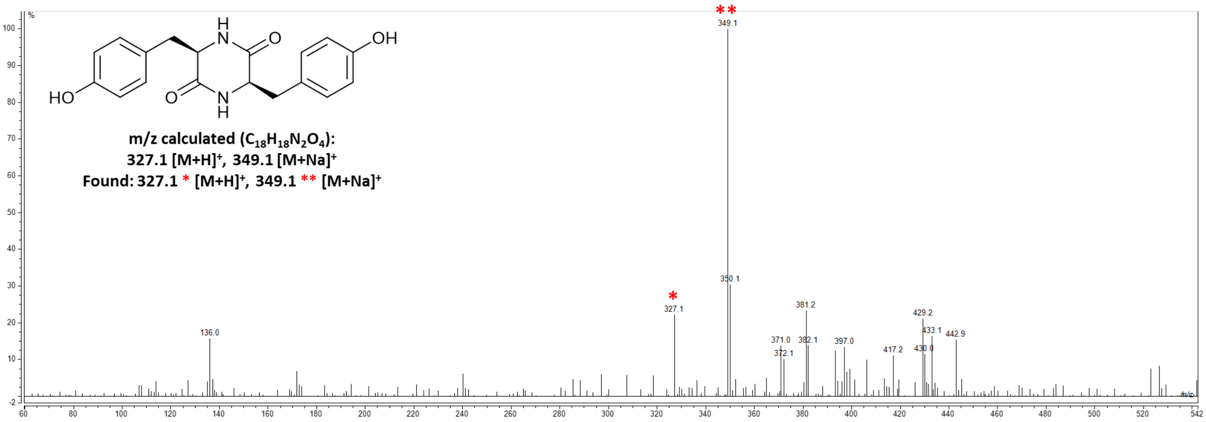

**Figure S 13.** Mass spectrum of **cYY-diol**.

#### cYY-tritYl

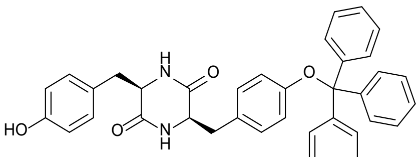

**cYY-trityl**

**cYY-diol** (0.5 g, 1.53 mmol) was dissolved in a minimal amount of anhydrous pyridine (100 mL). Dimethylaminopyridine (0.033 g, 0.27 mmol) and diisopropylethylamine (324 µL, 1.84 mmol) were then added and the solution was then stirred under nitrogen for 30 minutes. The reaction was then cooled to 0 °C before trityl chloride (0.427 g, 1.53 mmol) dissolved in anhydrous pyridine (5 mL) was added dropwise, after which the reaction was left at 0 °C for one hour. The reaction was left to come back up to room temperature before stirring overnight under nitrogen. After stirring, the solvents were removed *in vacuo*. The crude residue was purified using flash chromatography with a 10 g SNAP Ultra column (Biotage), running a gradient of dichloromethane (0.1% TEA) and methanol (0.1% TEA) (0% to 100% methanol (0.1% TEA)). The final product (**cYY-trityl**) was a white solid (0.269 g, 30.9%).

$^{1}$H NMR (400 MHz, DMSO-$d_6$) δ (ppm): 7.42 (9H, d, J = 7.3 Hz, trityl Ar-**H**), 7.34 (9H, t, J = 7.6 Hz, Ar-**H**), 7.26 (5H, m, Ar-**H**), 4.73 (2H, t, J = 5.7 Hz, $CH_2$-C**H**), 3.57 (4H, q, J = 5.5 Hz, -C**H₂**-), 2.98 (3H, t, J = 5.5 Hz, $CH_2$-C**H** and O**H**)

$^{13}$C NMR (100 MHz, DMSO-$d_6$) δ (ppm): 167.64, 166.87, 156.50, 148.14, 131.57, 131.25, 128.18, 128.01, 127.15, 126.91, 126.18, 115.52, 115.27, 81.03, 63.17, 56.18, 55.24, 37.40

MS: m/z calculated ($C_{37}H_{32}N_2O_4$): 569.2368 $[M+H]^+$ Found: 569.2782 $[M+H]^+$

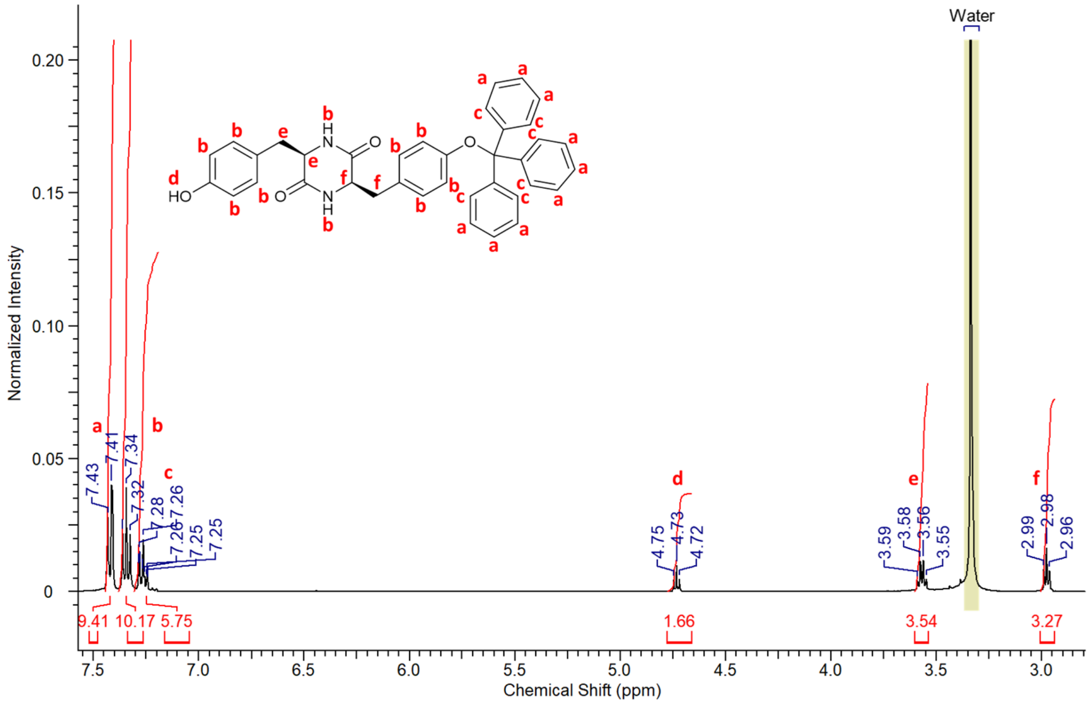

**Figure S 14**. $^{1}$H NMR spectrum of **cYY-DMT**.

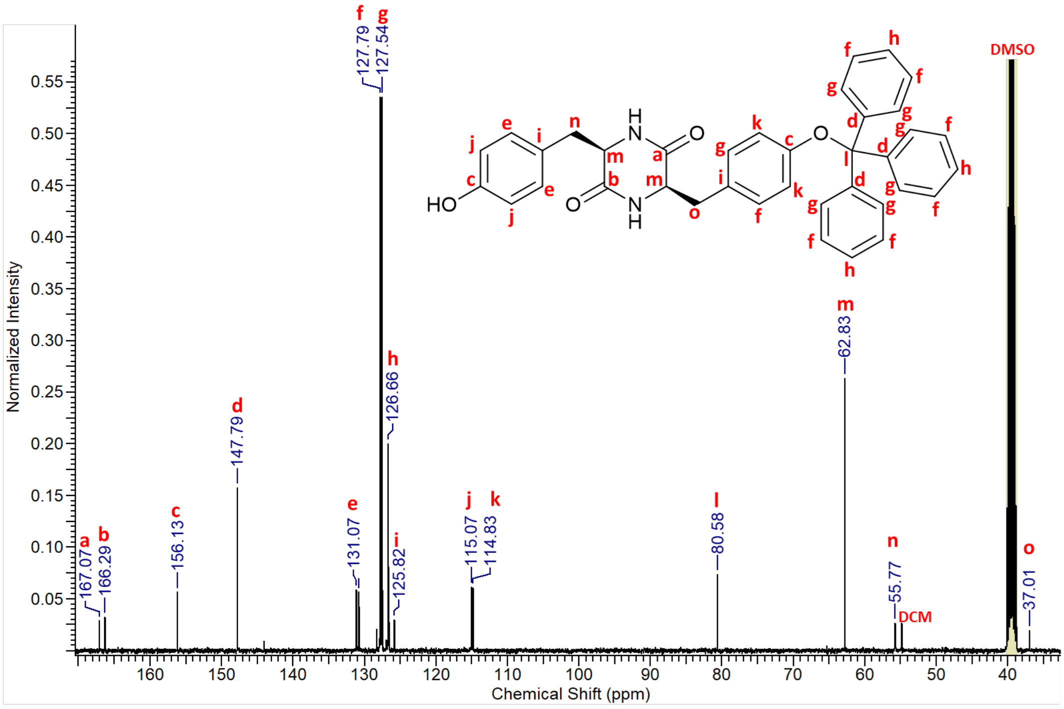

**Figure S 15.** $^{13}$C NMR spectrum of **cYY-trityl**.

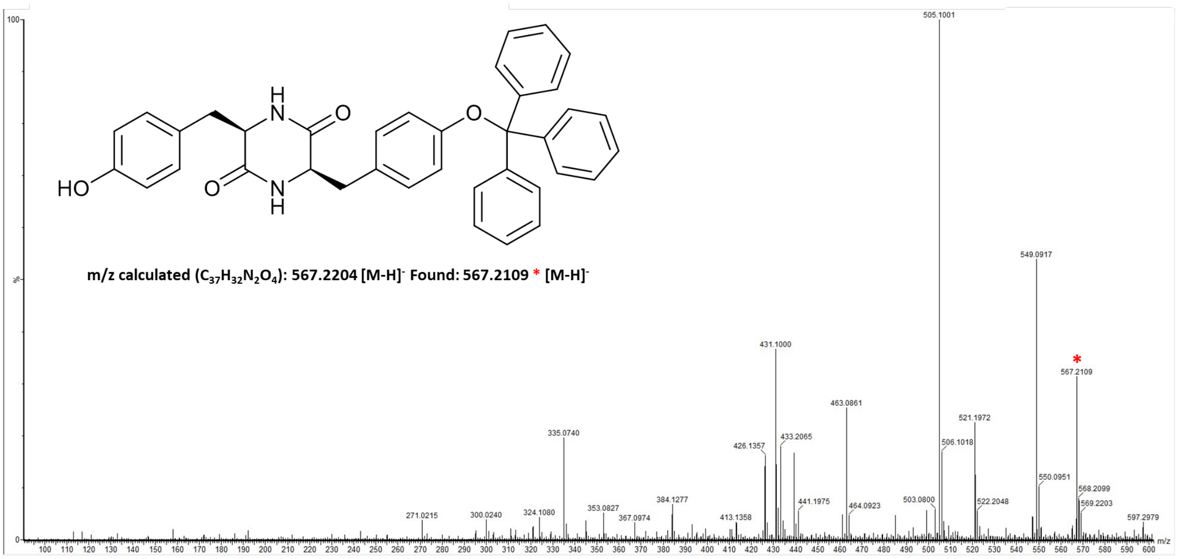

**Figure S 16.** Mass spectrum of **cYY-trityl**.

**cYY phosphoramidite monomer**

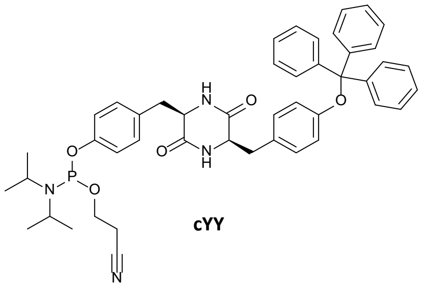

cYY

**cYY-trityl** (0.225 g, 0.396 mmol), dimethylaminopyridine (0.005 g, 0.040 mmol), and diisopropylethylamine (344 µL, 1.978 mmol), were all dissolved in anhydrous dichloromethane (25 mL) and stirred under nitrogen. 2-cyanoethyl N, N-diisopropyl chlorophosphoramidite (265 µL, 1.187 mmol) was added after, and the reaction was left to stir for 2 hours at room temperature. After stirring the dichloromethane was removed *in vacuo*. The reaction mixture as a yellow oil (0.189 g, 62.1%). $^{31}$P NMR was run on the product, to prove the addition of the phosphoramidite group. No other analysis was run on it because of the air sensitive nature of the product.

$^{31}$P NMR (160 MHz, DMSO-d$_6$ capillary in DCM) δ (ppm): 147.94, 138.57, 138.34

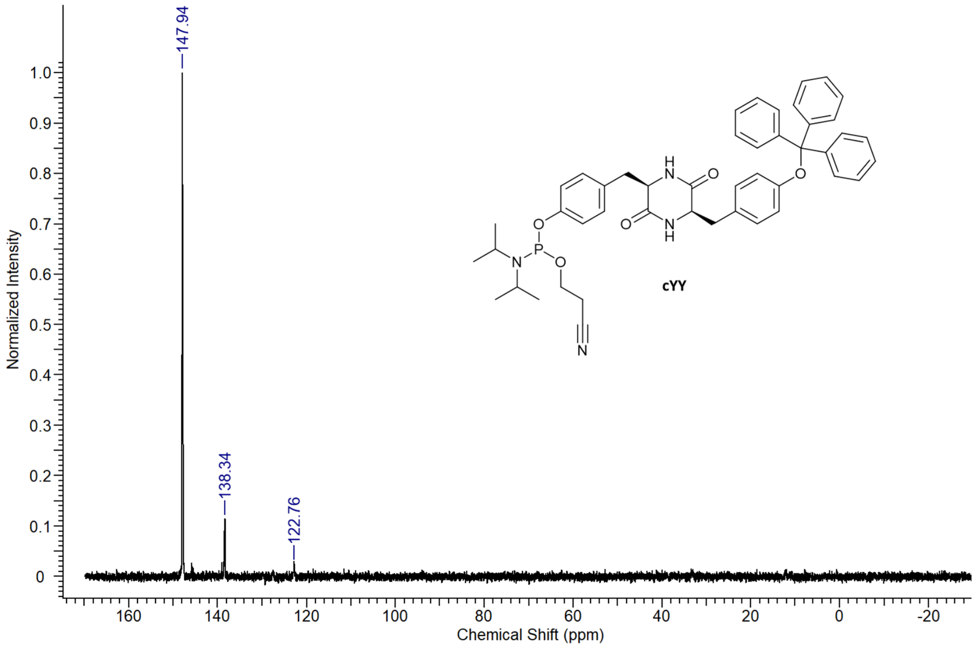

**Figure S 17.** $^{31}$P NMR spectrum of **cYY**.

**NDI-diol**

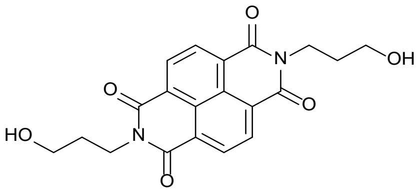

**NDI-diol**

Naphthalenetetracarboxylic dianhydride (0.5 g, 1.86 mmol) was added to a 30 mL microwave tube, followed by 3-aminopropan-1-ol (282 µL, 3.69 mmol) and water (20 mL). This mixture was heated using microwave irradiation for 30 minutes at 200 °C. After this the mixture was cooled to room temperature and filtered. The solid was washed with water (2 x 20 mL) and diethyl ether (2 x 20 mL), producing a pink powder (**NDI-diol**) (0.581 g, 81.6%).

$^{1}H$ NMR (400 MHz, DMSO-$d_6$) δ (ppm): 8.45 (4H, s, Ar-**H**), 4.57 (2H, t, J = 5.1 Hz, O**H**), 4.05 (4H, t, J = 7.3 Hz, N-C**H₂**), 3.52 (4H, q, J = 6.0 Hz, C**H₂**-OH), 1.80 (4H, quin, J = 7.1 Hz, $CH_2$-C**H₂**-$CH_2$)

$^{13}C$ NMR (100 MHz, DMSO-$d_6$) δ (ppm): 162.72, 130.67, 126.39, 59.44, 38.55, 31.18

MS: m/z calculated ($C_{20}H_{18}N_2O_6$): 383.12 $[M+H]^+$, 405.11 $[M+Na]^+$ Found: 383.0 $[M+H]^+$, 405.5 $[M+Na]^+$

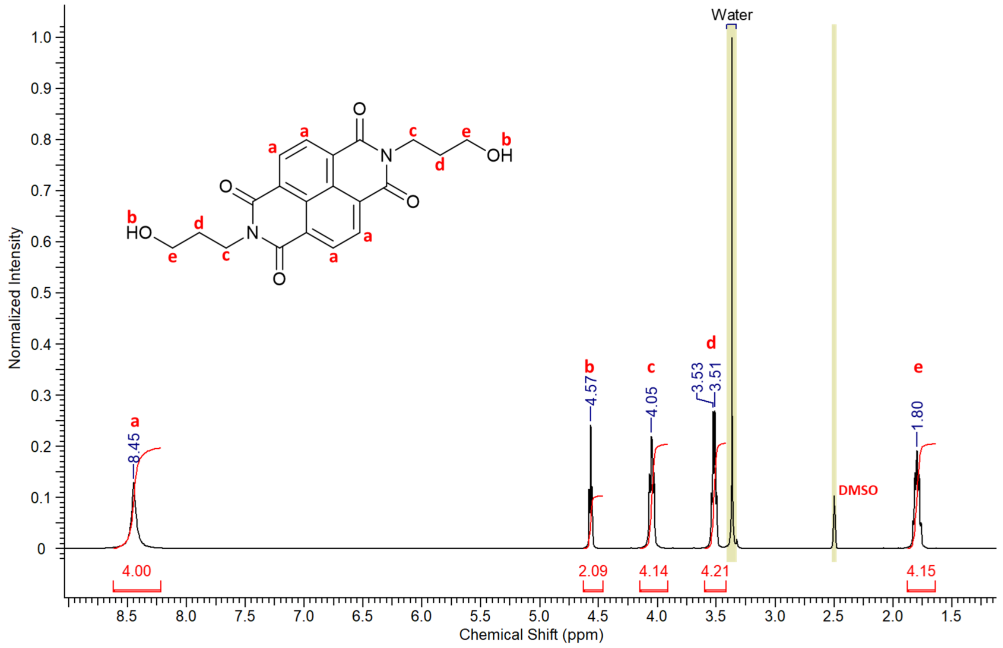

**Figure S 18.** $^{1}H$ NMR spectrum of **NDI-diol**.

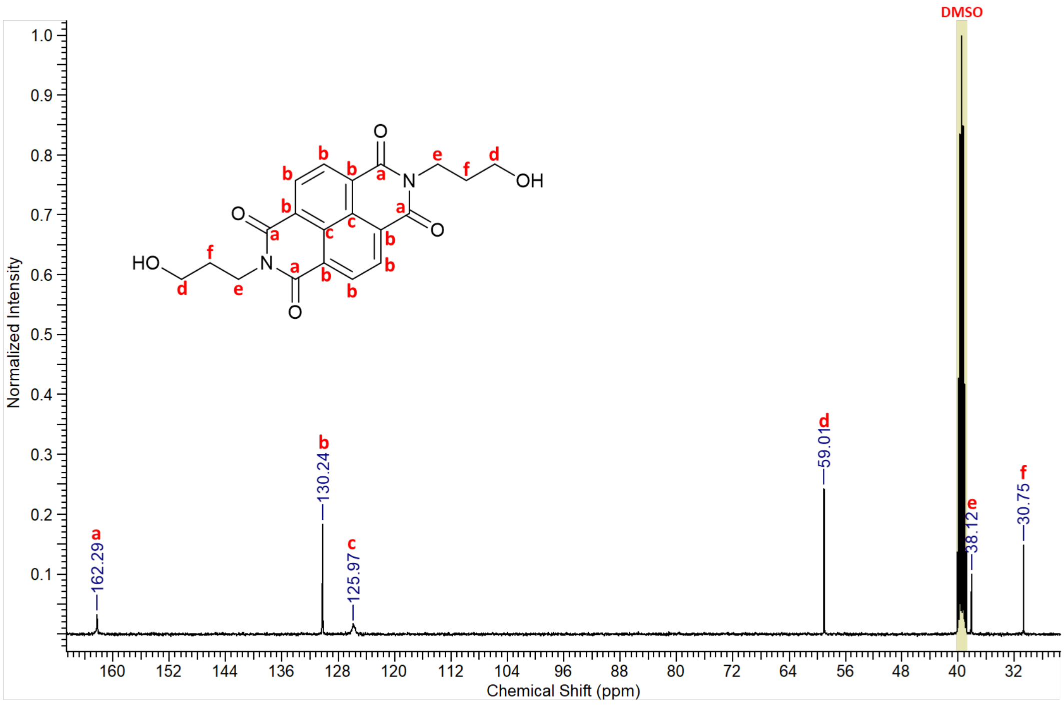

**Figure S 19.** $^{13}C$ NMR spectrum of **NDI-diol**.

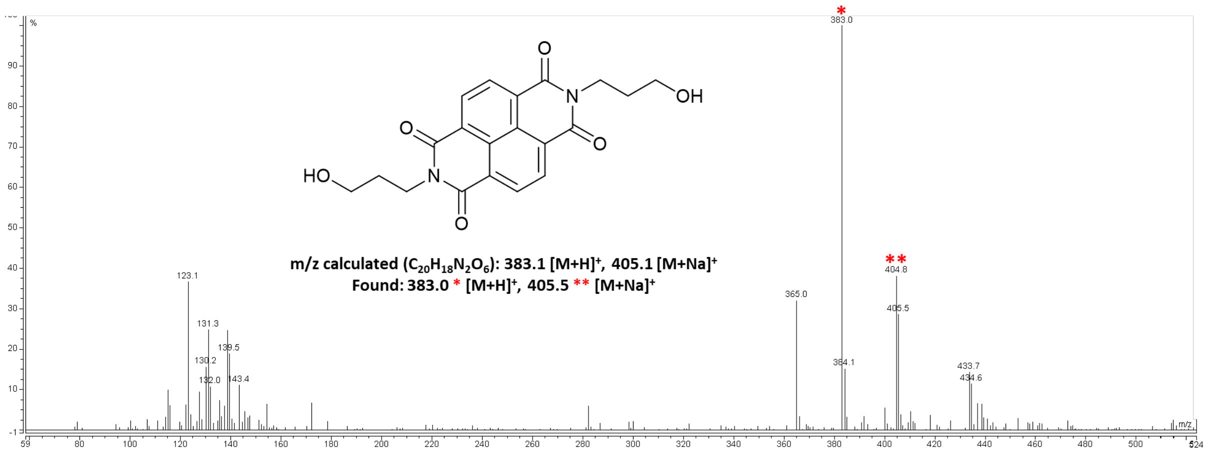

**Figure S 20.** Mass spectrum of **NDI-diol**.

**NDI-DMT**

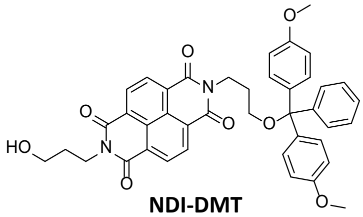

**NDI-DMT**

**NDI-diol** (0.900 g, 2.35 mmol) was dissolved in anhydrous pyridine (40 mL) and stirred under nitrogen. Dimethoxytrityl chloride (0.638 g, 1.88 mmol) was dissolved in anhydrous pyridine (100 mL) and added dropwise at room temperature. The reaction was left stirring under nitrogen at room temperature for 3 hours, after which the pyridine was removed *in vacuo*. The crude residue was dissolved in dichloromethane (120 mL) and washed twice with saturated $NaHCO_3$ (2 x 100 mL). The organic phase was dried over $MgSO_4$, filtered, and dried *in vacuo*. The crude residue was purified with flash chromatography using the Biotage Isolera One, running a hexane and ethyl acetate gradient. This produced a yellow oil (**NDI-DMT**) (0.434 g, 33.7%).

$^1$H NMR (400 MHz, DMSO-$d_6$) δ (ppm): 8.55 (4H, s, Ar-**H**), 7.19 (5H, m, trityl Ar-**H**), 7.08 (4H, d, J = 8.8 Hz, trityl Ar-**H**), 6.69 (4H, d, J = 8.9 Hz, trityl Ar-**H**), 4.56 (1H, t, J = 5.1 Hz, O**H**), 4.13 (4H, q, J = 7.3 Hz, C**H₂**-N), 3.66 (6H, s, COC**H₃**), 3.53 (2H, q, J = 5.9 Hz, C**H₂**-OH), 3.06 (2H, t, J = 4.9, C**H₂**-O-DMT), 1.98 (2H, br. s., C**H₂**-$CH_2$-OH), 1.84 (2H, quin, J = 6.7, C**H₂**-$CH_2$-O-DMT)

$^{13}$C NMR (100 MHz, DMSO-$d_6$) δ (ppm): 162.44, 162.36, 157.79, 144.63, 135.72, 130.33, 129.40, 127.67, 127.55, 126.48, 125.99, 112.86, 85.41, 61.64, 58.97, 54.93, 38.34, 38.09, 30.83, 27.70

MS: m/z calculated ($C_{41}H_{36}N_2O_8$): 685.2477 $[M+H]^+$, 707.2375 $[M+Na]^+$ Found: 685.2565 $[M+H]^+$, 707.2385 $[M+Na]^+$

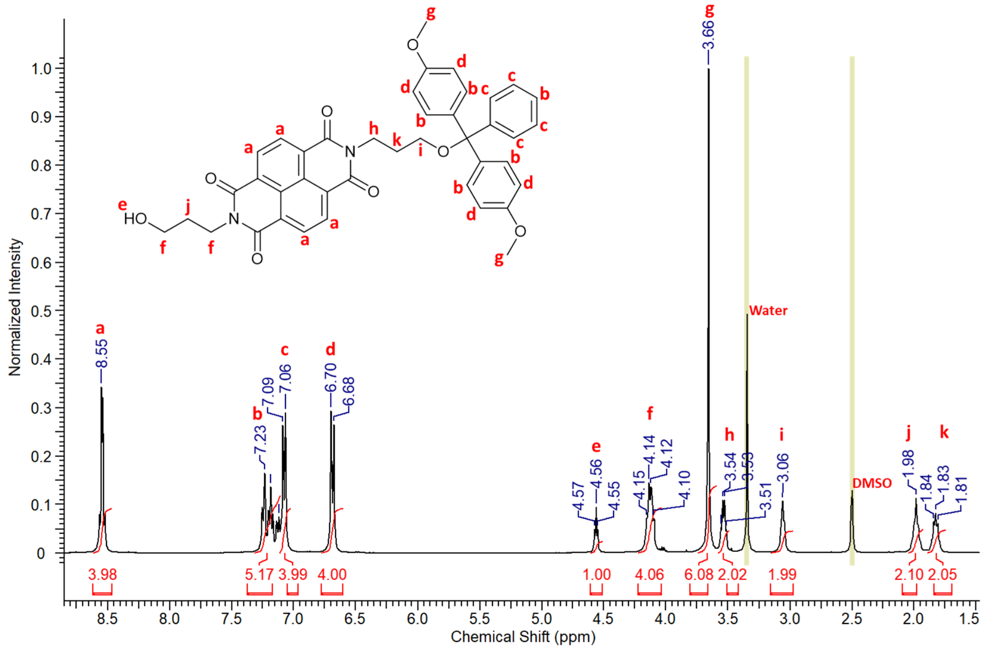

**Figure S 21.** $^{1}H$ NMR spectrum of **NDI-DMT**.

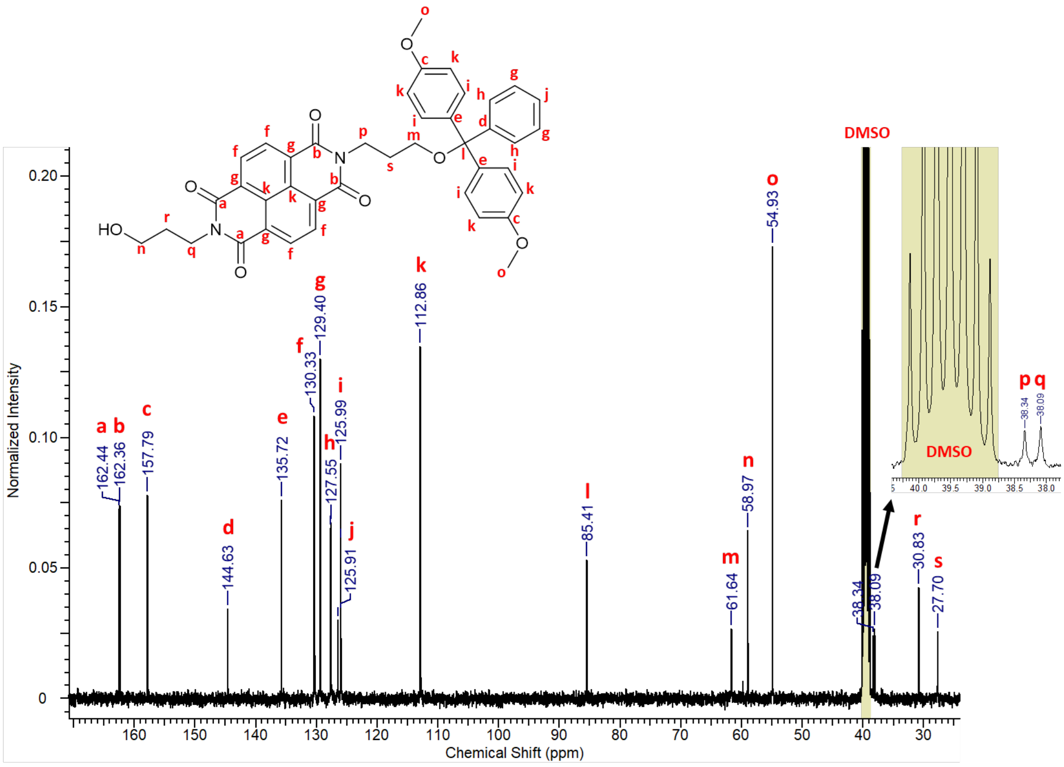

**Figure S 22.** $^{13}C$ NMR spectrum of **NDI-DMT**.

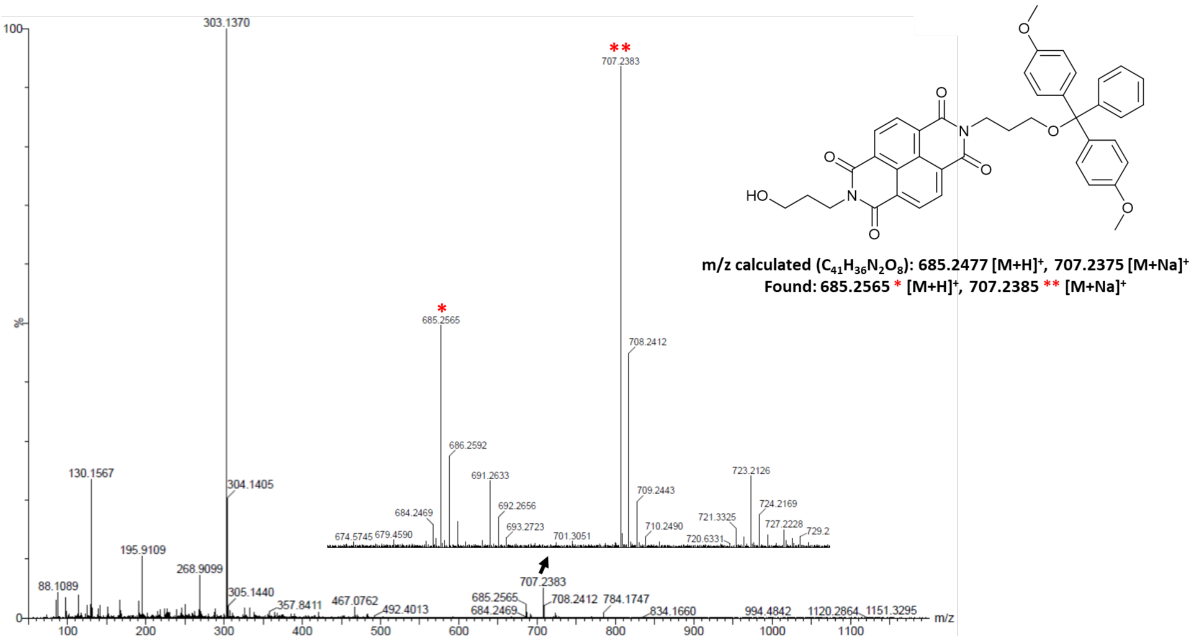

**Figure S 23.** Mass spectrum of **NDI-DMT**.

**NDI phosphoramidite monomer**

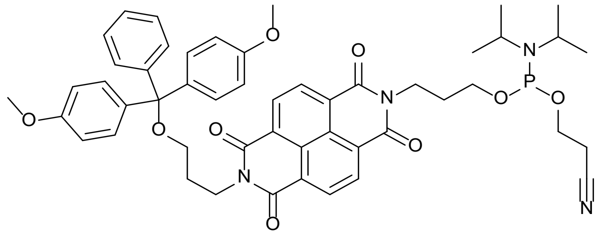

**NDI**

**NDI-DMT** (0.215 g, 0.314 mmol), dimethylaminopyridine (0.004 g, 0.031 mmol), and diisopropylethylamine (273 µL, 1.570 mmol) were all dissolved in anhydrous dichloromethane (25 mL) and stirred under nitrogen. 2-cyanoethyl N, N-diisopropylchlorophosphoramidite (210 µL, 0.942 mmol) was then added, and the reaction mixture was stirred for 2 hours under nitrogen. The dichloromethane was removed *in vacuo*, leaving a yellow oil (0.163 g, 58.7%). $^{31}$P NMR was run on the product, to prove the addition of the phosphoramidite group. No other analysis was run on it because of the air sensitive nature of the product.

$^{31}$P NMR (160 MHz, DMSO-$d_6$ capillary in DCM) δ (ppm): 147.11, 138.36

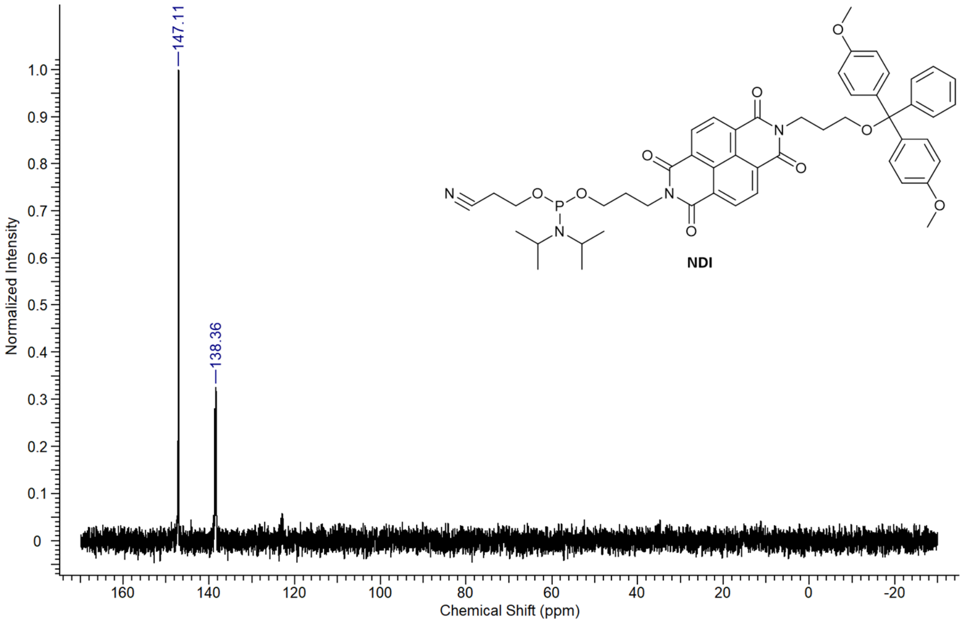

**Figure S 24.** $^{31}$P NMR spectrum of **NDI**.

**DAN-diol**

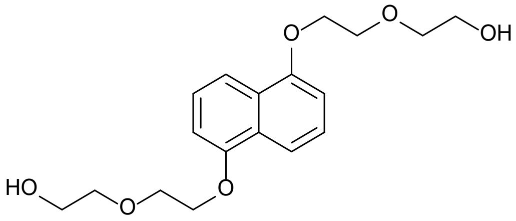

**DAN-diol**

Potassium carbonate (25.86 g, 0.19 mol) was dissolved in anhydrous acetonitrile (160 mL) and stirred under nitrogen. After stirring, 1, 5-dihydroxynaphthalene (3.00 g, 18.73 mmol) was added into the solution. In a separate flask, 2-(2-chloroethoxy)ethanol (4.35 mL, 41.21 mmol) was dissolved in acetonitrile (40 mL) and stirred under nitrogen. The chloroethoxyethanol solution was added dropwise over 30 minutes to the dihydroxynaphthalene solution and left to reflux under nitrogen for 5 days. After reflux, the acetonitrile was removed *in vacuo* and the remaining residue was dissolved in dichloromethane (50 mL). The solution was washed with water (2 x 50 mL), 10% aqueous sodium hydroxide solution (2 x 50 mL), and brine (2 x 50 mL). The organic layer was dried over magnesium sulphate before removing the dichloromethane *in vacuo* to produce a light orange powder (**DAN-diol**) (5.076 g, 80.6%).

$^{1}$H NMR (400 MHz, $CDCl_3$) δ (ppm): 7.89 (2H, d, J = 8.6 Hz, Ar-**H** para), 7.37 (2H, t, J = 8.1 Hz, Ar-**H** meta), 6.87 (2H, d, J = 7.6 Hz, Ar-**H** ortho), 4.31 (4H, t, J = 4.8 Hz, C**H₂**-O-Ar), 4.00 (4H, t, J = 4.8 Hz, $CH_2$-C**H₂**-O), 3.75 (8H, m, C**H₂**-C**H₂**-OH), 2.21 (2H, t, J = 5.8 Hz, O**H**).

$^{13}$C NMR (100 MHz, $CDCl_3$) δ (ppm): 154.29, 126.76, 125.22, 114.63, 105.79, 72.60, 69.79, 67.89, 61.88

MS: m/z calculated ($C_{18}H_{24}O_6$): 337.16 $[M+H]^+$, 359.15 $[M+Na]^+$ Found: 337.2 $[M + H]^+$, 359.1 $[M + Na]^+$

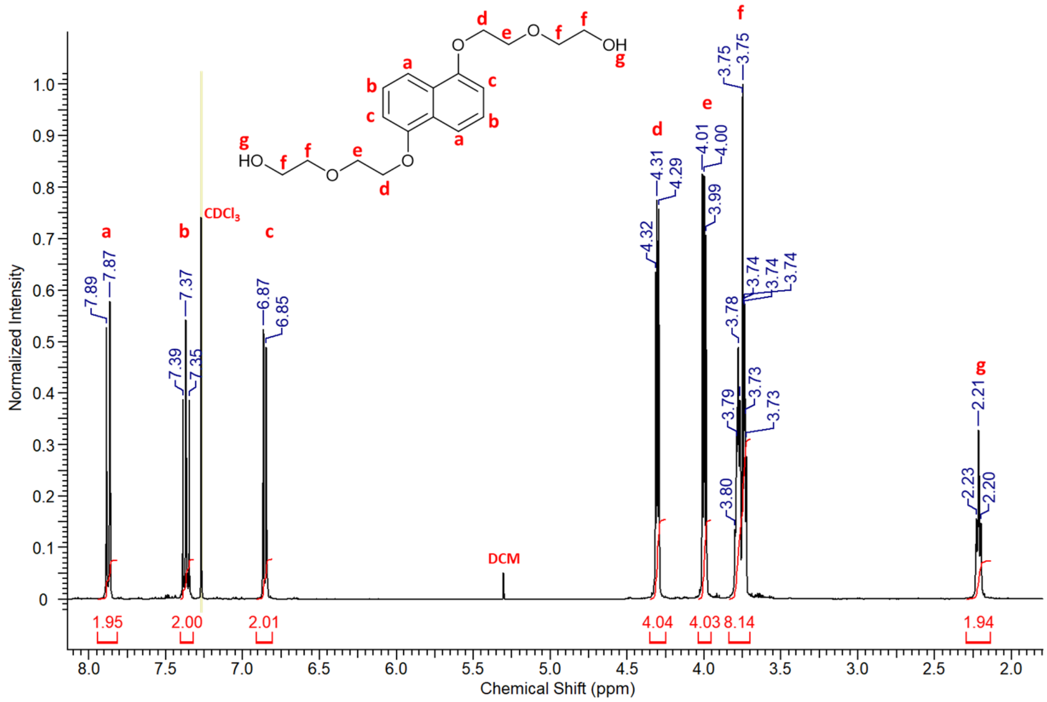

**Figure S 25.** $^{1}$H NMR spectrum of **DAN-diol**.

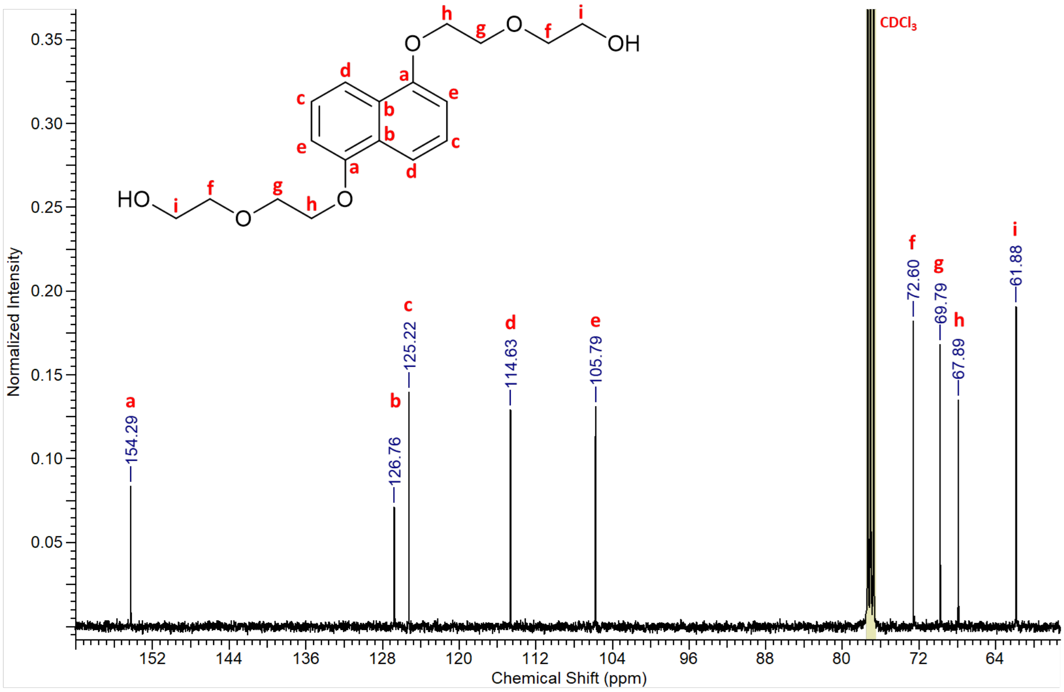

**Figure S 26.** $^{13}$C NMR spectrum of **DAN-diol**.

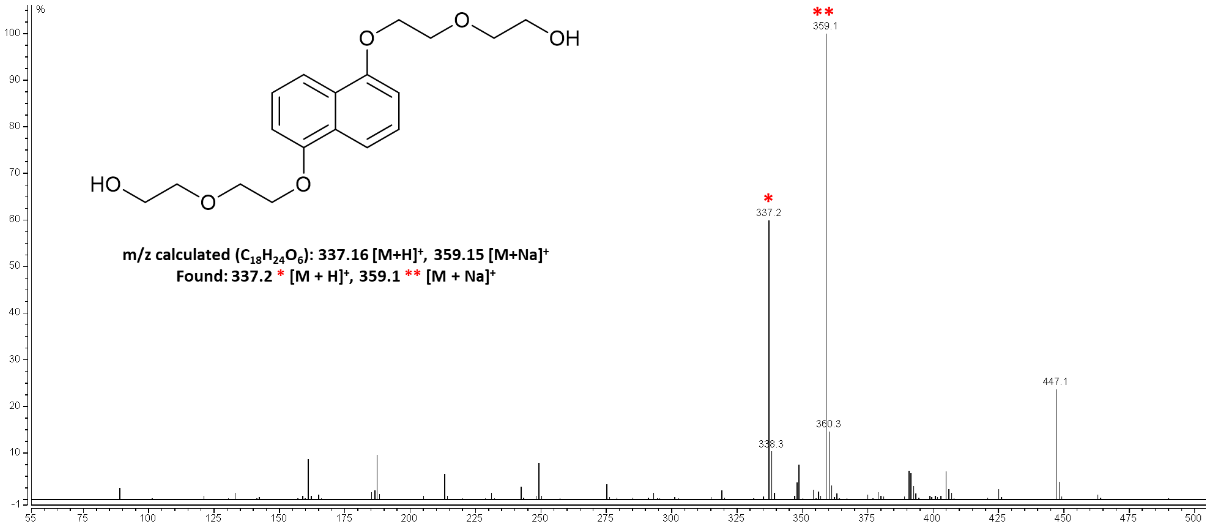

**Figure S 27.** Mass spectrum of **DAN-diol**.

**DAN-DMT**

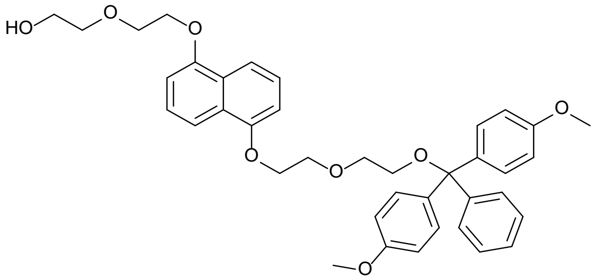

**DAN-DMT**

**DAN-diol** (2.130 g, 7.71 mmol) was dissolved in anhydrous pyridine (40 mL) and left to stir under nitrogen for 30 minutes. Dimethoxytrityl chloride (2.146 g, 7.71 mmol) was dissolved in anhydrous pyridine (10 mL) and added dropwise over 1 hour at room temperature. The reaction was left stirring overnight under nitrogen, after which the pyridine was removed *in vacuo*. The crude residue was purified using flash chromatography using a 10 g SNAP KP-Sil column (Biotage), running a hexane and ethyl acetate gradient (0% to 100% ethyl acetate). This produced a red oil (**DAN-DMT**) (1.521 g, 37.6%).
$^1$H NMR (400 MHz, DMSO-d$_6$) δ (ppm): 7.76 (2H, t, J = 7.7 Hz, Ar-**H**), 7.33 (11H, m, trityl Ar-**H**), 7.01 (2H, dd, J = 14.4, 7.2 Hz, trityl Ar-**H**), 6.82 (4H, d, J = 8.9 Hz, Ar-**H**), 4.66 (1H, t, J = 5.1 Hz, O**H**), 4.31 (2H, t, J = 4.4 Hz, C**H₂**-O-Ar), 4.26 (2H, t, J = 4.6 Hz, C**H₂**-O-Ar), 3.94 (2H, t, J = 4.5 Hz, C**H₂**-CH$_2$-O-Ar), 3.89 (2H, t, J = 4.6 Hz, C**H₂**-CH$_2$-O-Ar), 3.73 (2H, t, J = 4.7 Hz, C**H₂**-OH), 3.70 (6H, s, Ar-C**H₃**), 3.57 (4H, m, C**H₂**-CH$_2$-O-trityl), 3.11 (2H, t, J = 4.8 Hz, C**H₂**-O-trityl)
$^{13}$C NMR (100 MHz, DMSO-d$_6$) δ (ppm): 158.42, 154.41, 145.48, 136.23, 130.12, 128.23, 128.18, 127.04, 126.48, 125.85, 113.55, 106.38, 85.74, 70.44, 69.61, 68.29, 63.42
MS: m/z calculated ($C_{39}H_{42}O_8$): 661.2783 [M+Na]$^+$, Found: 661.2780 [M+Na]$^+$

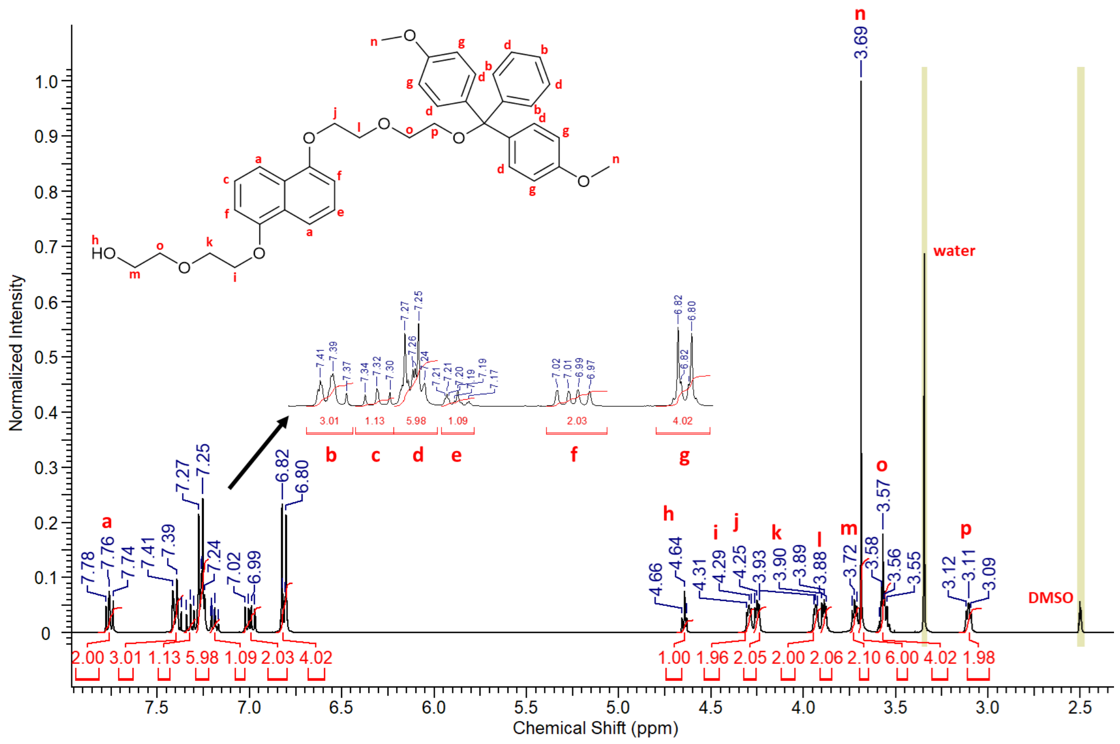

**Figure S 28.** $^{1}$H NMR spectrum of **DAN-DMT**.

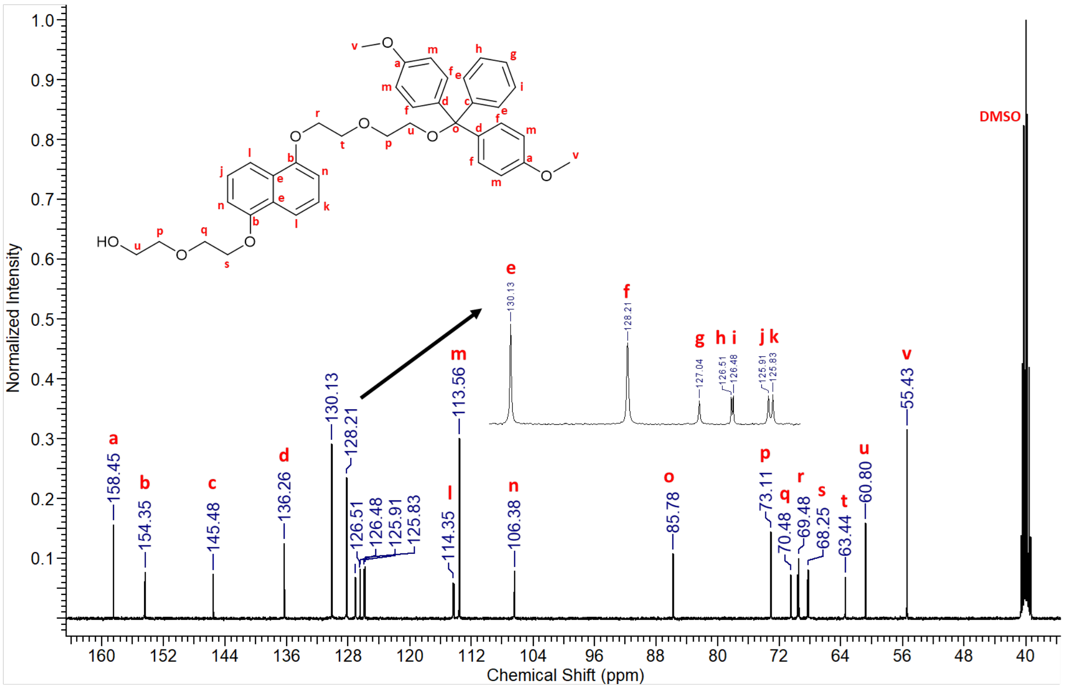

**Figure S 29.** $^{13}$C NMR spectrum of **DAN-DMT**.

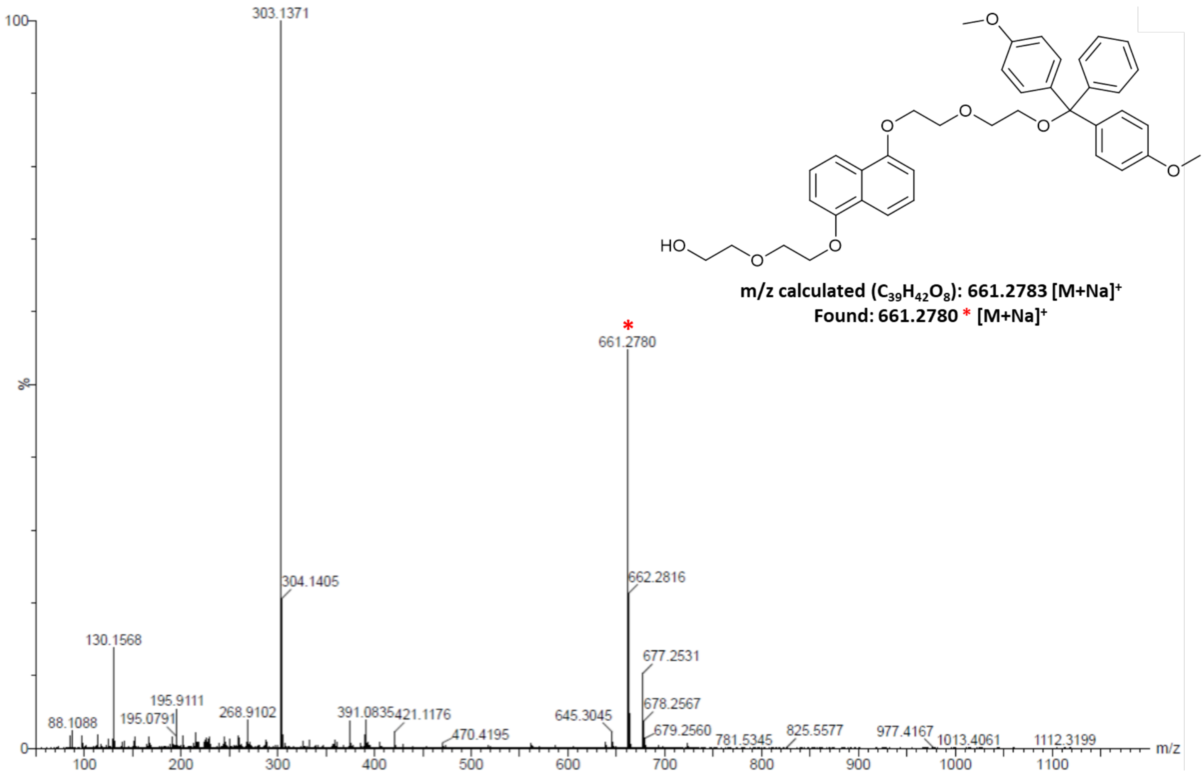

**Figure S 30.** Mass spectrum of **DAN-DMT**.

**DAN phosphoramidite monomer 7**

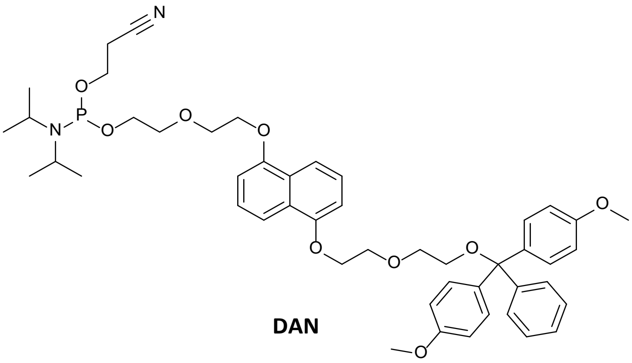

**DAN**

DAN-DEG DMT (0.269 g, 0.421 mmol), dimethylaminopyridine (0.005 g, 0.042 mmol), and diisopropylethylamine (367 µL, 2.106 mmol) were all dissolved in anhydrous dichloromethane (25 mL) and stirred under nitrogen. 2-cyanoethyl N, N-diisopropylchlorophosphoramidite (282 µL, 1.263 mmol) was then added. The reaction mixture was then left to stir for 2 hours under nitrogen. After stirring the dichloromethane was removed *in vacuo*, producing a yellow oil (0.248 g, 70.2%). $^{31}$P NMR was run on the product, to prove the addition of the phosphoramidite group. No other analysis was run on it because of the air sensitive nature of the product.

$^{31}$P NMR (160 MHz, DMSO-$d_6$ capillary in DCM) δ (ppm): 147.86, 138.62, 138.38

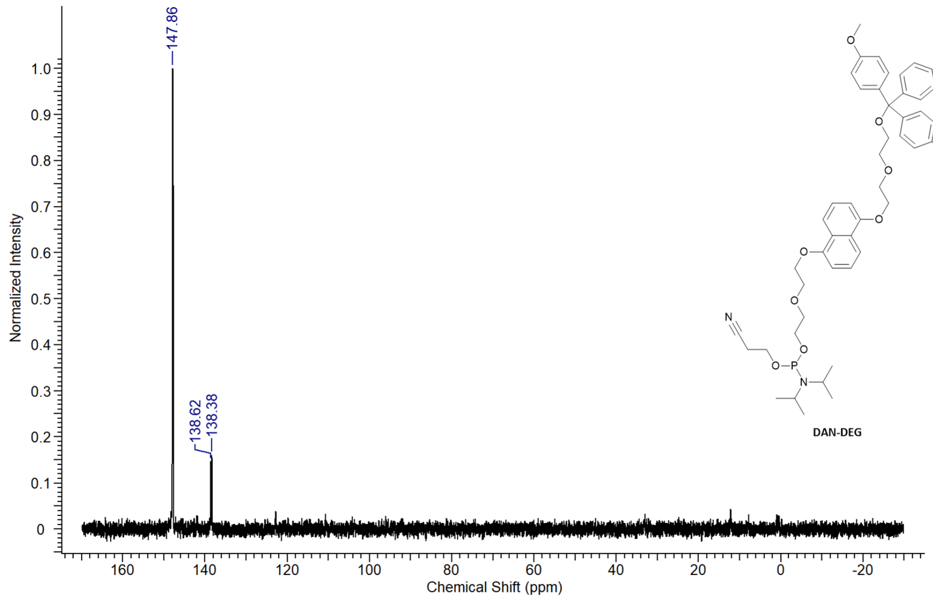

**Figure S 31.** $^{31}$P NMR spectrum of **7**.

#### *2.2 Synthesis of control compound Ch-3*

**Step 1 – Ch-3 precursor**

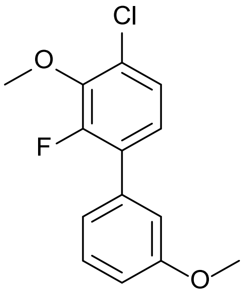

**Ch-3 precursor**

3-Bromoanisole (0.600 g, 3.21 mmol), potassium carbonate (1.330 g, 9.62 mmol), 4-chloro-2-fluoro-3-methoxyphenylboronic acid (0.721 g, 3.53 mmol) and Pd(dppf)$Cl_2$ (0.117 g, 0.16 mmol) were all added sequentially to a 35 mL microwave vial. The vial was fitted with a rubber septum and nitrogen was purged through the reaction mixture. A degassed solution of 1,4-dioxane: water (5:1) (10 mL) was added to the microwave vial which was then sealed. The reaction was heated using microwave irradiation at 100 °C for 18 hours. After this the mixture was cooled, filtered, diluted with ethyl acetate (30 mL) and washed with a 50/50 water/brine solution (3 x 30 mL). The mixture was dried over magnesium sulphate and the ethyl acetate was removed *in vacuo.* The remaining crude was dissolved in ethyl acetate: pentane (1:99) and filtered. The crude residue was purified using flash chromatography using a 10 g SNAP Ultra column (Biotage), running an ethyl acetate and pentane gradient (0% to 20% ethyl acetate). This produced a yellow oil (**Ch-3 precursor**) (0.624 g, 72.9%).

$^{1}$H NMR (400 MHz, $CDCl_3$) δ (ppm): 7.38 (1H, t, J = 8.0 Hz, CH-C**H**-CH-$COCH_3$), 7.22 (1H, dd, J = 8.5, 1.7 Hz, CH-C-C**H**-$COCH_3$), 7.11 (2H, m, Cl-C-C**H**-C**H**), 7.07 (1H, m, C**H**-CH-CH-$COCH_3$), 6.96 (1H, ddd, J = 8.2, 2.5, 0.8 Hz, CH-CH-C**H**-$COCH_3$), 4.01 (3H, d, J = 1.1 Hz, CF-COC**H₃**-CCl), 3.87 (3H, s, CH-COC**H₃**).

$^{13}$C NMR (100 MHz, $CDCl_3$) δ (ppm): 159.56, 154.76, 152.27, 144.85, 144.71, 136.00, 129.53, 129.16, 127.49, 125.06, 124.77, 114.58, 113.53, 61.61, 55.28

MS: m/z calculated ($C_{14}H_{12}ClFO_2$): 265.70 $[M-H]^-$. Found: 265.2 $[M-H]^-$

**Step 2 – Ch-3**

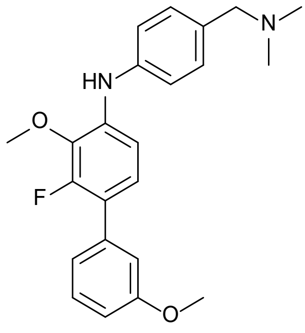

**Ch-3**

**Ch-3 precursor** (0.225 g, 0.844 mmol), caesium carbonate (0.825 g, 2.531 mmol), 4-amino-N, N-dimethylbenzylamine (0.165 g, 1.097 mmol), XPhos (0.040 g, 0.084 mmol), and $Pd(OAc)_2$ (0.009 g, 0.042 mmol) were all added sequentially to a 10 mL microwave vial. The vial was fitted with a rubber septum and nitrogen was purged through the reaction mixture. A degassed solution of 1,4 – dioxane (6 mL) was added to the microwave vial which was then sealed. The reaction was heated using microwave irradiation at 100 °C for 24 hours. After this the mixture was cooled, filtered, diluted with ethyl acetate (30 mL) and washed with a 50/50 water/brine solution (3 x 30 mL). The mixture was dried over magnesium sulphate and the ethyl acetate was removed *in vacuo.* The remaining crude was dissolved in dichloromethane: methanol (9:1) and filtered. The crude residue was purified using flash chromatography using a 10 g SNAP Ultra column (Biotage), running a dichloromethane and methanol gradient (0% to 10% methanol). This produced a yellow oil (**Ch-3**) (0.287 g, 89.4%).

$^1$H NMR (400 MHz, DMSO-$d_6$) δ (ppm): 7.89 (1H, s, NH), 7.36 (1H, t, J = 7.9 Hz, CH-C**H**-CH-C($OCH_3$)), 7.16 (4H, m, C**H**-C(NH)-C**H**, NH-C-C**H**-CH, C-CH-C**H**-C($OCH_3$)), 7.07 (4H, m, NH-C-C**H**, C**H**-C($CH_2$N)-C**H**, CF-C-C-C**H**), 6.92 (1H, ddd, J = 8.3, 2.6, 0.9 Hz, (C**H**-C($OCH_3$)-CH), 3.84 (3H, s, CF-C(OC**H₃**)-C(NH)), 3.80 (3H, s, CH-C(OC**H₃**)-CH), 3.31 (2H, s, C-C**H₂**-N-($CH_3$)$_2$), 2.13 (6H, s, CH2-N-**($CH_3$)$_2$**)

$^{13}$C NMR (100 MHz, DMSO-$d_6$) δ (ppm): 159.33, 154.49, 152.05, 141.07, 138.14, 136.71, 136.58, 131.61, 129.58, 124.51, 120.80, 119.86, 118.86, 113.98, 112.64, 110.94, 63.07, 60.99, 60.93, 55.12, 44.95

MS: m/z calculated ($C_{23}H_{25}FN_2O_2$): 381.19 $[M+H]^+$. Found: 381.14 $[M+H]^+$

#### 2.3 Modification of TentaGel beads

**Synthesis of -OH modified TG-beads**

*N*,*N*-Diisopropylcarbodiimide (0.30 g, 2.38 mmol) was dissolved in DMF (10 mL), followed by the addition of 10-hydroxydecanoic acid (0.60 g, 3.19 mmol). In a separate vessel, TentaGel® M $NH_2$ Monosized Amino TentaGel® Microspheres (0.15 g) were dissolved in DMF (10 mL) and stirred for 30 minutes to allow the spheres to swell. After stirring, the mixture of DIC and 10-hydroxydecanoic acid was added to the TentaGel® mixture, and the reaction was stirred at room temperature for 3 hours. The reaction mixture was centrifuged, and the supernatant was removed and discarded. The yellow solid was filtered and washed with DMF (2 x 10 mL) and ethanol (2 x 10 mL). A Kaiser test was performed on the dried solid, producing a yellow colour, meaning there were no free primary amines left in the solution and the reaction was successful. The final product was a pale-yellow solid (0.147 g, 98 %).

**Synthesis of fluorescein TG-Beads**

*N*,*N*-Diisopropylcarbodiimide (0.15 g, 1.19 mmol) was dissolved in DMF (10 mL). In a separate vessel, TentaGel® M $NH_2$ Monosized Amino TentaGel® Microspheres (0.05 g) were dissolved in DMF (10 mL) and stirred for 30 minutes to allow the spheres to swell. After stirring, the mixture of DIC was added to the TentaGel® spheres, and the mixture was left stirring at room temperature for 30 minutes before 5(6)-carboxyfluorescein (0.04 g, 0.11 mmol) was added directly to the reaction mixture. The reaction was stirred at room temperature for 3 hours. After stirring, the mixture was centrifuged, and the supernatant was removed and discarded. The solid was filtered and washed with DMF (2 x 10 mL) and ethanol (2 x 10 mL). The final product was an orange solid (TG-CF100) (0.042 g, 84%).

Fluorescently tagged TentaGel® microspheres from 75 – 1% followed the same experimental procedure with a difference in 5(6)-carboxyfluorescein added (Table S 2).

**Table S 2.** Quantities for partial labelling of TentaGel® beads with fluorescein.

| TG-Bead Sample | Weight fluorescein added (g) | Product Weight (g) | Yield (%) |
|---|---|---|---|
| **TG-CF75** | 0.030 | 0.036 | 72 |
| **TG-CF50** | 0.020 | 0.035 | 70 |
| **TG-CF25** | 0.010 | 0.036 | 72 |
| **TG-CF10** | 0.004 | 0.043 | 86 |
| **TG-CF5** | 0.002 | 0.038 | 76 |
| **TG-CF1** | 0.0004 | 0.040 | 80 |

**Synthesis of rhodamine TG-Beads**

*N*,*N*-Diisopropylcarbodiimide (0.15 g, 1.19 mmol) was dissolved in DMF (10 mL). In a separate vessel, TentaGel® M $NH_2$ Monosized Amino TentaGel® Microspheres (0.05 g) were dissolved in DMF (10 mL) and stirred for 30 minutes to allow the spheres to swell. After stirring, the mixture of DIC was added to the TentaGel® spheres, and the mixture was left stirring at room temperature for 30 minutes before rhodamine B (0.04 g, 0.08 mmol) was added directly to the reaction mixture. The reaction was stirred at room temperature for 3 hours. After stirring, the mixture was centrifuged, and the supernatant was removed and discarded. The solid was filtered and washed with DMF (2 x 10 mL) and ethanol (2 x 10 mL). The final product was a pink solid (TG-RB100) (0.039 g, 78%).

Fluorescently tagged TentaGel® microspheres from 75 – 1% followed the same experimental procedure with a difference in rhodamine B weight (Table S 3).

**Table S 3.** Quantities for partial labelling of TentaGel® beads with rhodamine.

| TG-Bead Sample | Weight Added (g) | Product Weight (g) | Yield (%) |
|---|---|---|---|
| **TG-RB75** | 0.030 | 0.047 | 94 |
| **TG-RB50** | 0.020 | 0.040 | 80 |
| **TG-RB25** | 0.010 | 0.043 | 86 |
| **TG-RB10** | 0.004 | 0.036 | 72 |
| **TG-RB5** | 0.002 | 0.042 | 84 |
| **TG-RB1** | 0.0004 | 0.022 | 44 |

#### 2.4 Automated oligomer synthesis

**General Expedite™ 8909 DNA Synthesiser set up for the synthesis of oligomer library.**

The phosphoramidite monomers were dissolved in 5 mL of anhydrous DCM and placed into the appropriate synthesiser lines. The other reagents on the synthesiser were: Cap A Mix (THF/Pyridine/acetic anhydride 8:1:1), Cap B Mix (10% methylimidazole in THF), deblock (3% trichloroacetic acid in DCM) and ETT activator solution (0.25 M, 5-ethylthio-1H-tetrazole in acetonitrile). Before synthesis a leak test is run to ensure there are no nitrogen leaks from the synthesiser lines, after which all the reagents are flushed through the system to prime them. The beads are then added to the column, attached to the synthesiser, and primed with anhydrous acetonitrile. Validate XP software (version 5.4.15) is used to select the appropriate sequence and protocol for oligomer synthesis, while the trityl monitor is used to monitor each addition.

**Synthesis of oligomer library**

Modified (-OH) TG-beads (0.166 g) were added to a synthesiser column. This was enough to synthesise 300 copies of each sequence. The first round of addition was the addition of the photocleavable linker (PC-Linker); after the initial addition of the PC-Linker, the TG-beads were split across seven synthesiser columns (0.023 – 0.024 g per column) for each monomer addition. After the seven monomers were added to an individual column, all TG-beads were mixed and split out to seven individual columns again. The final weight after library synthesis was 0.148 g, resulting in an 11% loss of library mass, 268 copies of each sequence in the library, and 220,832,137 TG-Beads in total.

Table S 4 shows the response values given from the trityl monitor. The addition of each monomer typically had a response between $10^4$ and $10^6$, with considerable variation between each round. When the signal is below $10^4$ the trityl monitor gives a response of 1, but the visual conformation of the DMT/trityl cation cleavage manifested as a dark orange colour similar in intensity to other couplings suggests these were successful coupling reactions as well. In our experience, the particular trityl monitor setup in our instrument is susceptible to false negative readings due to occasional bubbles passing through the system.

**Table S 4.** Trityl monitor values for library synthesis

| | Addition | | | | | | |
|---|---|---|---|---|---|---|---|
| **Monomer** | **1st** | **2nd** | **3rd** | **4th** | **5th** | **6th** | **7th** |
| 1 | $1.64 \times 10^4$ | 1 | 1 | $3.71 \times 10^5$ | $3.12 \times 10^5$ | $3.17 \times 10^5$ | 1 |
| 2 | $3.24 \times 10^5$ | $6.09 \times 10^5$ | $4.84 \times 10^5$ | $2.99 \times 10^5$ | $6.43 \times 10^5$ | $1.38 \times 10^5$ | $6.94 \times 10^5$ |
| 3 | $8.16 \times 10^5$ | $9.83 \times 10^5$ | $6.04 \times 10^5$ | $4.24 \times 10^5$ | $6.02 \times 10^5$ | 1 | $7.50 \times 10^5$ |
| 4 | $5.18 \times 10^5$ | $6.63 \times 10^5$ | $3.18 \times 10^5$ | $2.46 \times 10^5$ | $6.09 \times 10^5$ | $2.24 \times 10^6$ | $6.27 \times 10^5$ |
| 5 | $6.96 \times 10^5$ | $1.24 \times 10^6$ | $1.63 \times 10^4$ | $2.36 \times 10^5$ | $6.13 \times 10^5$ | 1 | $5.88 \times 10^5$ |
| 6 | $4.48 \times 10^5$ | $1.25 \times 10^6$ | $4.66 \times 10^5$ | $2.37 \times 10^5$ | $7.27 \times 10^5$ | $2.21 \times 10^5$ | 1 |
| 7 | $1.12 \times 10^6$ | $6.09 \times 10^5$ | $2.94 \times 10^5$ | $3.11 \times 10^5$ | $3.94 \times 10^5$ | 1 | $7.50 \times 10^5$ |

**Resynthesis of selected sequences**

CPG spheres (0.021 g) were added to a synthesiser column and the synthesiser was set up as described above. The sequences input into the DNA synthesiser were as shown in Table S 5.

**Table S 5.** Oligomer syntheses.

| Oligomer | Sequence (OH – P) | Repeat syntheses (n = ) |
|---|---|---|
| **48** | C12-DAN-DEG-HEG-HEG-HEG | 2 |
| **49** | BPA-NDI-C12-C12-C12-NDI | 2 |
| **59** | cYY-cYY-HEG-HEG-HEG | 2 |
| **81** | C12-C12-C12-C12-HEG-C12-C12 | 5 |
| **144** | DAN-DEG-HEG-DAN-DEG-NDI-HEG | 2 |
| **200** | C12-NDI-BPA-cYY | 3 |

The trityl monitor was used to monitor the couplings and multiple syntheses of each oligomer were performed (repeat syntheses). Each synthesis was run to completion with 'trityl on' meaning the final dimethoxytrityl (DMT) group was not cleaved (as DMT was used for subsequent quantification).

To cleave the oligomers from the CPG solid support, the solids were removed from the synthesiser column and transferred into a screw-cap centrifuge tube, with repeated syntheses of the same oligomer combined into one centrifuge tube. To the centrifuge tube 1.5 mL of ammonia (35%) was added and the oligomers were incubated in a water bath at 60 °C for 8 hours. The oligomers were left to dry and re-suspended in 1 mL acetic acid solution (80%).

Using the Nanodrop UV-Vis spectrophotometer, the absorbance at 500 nm was collected for DMT standards dissolved in 80% acetic acid at 5 µM/mL, 2 µM/mL, 1.5 µM/mL, 1 µM/mL, 0.75 µM/mL, 0.5 µM/mL, 0.25 µM/mL, 0.1 µM/mL, 0.05 µM/mL, 0.01 µM/mL, and all oligomer data were collected in triplicate. The standard data was used to create a calibration curve and from this the concentration of oligomers was determined. To remove the cleaved -DMT/trityl protecting group, the oligomer was left to dry and partitioned between 0.5 mL dichloromethane and water. The organic layer was removed, and the water layer was left to dry before being resuspended in KRAS Protein Buffer.

After determining their concentration each oligomer solution was desalted using Pierce™ C18 spin tips and columns following the manufacturers protocol.[1] Briefly, the C18 tips were wetted with 20 µL of Milli Q® water containing 0.1% TFA, centrifuging at 1000 g for 1 minute. The C18 tips were then equilibrated with 20 µL of Milli Q® water containing 0.1% TFA, centrifuging at 1000 g for 1 minute. The sample (between 20 and 50 µL) was then added to the C18 tips and centrifuged at 1000 g for 1 minute. The tip was washed by adding 20 µL of Milli Q® water containing 0.1% TFA, centrifuged at 1000 g for 2 x 1 minute. The sample was eluted using 20 µL of acetonitrile containing 0.1% TFA and centrifuging at 1000 g for 2 x 1 minute. The sample was dried and resuspended in 20 µL of KRAS Protein buffer.

A 15% polyacrylamide gel (with 4% stacking gel) was polymerised with APS (100 µL) and TEMED (10 µL). Oligomers and the DNA standard were made up to 100 µL in SDS sample buffer (see 5.2.3.1) and heated at 95 °C for 15 minutes. The samples were left to cool to room temperature and then 10 µL of each oligomer and 10 µL of 2 µM DNA standard were loaded onto the gel and run for 60 minutes at 100 V using a tris-glycine SDS running buffer (see 5.2.3.1). The gel was stained with a Pierce™ Silver Staining Kit and images were taken using an Epsom scanner.

### 3 Protein expression

For generation of recombinant KRas proteins, sequences encoding KRas4B(1-169) [Uniprot # P01116-2] and KRas4B(1-169)[G12D], optimised for expression in *E.coli*, were subcloned into the *NcoI* & *XhoI* sites of the pET28 (Novagen) expression vector with an N-terminal His6-Avi·tag-TEV- fusion tag. Plasmids were transformed into BL21(DE3) Gold competent *E.coli* cells (Agilent). Cells were cultured in Terrific broth (Formedium) containing 25 µg/mL Kanamycin. Once cultures had reached on OD600nm of 0.6, expression was induced with 0.2 mM IPTG and performed at 30°C for 4 hrs. Harvested cells were resuspended in 20 mM HEPES (pH 7.3), 300 mM NaCl, 2 mM TCEP, 5 mM $MgCl_2$, 10 mM imidazole. The cell suspensions were supplemented with 1 mM PMSF, 2 µg/ml Leupeptin, 2 µg/mL Pepstatin and Protease inhibitor cocktail tablets to prevent proteolytic degradation. The cell suspension was lysed by mechanical homogenization and centrifuged at 20000 rpm for 90 mins to pellet insoluble cell debris. The clarified supernatant was passed through a 5 mL HisTrap FF column (Cytiva), washed with 20 mM HEPES (pH 7.3), 300 mM NaCl, 2 mM TCEP, 5 mM $MgCl_2$, 10 mM imidazole, then eluted with 20 mM HEPES (pH 7.3), 300 mM NaCl, 2 mM TCEP, 5 mM $MgCl_2$, 300 mM imidazole. The eluted protein was then buffer exchanged into SEC buffer (10 mM Tris (pH 7.5), 50 mM NaCl, 2 mM $MgCl_2$) and incubated with 400 nM BirA, 1 mM Biotin and 2 mM ATP to biotinylate the Avi·tag. The biotinylation reaction was further purified by size exclusion chromatography (SEC) on a HiLoad 16/600 Superdex 75 pg column (Cytiva) with SEC buffer.

To generate uniformly loaded GDP samples, the purified protein was incubated with a 20-fold molar excess of GDP and 25 mM EDTA at 4°C for 2h. $MgCl_2$ was then added to a final concentration of 50 mM at 4°C for 2h. The protein was then buffer exchanged into SEC buffer.

To generate uniformly loaded GMPPnP samples, the purified protein was exchanged into Exchange buffer (40 mM Tris-pH 8, 200 mM $(NH_4)_2SO_4$, 5 mM DTT, 4 uM $ZnCl_2$). GMPPnP was added to a 5-fold molar excess, then alkaline phosphatase (Roche) was added at 5 U per mg of target protein. The mixture was left at 4°C overnight. The sample was then purified by SEC on a HiLoad 16/600 Superdex 75pg column (Cytiva) with SEC buffer.

For generation of recombinant RAF1 RBD, a sequence encoding RAF1(51-131) [Uniprot # P04049-1] optimised for expression in *E.coli*, was subcloned into the NcoI & XhoI sites of the pET28 expression vector with a C-terminal His10-TEV-Avi·tag. The plasmid was transformed into CVB101 (Avidity) previously lysogenized with the λ phage DE3 by using the λDE3 Lysogenization Kit (Merck). Cells were cultured in Terrific broth supplemented with 50 uM biotin. Once cultures had reached on OD600 nm of 0.6, expression was induced with 0.2 mM IPTG and performed at 30°C for 16h. Harvested cells were resuspended in 50 mM Tris HCl (pH 8), 300 mM NaCl, 1 mM $MgCl_2$, 100 µg/mL PMSF, 1.5 mM DTT, 15 mM Imidazole. The cell suspension was lysed by mechanical homogenization and centrifuged at 20000 rpm for 90 mins to pellet insoluble cell debris. Clarified supernatant was passed through a 5 mL HisTrap FF column (Cytiva), washed with 50 mM Tris-HCl (pH 8), 300 mM NaCl, 30 mM Imidazole, then eluted with 50 mM Tris-HCl (pH 8), 300 mM NaCl, 300 mM Imidazole. Eluted protein was then purified by SEC on a HiLoad 16/600 Superdex 75pg column with 20 mM Tris HCl (pH 7.5), 150 mM NaCl, 5 mM $MgCl_2$.

For generation of GFP-tagged RAF1 RBD, a sequence encoding RAF1(51-131) optimised for expression in *E.coli*, was subcloned into the NcoI & XhoI sites of the pET28 expression vector, with an N-terminal His8-His8-TEV- tag and a C-terminal GFP (1- 238) [Uniprot # P42212] fusion. The plasmid was transformed into BL21(DE3) cells (Novagen). Cells were cultured in Terrific broth with 25 µg/mL Kanamycin. Once cultures had reached on OD600nm of 0.6, expression was induced with 0.2 mM IPTG and performed at 30C for 16h Purification was performed as with the His10-TEV-Avi-RAF1 RBD, except a TEV cleavage step was included prior to SEC to remove the His8-His8- tag.

All protein samples were quantified from absorbance at 280 nm based upon predicted molecular weights and molar extinction coefficients. Samples were separated into single use aliquots, flash frozen then stored at -80°C until use.

### 4 Fluorescence-Activated Bead Sorting

#### 4.1 *General flow cytometer calibration and set up.*

Each laser of the BD FACSJazz™ flow cytometer was calibrated using Sphero™ Rainbow Calibration Beads (8-peak). Before analysing any sample, unlabelled -OH modified TG-beads (5 mg) were dispersed in 2 mL sheath fluid and analysed to check the calibration. To calibrate the machine for both 2-way and 96-well plate sorting, BD FACS™ Accudrop Beads were used. Fluorescein and 5(6)-carboxyfluorescein were excited by the 488 nm laser and detected by 513/17 and 542/27 nm bandpass filters. Rhodamine B and rhodamine Red™-X were excited by the 561 nm laser and detected by 585/29 and 600 nm bandpass filters. All flow cytometry data was processed using BD FACS™ Sortware program.

#### 4.2 *FABS analysis of fluorophore-labelled beads*

**FACS analysis of 5(6)-carboxyfluorescein TG-Beads.**

The flow cytometer was calibrated as above. All samples (5 mg) were dispersed in 2 mL of sheath fluid. After checking the calibration with unlabelled TG-Beads, samples, TG-CF100, TG-CF75, TG-CF50, TG-CF25, TG-CF10, TG-CF5 and TG-CF1 were analysed respectively. A clear progression in fluorescence intensity was seen using the 488 nm laser which is tuned to the fluorophore (Figure S 32) but not with the 561 nm laser tuned to rhodamine (Figure S 33).

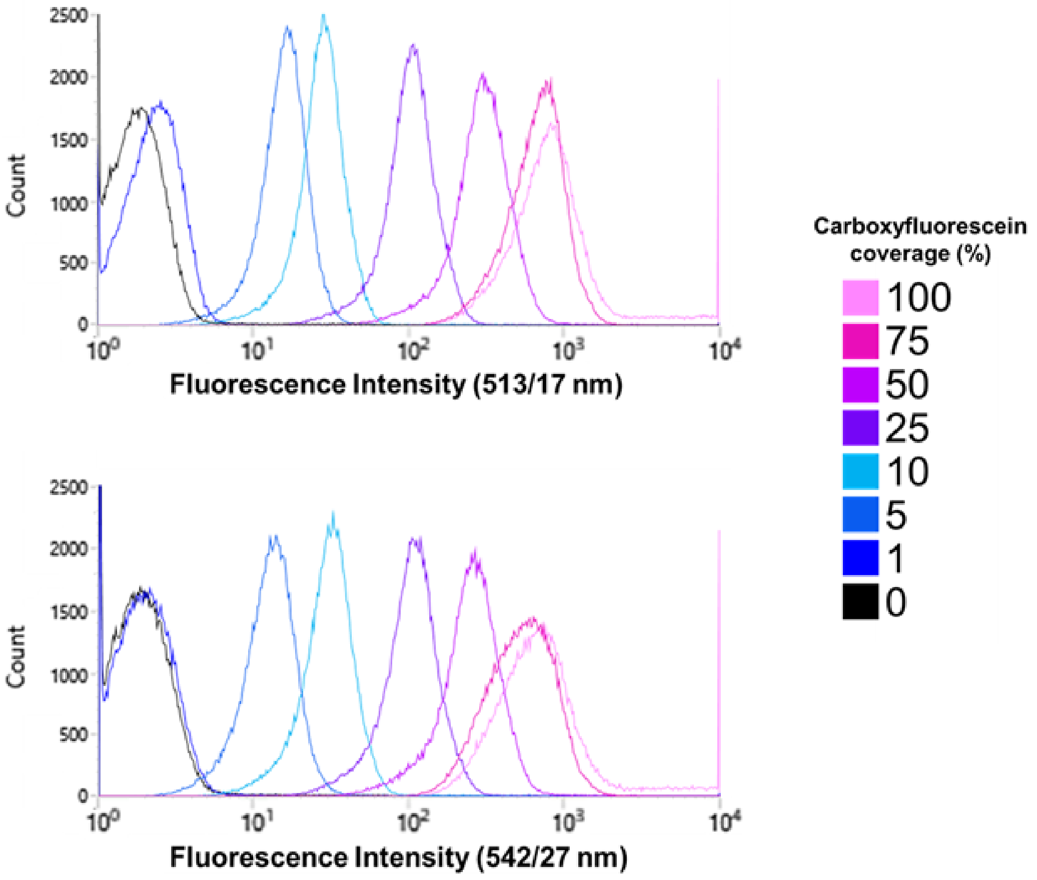

**Figure S 32.** FABS data of 5(6)-carboxyfluorescein tagged microspheres using the 488 nm laser.

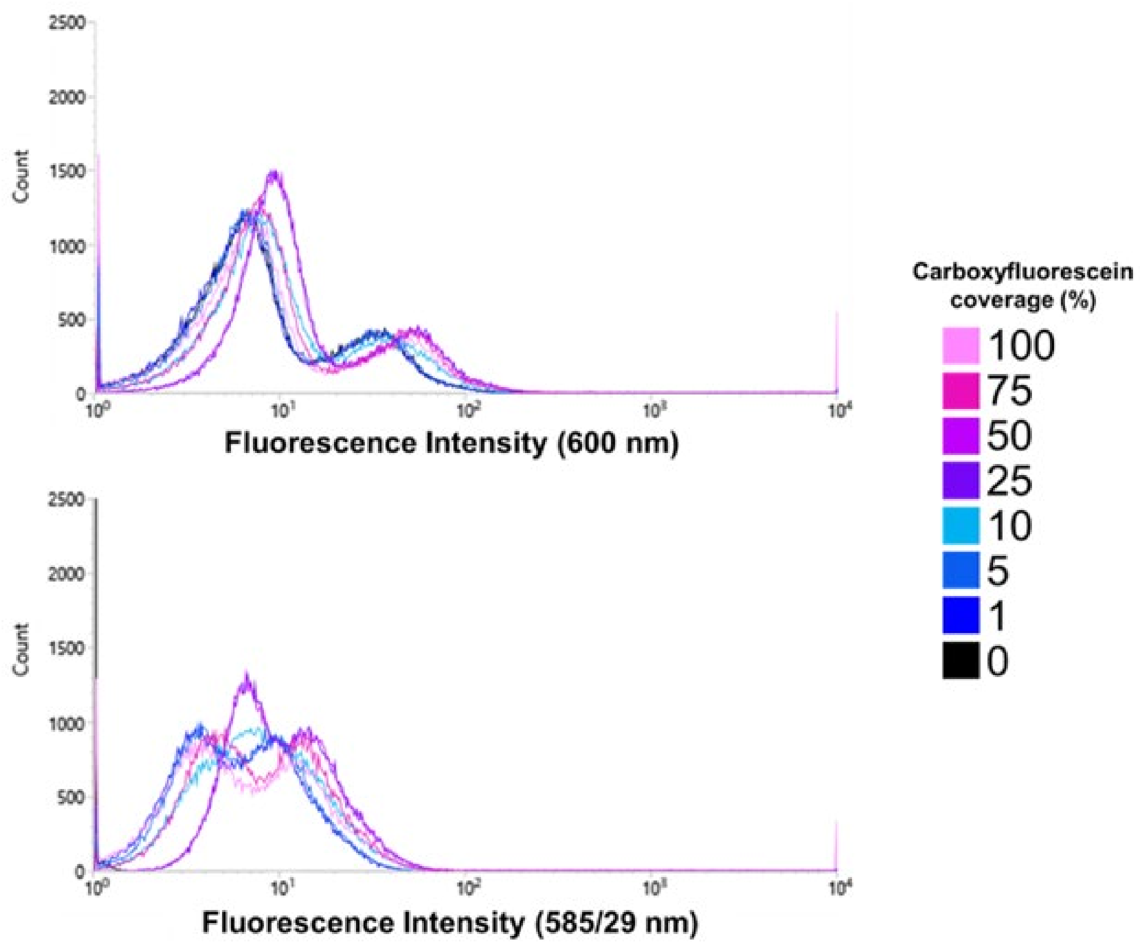

**Figure S 33.** FABS data of 5(6)-carboxyfluorescein tagged microspheres using the 561 nm laser.

**FACS analysis of rhodamine TG-Beads.**

The flow cytometer was calibrated as above. All samples (5 mg) were dispersed in 2 mL of sheath fluid. After checking the calibration with unlabelled TG-Beads, samples, TG-RB100, TG-RB75, TG-RB50, TG-RB25, TG-RB10, TG-RB5 and TG-RB1 were analysed respectively. A clear progression in fluorescence intensity was seen using the 561 nm laser which is tuned to the fluorophore (Figure S 34) but not with the 488 nm laser tuned to fluorescein (Figure S 35).

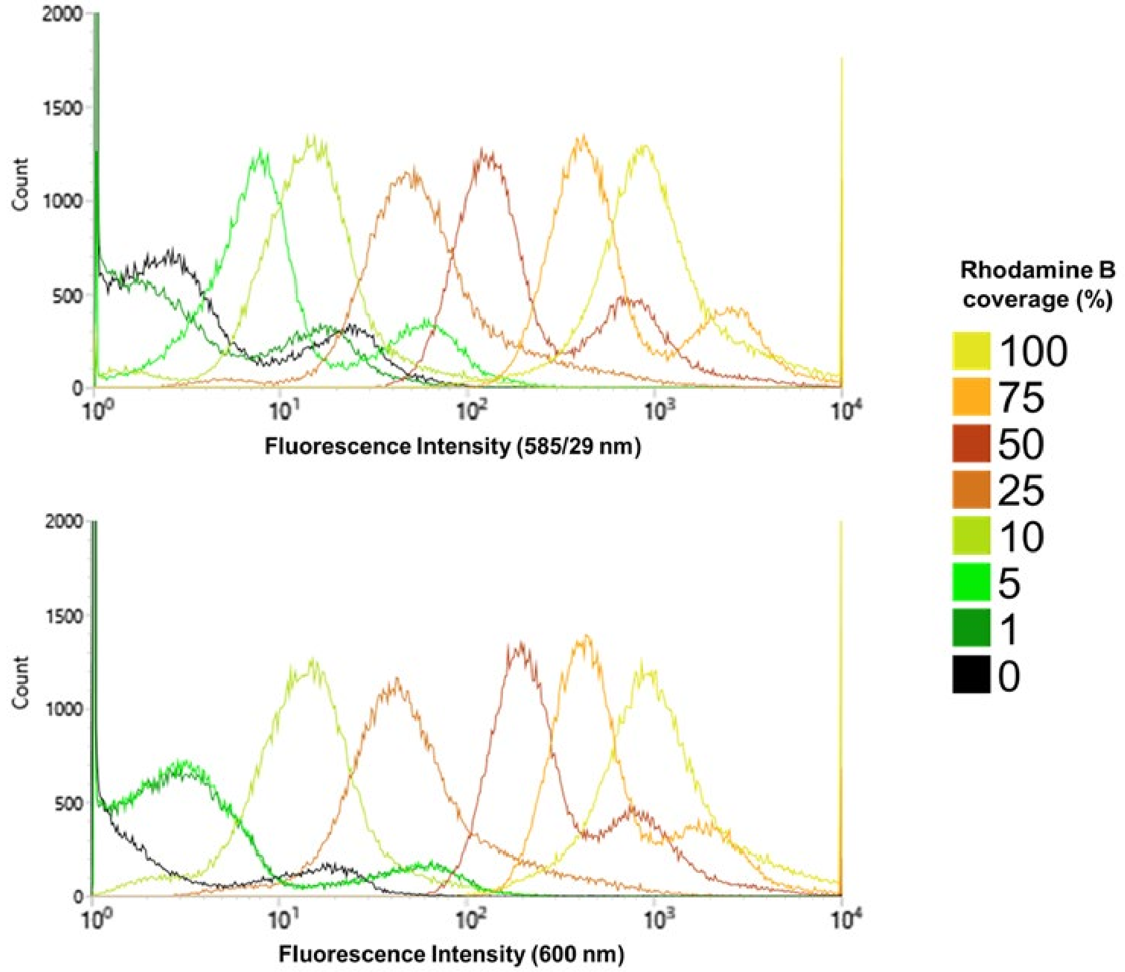

**Figure S 34.** FABS data of rhodamine B tagged microspheres using the 561 nm laser.

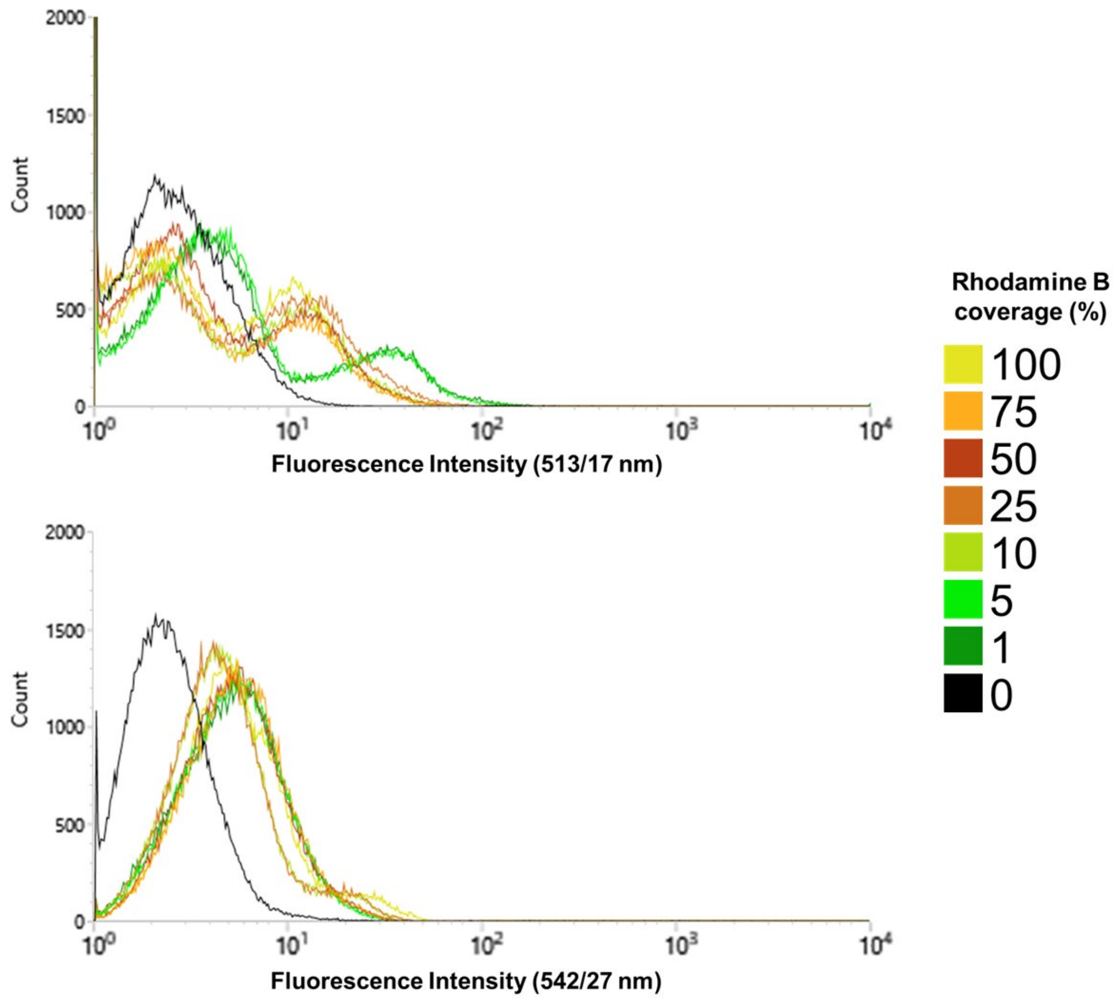

**Figure S 35.** FABS data of rhodamine B tagged microspheres using the 488 nm laser.

**96-well plate sort of 100% (5)6-carboxyfluorescein and 100% rhodamine B TG-Bead mixture.**

The flow cytometer was calibrated as above. TG-CF100 (2 mg) and TG-RB100 (2 mg) were dispersed in 2 mL of sheath fluid. Using the 488 nm laser and bandpass filters, a selection gate for the highest fluorescence signals was created for TG-CF100. Using the 561 nm laser and bandpass filters, a selection gate for the highest fluorescence signals was created for TG-RB100. A ½ log digression starting at 50,000 TG-beads progressing down to 5 TG-Beads was used to collect both TG-CF100 (wells A1-9, B1-9, C1-9, and D1-9) and TG-RB100 (wells E1-9, F1-9, G1-9, and H1-9). A fluorescence plate reader (ClarioStar, BMG LabTech) was used to measure the emission spectrum of fluorescein (500 – 600 nm) and rhodamine (570 – 670 nm). GraphPad Prism software (version 9.5.1) was used to analyse the plate reader data and calculate the area under the curve (AUC) of all emission spectra (Figure S 36).

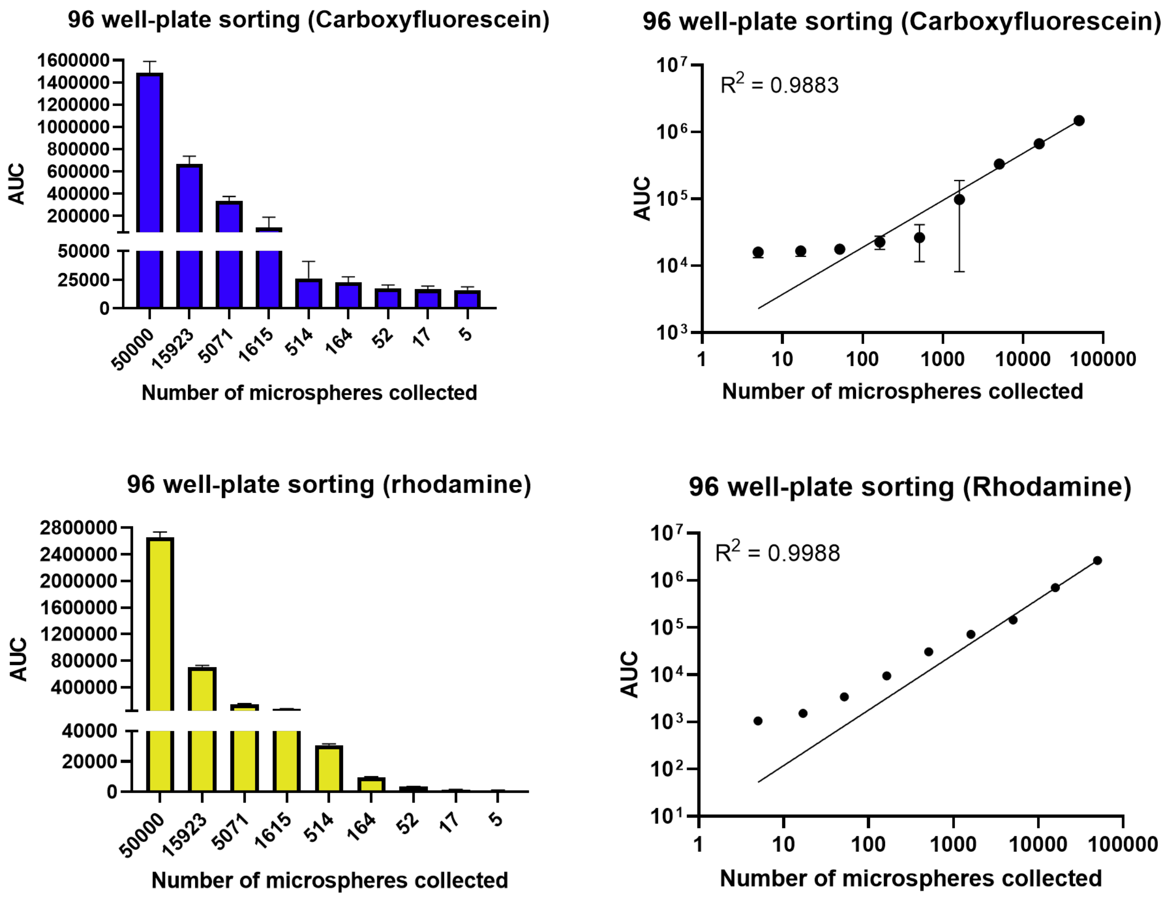

**Figure S 36.** Fluorescent analysis for carboxyfluorescein and rhodamine samples after 96 well plate sort.

#### *4.3 Fluorophore-labelling of proteins*

**Fluorescent labelling of KRAS$^{G12D}$-GMPPnP and KRAS$^{G12D}$-GDP with fluorescein.**

A 2 µM solution of KRAS$^{G12D}$-GMPPnP (20 µL) was incubated with a 2 µM solution of streptavidin-fluorescein conjugate (80 µL) at room temperature for 1 hour, stirring at 300 rpm. After incubation, a native polyacrylamide gel was run to check the conjugation. A 12% polyacrylamide gel (with 4% stacking gel) was run for 90 minutes at 150 V using a tris-glycine running buffer (see Buffers for exact concentrations used). 10 µL of each conjugated sample was added to wells and compared with 10 µL of 2 µM untagged protein and 10 µL of 2 µM streptavidin-fluorescein. The gel was scanned for fluorescence and stained with a Pierce™ Silver Staining Kit (Figure S 37). Fluorescent images were captured using GeneSys Image Capture Software (version 1.5.2.0), using a fluorescein filter (Blue LED Module Filter 525), and stained images were taken using an Epsom scanner.

KRASG12D-GDP was conjugated, checked using a polyacrylamide gel, and stained using the same protocol (Figure S 37).

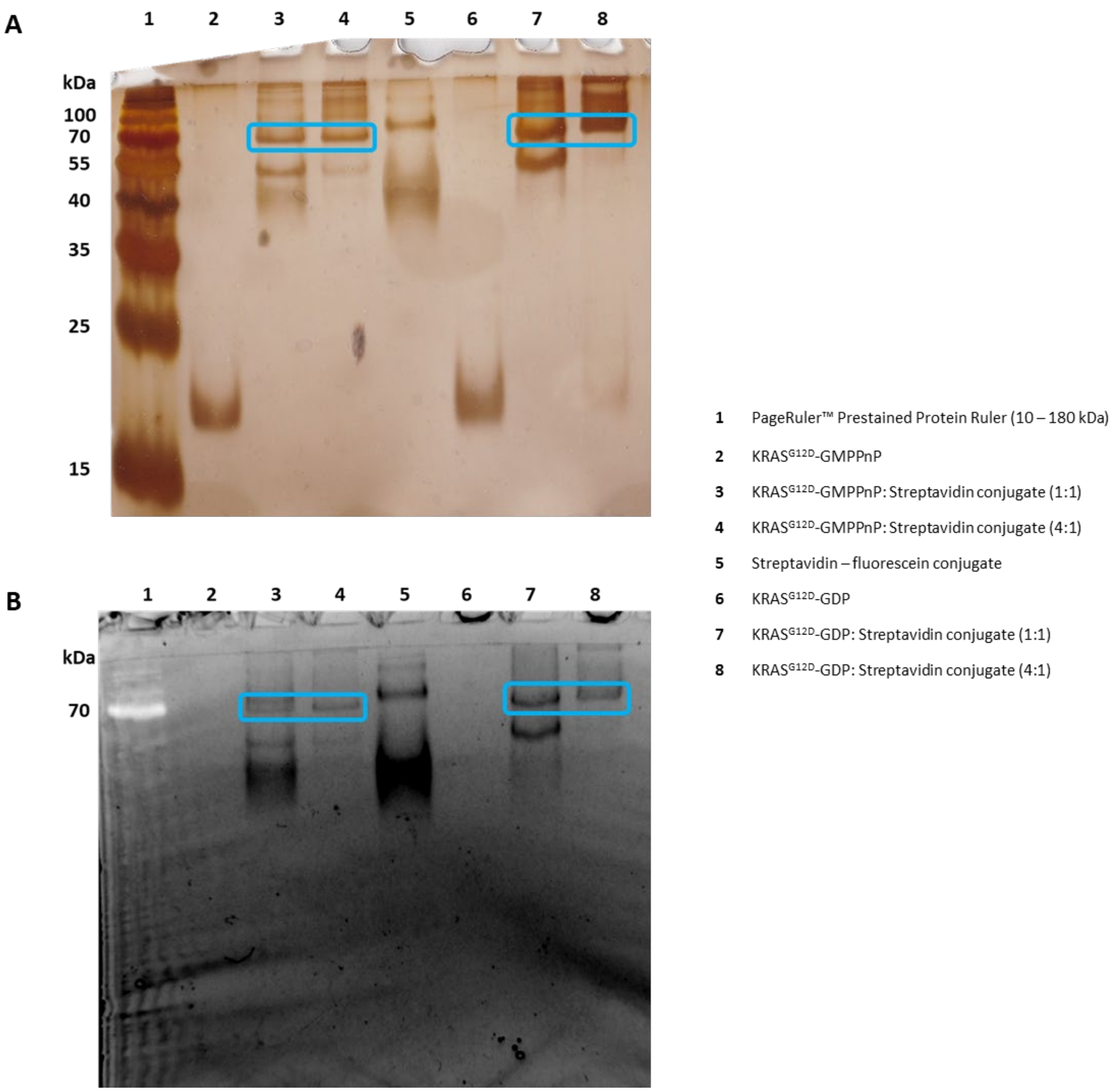

**Figure S 37.** Polyacrylamide gel showing the fluorescent tagging of KRAS$^{G12D}$-GMPPnP and KRAS$^{G12D}$-GDP. A) Silver-stained gel (12%) with blue bands highlighting the presence of the conjugate. B) Fluorescent Image of the same gel with blue bands highlighting the presence of the fluorescent conjugate. The molecular weight ladder, KRAS$^{G12D}$-GMPPnP and KRAS$^{G12D}$-GDP are not fluorescent and therefore are not seen in this gel.

#### Fluorescent labelling of RAF1-RBD with rhodamine.

A 2 µM solution of RAF1-RBD (20 µL) was incubated with a 2 µM solution of streptavidin-rhodamine conjugate (80 µL) at room temperature for 1 hour, stirring at 300 rpm. After incubation, a native polyacrylamide gel was run to check the conjugation. A 12% polyacrylamide gel (with 4% stacking gel) was run for 90 minutes at 150 V using a tris-glycine SDS running buffer (see Buffers for exact concentrations used). 10 µL of each conjugated sample was added to wells and compared with 10 µL of 2 µM untagged protein and 10 µL of 2 µM streptavidin-rhodamine. The gel was scanned for fluorescence and stained with a Pierce™ Silver Staining Kit (Figure S 38). Fluorescent images were captured using GeneSys Image Capture Software (version 1.5.2.0), using a rhodamine filter (Green LED Module Filter 605), and stained images were taken using an Epsom scanner.

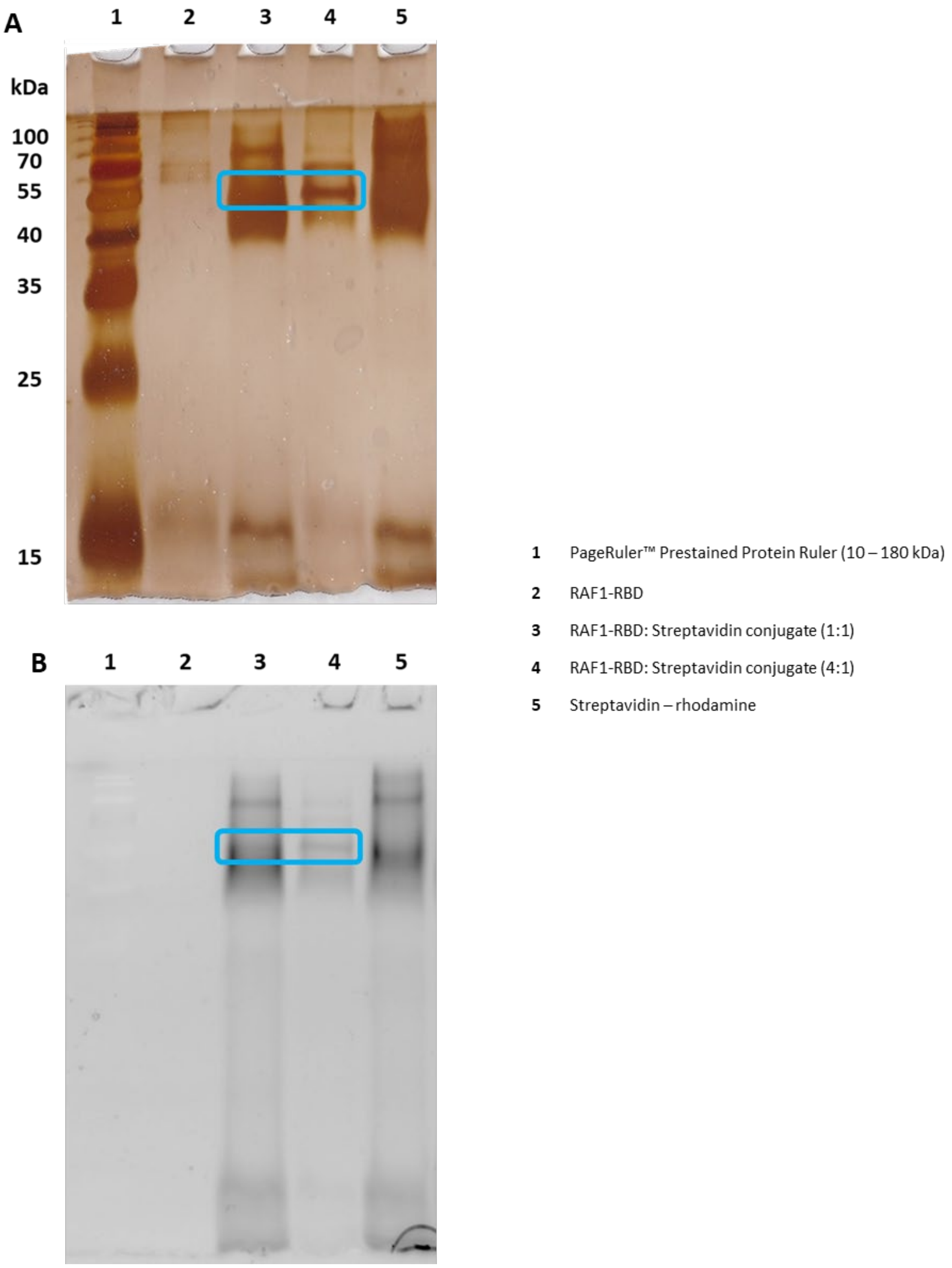

**Figure S 38.** Polyacrylamide gel showing the fluorescent tagging of RAF1-RBD. A) Silver-stained gel (12%) with blue bands highlighting the presence of the conjugate. B) Fluorescent Image of the same gel with blue bands highlighting the presence of the fluorescent conjugate. The molecular weight ladder and RAF1-RBD are not fluorescent and therefore are not seen in this gel.

#### *4.4 Checking for non-specific binding*

**FACS analysis for non-specific binding of oligomer library with fluorescent tags**

The FACS was calibrated for laser analysis and two-way sorting. A small subset of the oligomer library (2 mg) was removed and analysed using both the 488 nm and 561 nm lasers. Then the sample was incubated with 2 µM streptavidin-fluorescein tag (2 mL) at room temperature for 2 hours. After incubation the mixture was centrifuged, and the supernatant was removed to be analysed separately and the remaining library microspheres were suspended in sheath fluid (5 mL). The suspended microspheres and the supernatant were analysed using both the 488 nm and 561 nm lasers.

This process was repeated using the streptavidin-rhodamine conjugate.

**FACS analysis for non-specific binding of -OH modified TentaGel® microspheres with fluorescent tags**

The FACS was calibrated for laser analysis and two-way sorting. -OH Modified TentaGel® microspheres (20 mg) were analysed using both the 488 nm and 561 nm lasers to set limit for where autofluorescence might be expected (Figure S 39). On this basis, anything higher than $10^3$ AU would be considered a positive result.

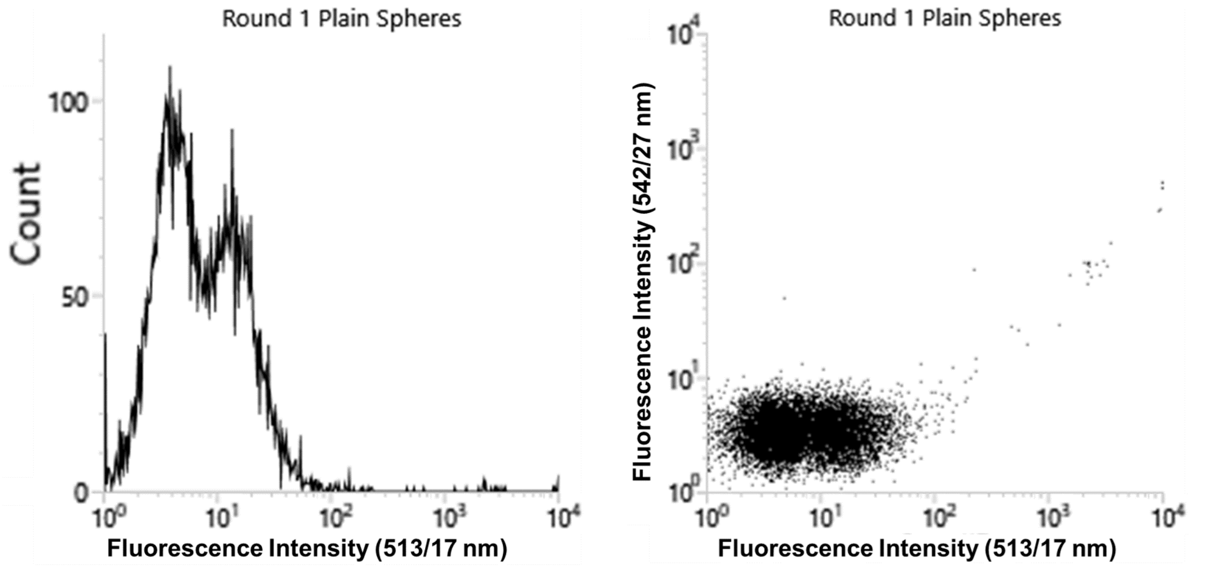

**Figure S 39.** FABS analysis of plain TentaGel® beads.

Then the sample was incubated with 2 µM streptavidin-fluorescein tag (2 mL) at room temperature for 2 hours. After incubation the mixture was centrifuged, and the supernatant was removed to be analysed separately and the remaining library microspheres were suspended in sheath fluid (5 mL). The suspended microspheres and the supernatant were analysed using both the 488 nm and 561 nm lasers.

This process was repeated using the streptavidin-rhodamine conjugate, and the 4:1 conjugates of $KRAS^{G12D}$-GMPPnP, $KRAS^{G12D}$-GDP and RAF1-RBD.

#### *4.5 Selection of phosphoestamers for selective PPI inhibition*

*Round 1 – positive selection for binding of $KRAS^{G12D}$-GMPPnP*

A 4:1 solution of $KRAS^{G12D}$-GMPPnP – fluorescein solution (2 µM) was prepared. 5 mL of this conjugated solution was added to the oligomer library (swelled in 110 mL of KRAS buffer). This mixture was shaken at room temperature at 400 rpm for 4 hours. After this the library solution was centrifuged, and the supernatant was removed. The remaining solid was dispersed in sheath fluid (50 mL).

The oligomer library (0.148 g) was dispersed in sheath fluid. The FACS lasers were calibrated, and a two-way calibration was set up. The 488 nm laser and bandpass filters were used to set the collection gates in the FACS and collect the microspheres with the highest fluorescent signal at 513/17 nm and 542/27 nm, giving 3,900,197 (1.76%) beads (Figure S 40). This process was repeated, with a narrower gate applied, giving 48,169 microspheres (Figure S 41) which were collected for the next Round.

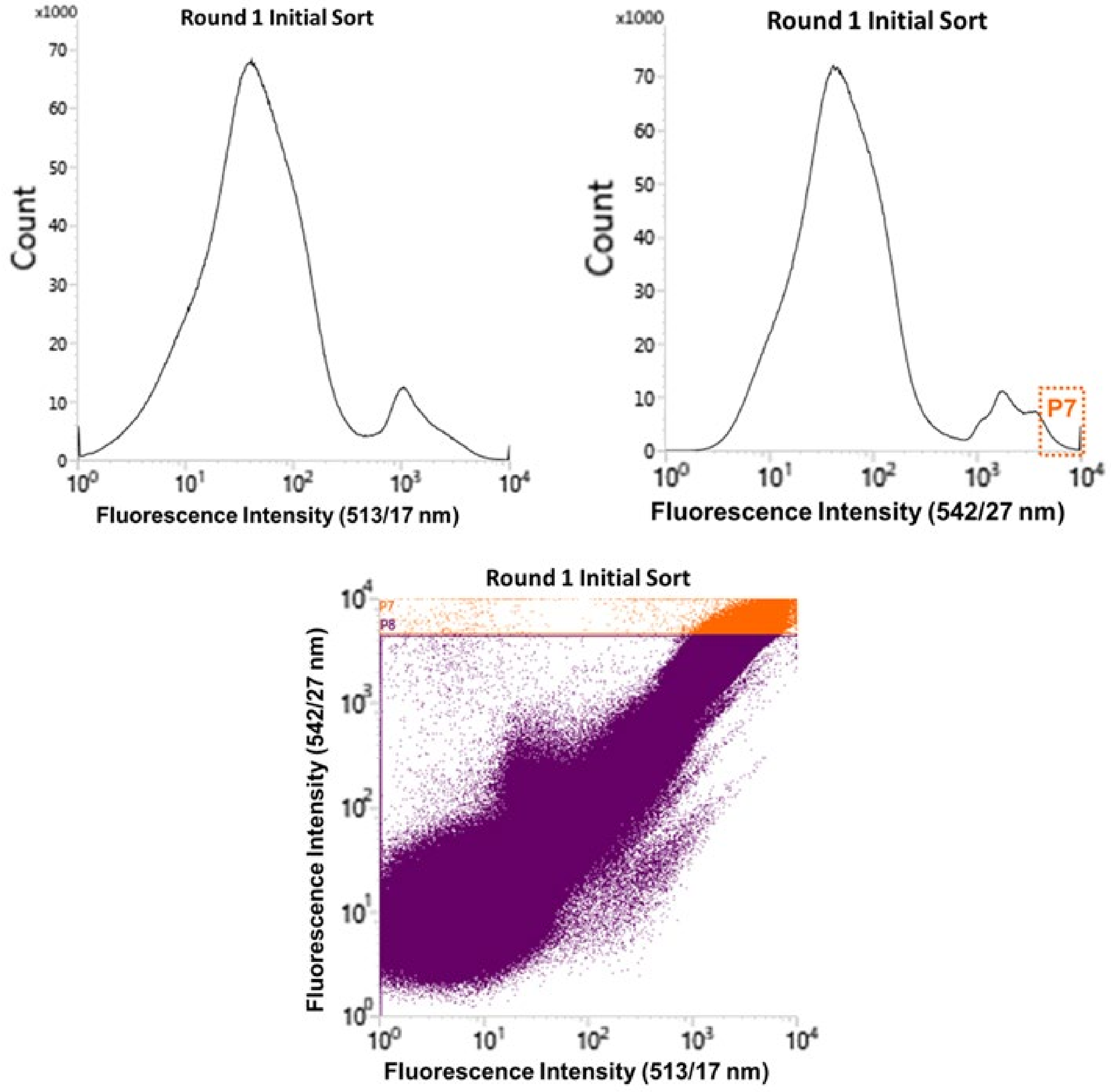

**Figure S 40.** FABS data from initial Round 1 library sorting. "P7" refers to the selection gate applied.

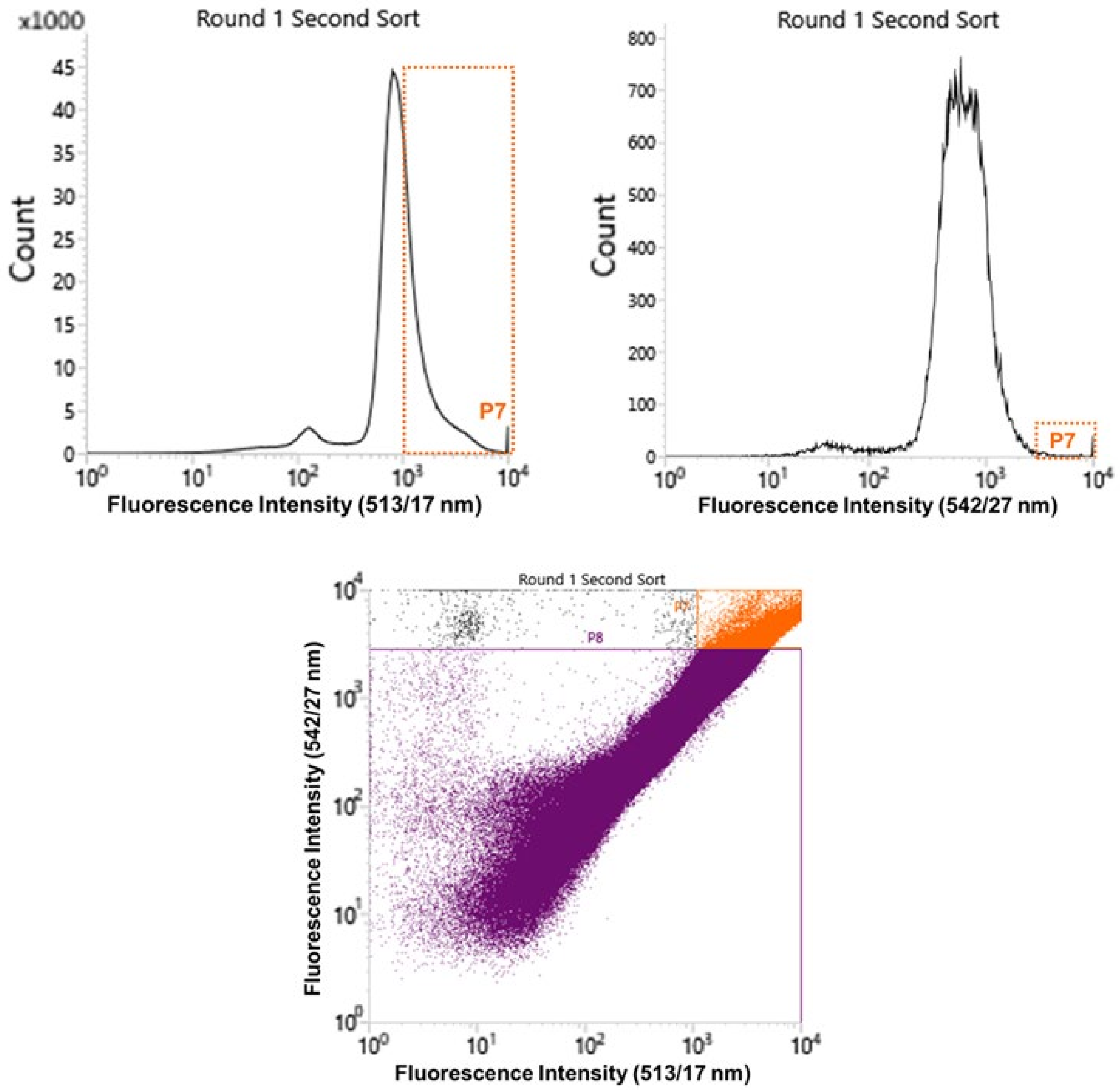

**Figure S 41.** FABS data from second Round 1 library sorting. "P7" refers to the selection gate applied.

*Removal of protein from the oligomer library after each selection round.*

After each selection round, the remaining microspheres were centrifuged at 300 rpm and the supernatant was removed. 2 mL of 0.2 M sodium hydroxide was added to the library and the mixture was stirred for 2 hours. The mixture was centrifuged at 300 rpm and the supernatant was removed. The FACS was calibrated for laser analysis and two-way sorting. A small subset was dispersed in 2 mL sheath fluid and analysed on the FACS using both the 488 and 561 nm lasers and bandpass filters to ensure there was no remaining fluorescent signal. This subset was collected using a two-way sorting calibration, returned to the main library and centrifuged, with the supernatant being removed.

*Round 2 – negative selection against binding of KRAS$^{G12D}$-GDP*

After Round 1 of FACS selection the KRAS$^{G12D}$-GMPPnP was removed with a NaOH wash. After this the microspheres were dispersed in RAS Buffer (2 mL) and 2 mL of 2 µM fluorescein-KRAS$^{G12D}$-GDP was added. The solution was stirred for one hour and then centrifuged at 300 rpm for 5 minutes. The supernatant was removed, and the library was resuspended in sheath fluid (3 mL). The FACS lasers were calibrated, and a two-way calibration was set up. The 488 nm laser and bandpass filters were used to set the collection gates in the FACS and collect the microspheres with the lowest fluorescent signal at 513/17 nm and 542/27 nm (Figure S 42). From this selection round, 12,111 microspheres were collected for the next step.

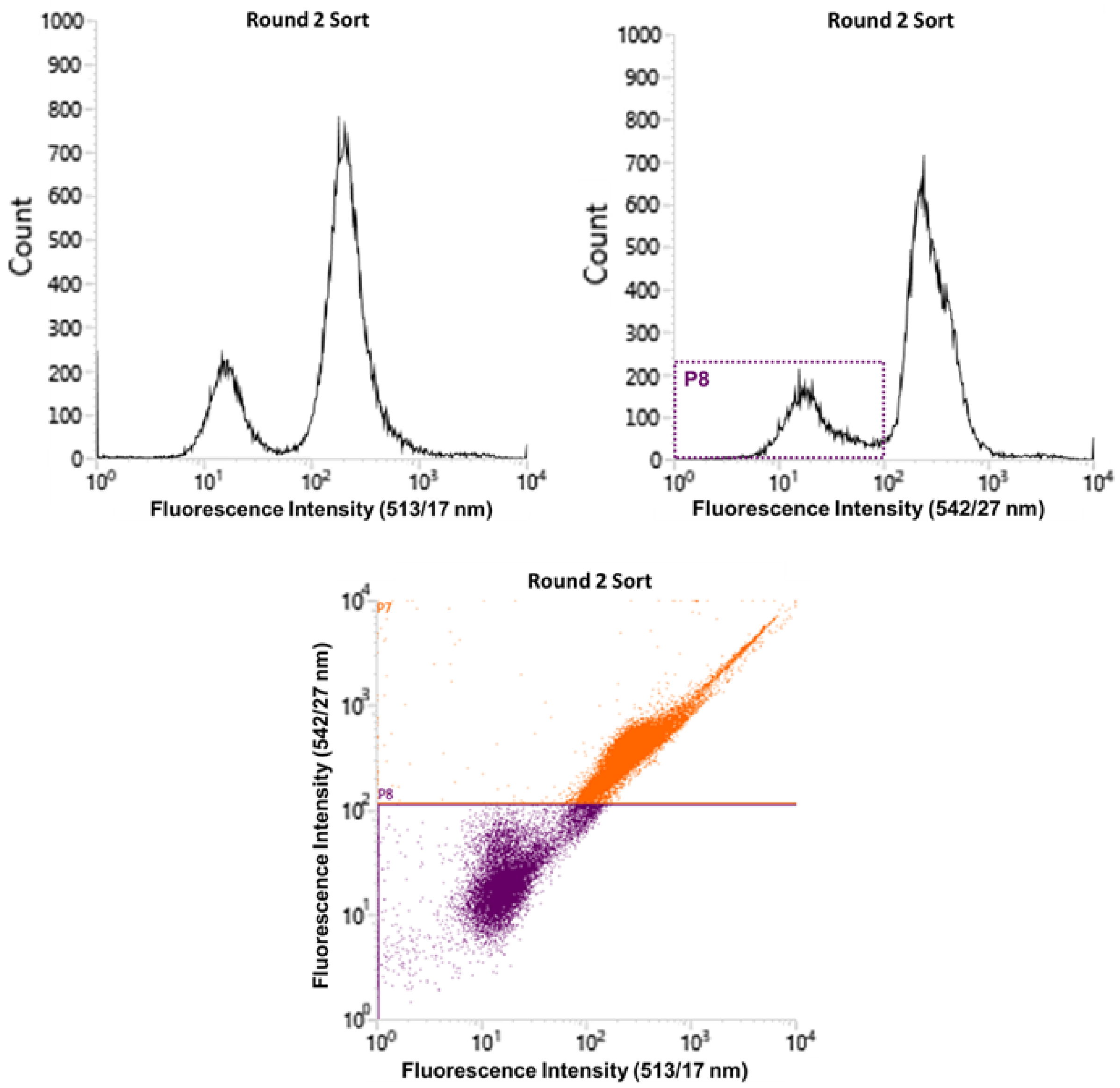

**Figure S 42.** FABS data from second Round 2 library sorting. “P8” refers to the selection gate applied.

*Round 3 – selection for blockage of KRAS$^{G12D}$-GMPPnP/RAS1-RBD interaction*

After Round 2 of FACS selection the KRAS$^{G12D}$-GDP was removed with a NaOH wash. The microspheres were dispersed in RAF Buffer (2 mL); 495 µL of 2 µM fluorescein-KRAS$^{G12D}$-GMPPnP and 1485 µL of rhodamine RAF1-RBD was added. This is enough KRAS to bind to 5,095 microspheres (42% of the remaining library), and the RAF would cover 15,286 spheres which was more than the entirety of the remaining library. If there were no oligomers within the library that disrupted the KRAS$^{G12D}$-RAF1-RBD interaction, then the FACS would show a strong rhodamine signal within the analysis. However, if the oligomers strongly bind to the KRAS$^{G12D}$ and stop any interaction with the RAF1-RBD then the protein would be removed after centrifuging and there would be seen a strong fluorescein signal (as well as microspheres with little to no rhodamine fluorescent signal). The solution was stirred for one hour and then centrifuged at 300 rpm for 5 minutes. The supernatant was removed, and the library was resuspended in sheath fluid (3 mL). The FACS lasers were calibrated, and a two-way calibration was set up. The 561 nm laser and bandpass filters were used to set the collection gates in the FACS and collect the microspheres with the lowest fluorescent signal at 585/29 nm and 600 nm (Figure S 43). Looking at the 488 nm laser there was a spread of data, and within the P7 gate there were 1,713 microspheres with a strong fluorescein signal (Figure S 43). When highlighting the P7 data using the 561 nm filters, it can be seen that the majority of these microspheres with a strong fluorescein signal do not have a strong rhodamine signal, consistent with disruption of the protein interaction (Figure S 44). From this selection round, 676 microspheres were collected for the final selection round.

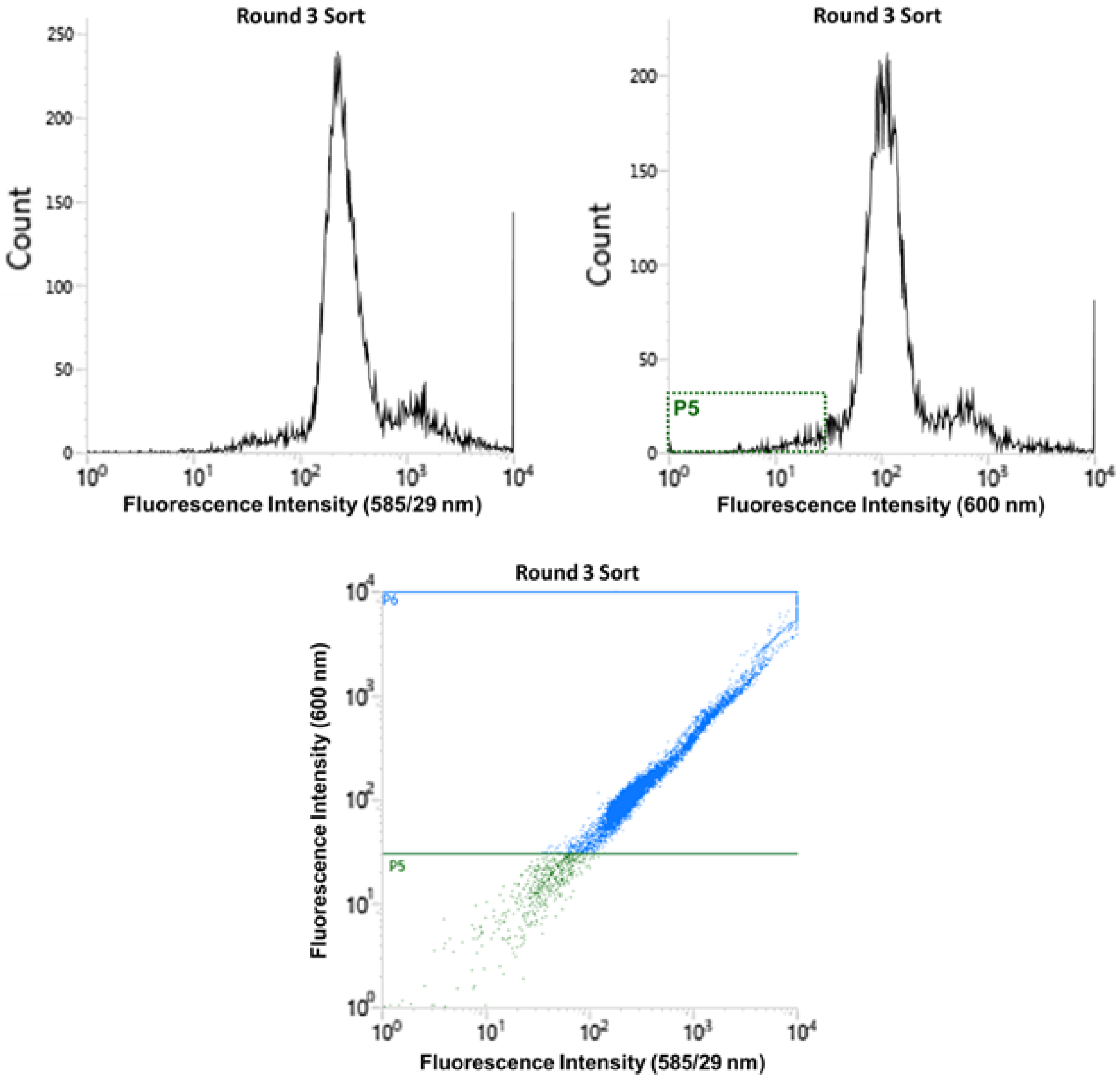

**Figure S 43.** FABS data from second Round 3 library sorting. “P5” refers to the selection gate applied to collect beads for the subsequent round.

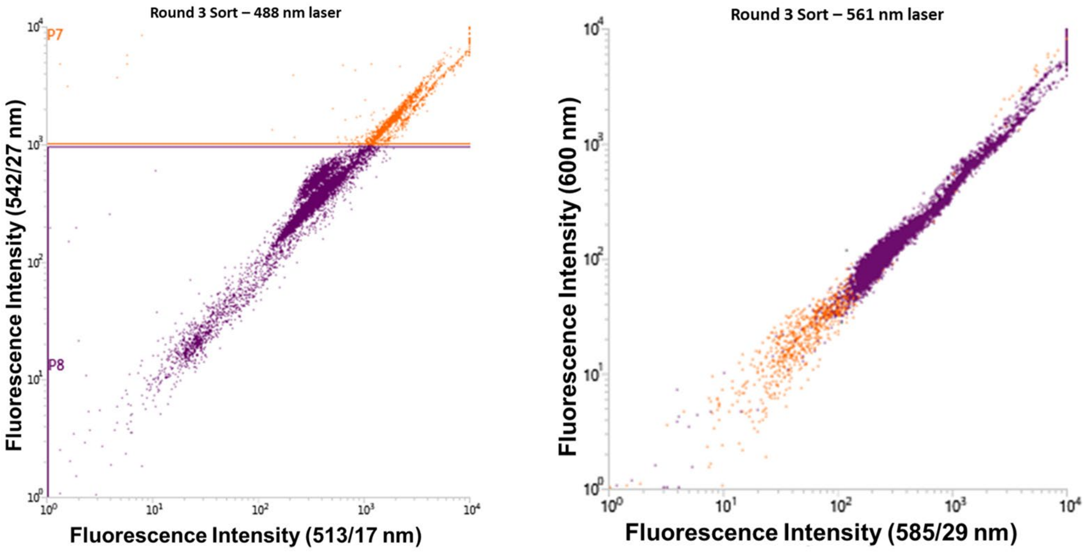

**Round 3 Sort**

| Populations | Events | % Total | % Parent |
|---|---|---|---|
| All Events | 12,311 | 100.00% | #### |
| P7 | 1,713 | 13.91% | 13.91% |
| P8 | 10,598 | 86.09% | 86.09% |

**Figure S 44.** FABS data from Round 3 library sorting. Left: Data from the 488 nm laser, with the population in each gate (used here for analysis rather than collection) below. Right: Data from the 561 nm laser highlighting the 488 nm gates (P7 in orange, P8 in purple).

##### *Round 4 – positive selection for strongest KRAS$^{G12D}$-GMPPnP binders*

After Round 3 of FACS selection the proteins were removed with a NaOH wash. The remaining library microspheres were dispersed in RAS Buffer (2 mL); 51 µL of 2 µM fluorescein-KRAS$^{G12D}$-GMPPnP was added. The solution was stirred for one hour and then centrifuged at 300 rpm for 5 minutes. The supernatant was removed, and the library was resuspended in sheath fluid (3 mL). The FACS lasers were calibrated, and a 96 well plate calibration was set up. The 488 nm laser and bandpass filters were used to set the collection gates in the FACS and collect the microspheres with the highest fluorescent signal at 513/17 nm and 542/27 nm. In this selection round, only the top 200 microspheres with the highest florescent signal were collected, with one TentaGel® microsphere sorted into 1 well of the well plate containing 20 µL of RAS buffer (Figure S 45).

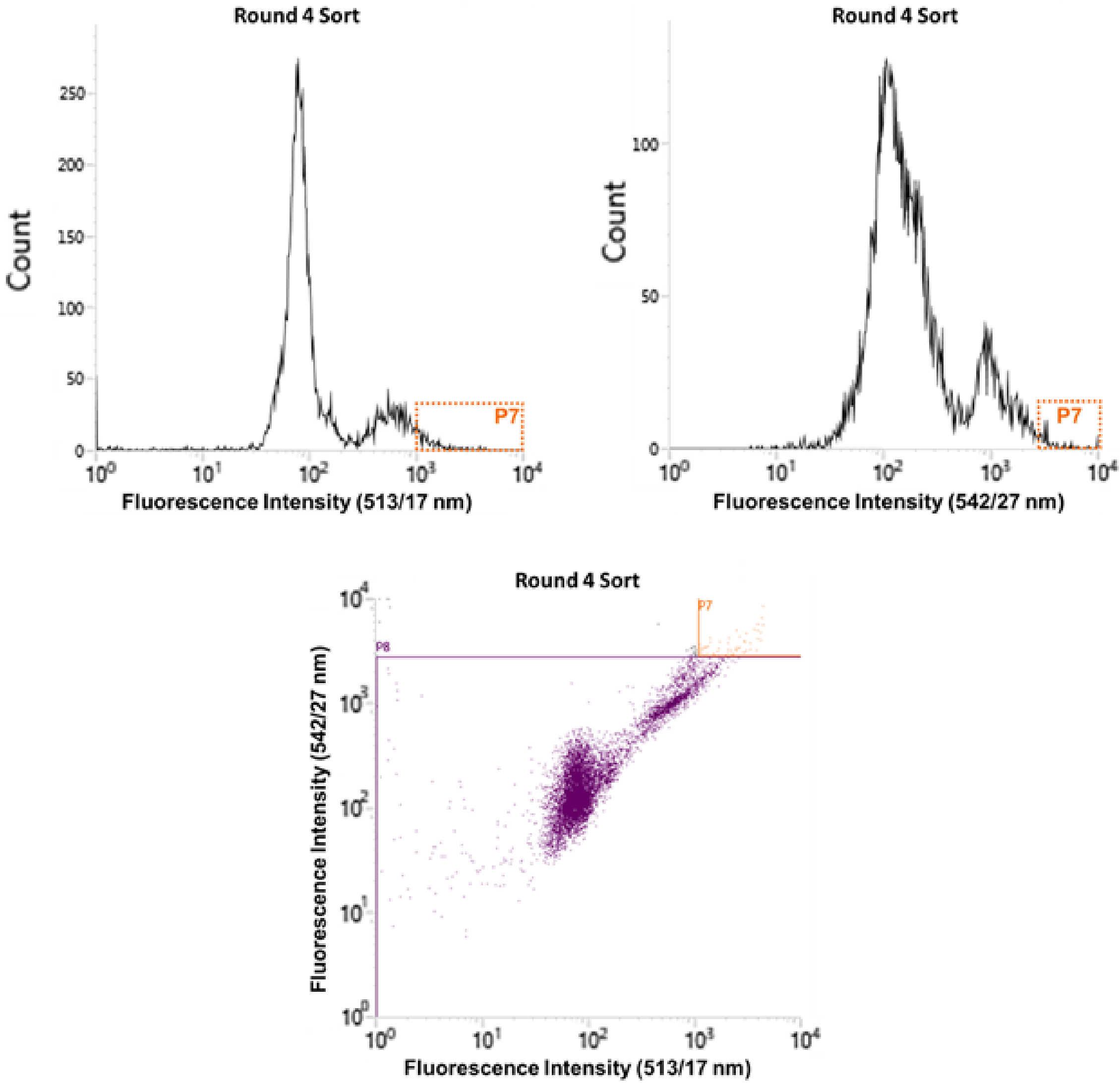

**Figure S 45.** FABS data from second Round 4 library sorting. "P8" refers to the selection gate applied to collect beads for the individual wells and MSMS sequencing.

### 5 Sequencing by mass spectrometry

#### 5.1 Preparation of samples for mass spectrometry

The 200 oligomers collected from the fluorescent selection were placed under a UV lamp and irradiated at 365 nm for 4 hours. After this the contents of each well were transferred into individual PCR tubes and stored at 4 °C.

To prepare the oligomers for MS/MS analysis, the oligomers were desalted using Pierce™ C18 Spin Tips and Columns following the manufacturers protocol.[1] Briefly, the C18 tips were wetted with 20 µL of Milli Q® water containing 0.1% TFA, centrifuging at 1000 x g for 1 minute. The C18 tips were then equilibrated with 20 µL of Milli Q® water containing 0.1% TFA, centrifuging at 1000 x g for 1 minute. The sample (between 20 and 50 µL) was then added to the C18 tips and centrifuged at 1000 x g for 1 minute. The tip was washed by adding 20 µL of Milli Q® water containing 0.1% TFA, centrifuged at 1000 x g for 2 x 1 minute. The sample was eluted using 20 µL of acetonitrile containing 0.1% TFA and centrifuging at 1000 x g for 2 x 1 minute. The sample was dried and resuspended in 6 µL of TEAB buffer (98:2 MS buffer A: MS buffer B).

#### 5.2 LC-MS/MS analysis of oligomer hits and DNA standards

Oligomers were randomly selected and desalted using the C18 spin tip procedure. Following this, oligomers and the DNA standards were analysed using an Acquity UHPLC system and a Bruker micrOTOF-Q II mass spectrometer in negative ion mode. 4 µL of oligomer was injected into the UHPLC system and held in the trap column (nanoEase™ *m/z* symmetry C18, 180 µM x 20 mm column) for 4.5 minutes using a flow rate of 10 µL at 98% Buffer A and 2% Buffer B (see 4.2.2.1). After trapping the samples go through the UHPLC column (nanoEase™ *m/z* HSS C18T3, 100 Å, 1.8 µM, 75 µM x 150 mm column) at a rate of 0.3 µL/min. The solvent gradient was as shown in Table S 6.

**Table S 6**. Buffer gradient used in LC MS/MS.

| Buffer A (%) | Buffer B (%) | Time (min) |
|---|---|---|
| 98 | 2 | 0 |
| 98 | 2 | 1 |
| 50 | 50 | 30 |
| 10 | 90 | 31 |
| 10 | 90 | 32 |
| 98 | 2 | 33 |
| 98 | 2 | 60 (end) |

The mass spectrometry method collects MS data from 10 to 50 minutes in the run between an *m/z* range of 100 to 3000 Da, with an individual scan time of 1 second. When an individual MS scan had a total ion current (TIC) intensity above 1000, the system switched to MS/MS analysis and the ion data from the 5 most intense MS/MS ions was collected, with an MS/MS scan rate of 0.5 seconds. Throughout the analysis, leucine enkephalin was used as a lock mass standard and analysed every 30 seconds, where drift time and mass data was calibrated accordingly. A buffer blank (4 µL at 98/2 Buffer A/B) was run before any samples and after every sample (oligomers and DNA standards) to avoid potential carryover. After this data was converted and analysed using RoboOligo software.[2]

The DNA standard was made at concentrations of 10 µM, 5 µM, 1 µM, 0.5 µM, 0.2 µM, 0.1 µM, 0.05 µM, 0.02 µM and 0.01 µM. The data was used to determine the limit of detection, and fragmentation data was compared to RoboOligo analysis to set up a methodology for analysis of unknowns.

Data collected off the mass spectrometer was MassLynx .Raw files. To use the RoboOligo software, the data had to first be converted into mascot generic format (mgf) files. ProteoWizard is a software project that provides open-source software for proteomics analysis.[3,4] ProteoWizard MSConvert is software used for file conversion and combines spectra with the same parent ion together using the PASEF MGF program. MSConvert was used to convert all MassLynx files into mgf files.

When identifying the sequence of the unknowns, the same four-step system was used for each oligomer (Figure S 46).

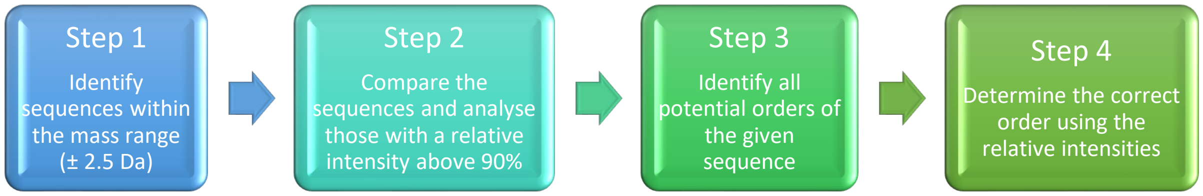

**Figure S 46.** Oligomer sequencing workflow.

Acceptable MS samples produced a clearly defined $[M-2H]^{2-}$ peak which was used to calculate a molecular mass. This mass was used to narrow down the selection of potential sequences and every sequence was checked and compared using RoboOligo. The software compares the combined intensity of the signal across the spectra and calculates the relative intensity, with the highest intensity value being given 100% and all other sequences compared to this data. For example, a sequence that does not fit the data very well will be given a much lower relative intensity (such as 40%) and so this is not the correct sequence. Comparing all the different iterations and different orders of iterations created a table of all the relative intensities. The sequences with an intensity value above 90% were analysed and selected for the next stage of sequencing. In every instance the sequences within this range consisted of the same monomers in a different order, so the correct order needed to be determined. Every order of one sequence was then analysed and the relative intensities compared again. The sequence with the highest relative intensity, meaning the most data fits the given spectra, was determined to be the correct monomer composition in the correct sequence. However, due to a large number of highly ranked sequences, resynthesis and validation was required to assess rectitude of MS/MS sequencing.

After the photocleavage process to remove the solid support beads from the oligomers in the library, the oligomers that remained for analysis contained an OH group at one end of the molecule and a phosphate group at the other (denoted with P) (Figure S 47). This introduces asymmetry into the oligomers and means any sequences that should be identical due to the symmetry of the monomers have these two different groups at either end that will affect their fragmentation patterns when analysing the MS/MS data.

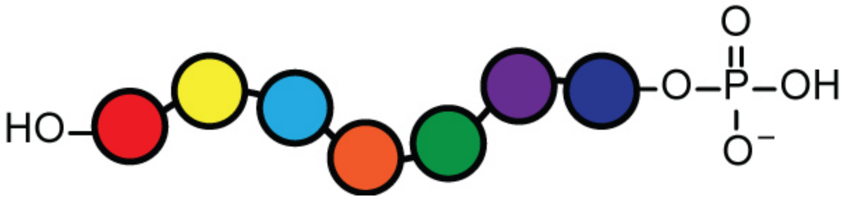

**Figure S 47.** General oligomer sequence after photocleavage.

Full methodological discussion is given for oligomer 1, **O1**. The data for the other oligomers should be interpreted accordingly.

**Oligomer 1**

**O1** showed a doubly charged peak at 854.978 *m/z*, meaning the monoisotopic mass would be 1711.956 Da and a $[M\text{-}H]^-$ peak at 1710.963 Da (Figure S 48), with the intensity of the $[M\text{-}H]^-$ across the peak being above the limit of 1000, so this precursor underwent MS2 analysis. This gave the advantage of comparing two sets of RoboOligo data, which is better for determining the sequence.

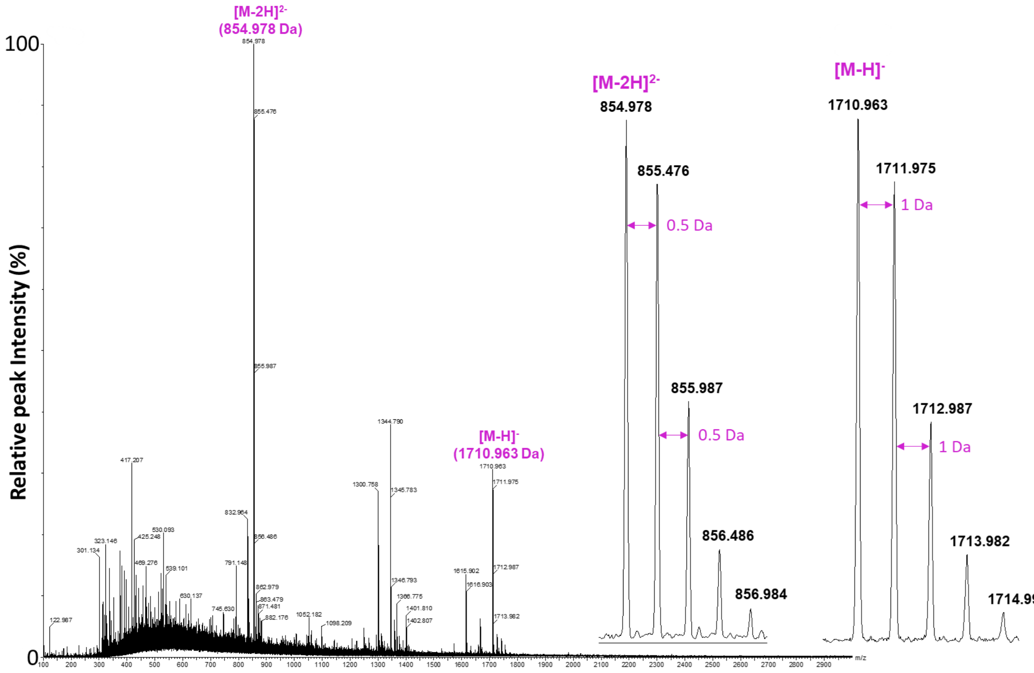

**Figure S 48.** MS1 data from oligomer **O1** with both $[M\text{-}H]^-$ and $[M\text{-}2H]^{2-}$ charge states highlighted.

Table S 7 shows all the potential monomer combinations ± 2.5 Da of 1711 Da. Step 1 of sequencing involved using the same spectra and analysing these different sequences (and 5 different sequences using the same composition) and comparing the relative intensities.

**Table S 7.** Composition possibilities within the mass range of **O1**.

| Mass | | Monomer Combinations (OH → P) | | | | |
|---|---|---|---|---|---|---|
| **1711.956** | 1 | BPA | C12 | DAN | DAN | HEG |
| | 2 | C12 | C12 | cYY | cYY | cYY |
| | 3 | C12 | DAN | DAN | DAN | cSS |
| | 4 | C12 | DAN | HEG | HEG | HEG |

As seen in Table S 1, there were no 7-mer sequences that fit in the mass range ± 2.5 Da, so while the molecular mass is within the library range, the oligomer itself is a truncated version. To determine which sequence was the correct one using RoboOligo analysis, 20 different sequences (5 combinations of the 4 sequences within the range, Table S 8) were randomly selected for comparison.

**Table S 8.** Initial sequences selected for RoboOligo analysis for O1.

| **Sequence Number** | **Sequence (OH → P)** |
|---|---|
| **1** | HEG-DAN-DAN-C12-BPA |
| **2** | DAN-HEG-BPA-DAN-C12 |
| **3** | DAN-C12-HEG-BPA-DAN |
| **4** | C12-DAN-BPA-HEG-DAN |
| **5** | BPA-C12-DAN-DAN-HEG |
| **6** | cYY-cYY-cYY-C12-C12 |
| **7** | cYY-C12-cYY-C12-cYY |
| **8** | C12-cYY-C12-cYY-cYY |
| **9** | C12-cYY-cYY-cYY-C12 |
| **10** | C12-C12-cYY-cYY-cYY |
| **11** | cSS-DAN-DAN-DAN-C12 |
| **12** | DAN-cSS-DAN-C12-DAN |
| **13** | DAN-DAN-C12-DAN-cSS |
| **14** | DAN-cSS-DAN-DAN-C12 |
| **15** | C12-DAN-DAN-DAN-cSS |
| **16** | HEG-HEG-HEG-DAN-C12 |
| **17** | HEG-DAN-HEG-C12-HEG |
| **18** | C12-HEG-HEG-HEG-DAN |
| **19** | DAN-HEG-HEG-C12-HEG |
| **20** | C12-DAN-HEG-HEG-HEG |

There were 2 different combined spectra to analyse for the *m/z* of 854.978 and 1 combined data spectrum to analyse at 1710.963 *m/z*. Figure S 49 shows an example of the different sequences and their ions detected in RoboOligo using one of the combined spectra with 854.978 *m/z* as a precursor, and the comparative relative intensities from this data.

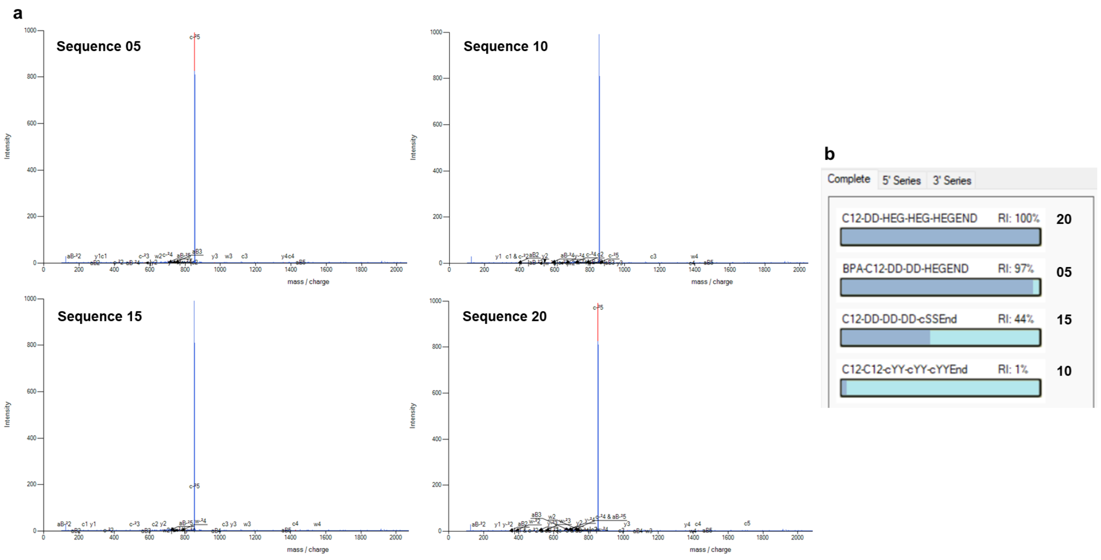

**Figure S 49.** (a) RoboOligo analysis of four selected sequences and (b) their relative intensities. DD = DAN

Combining the intensities across all of RoboOligo data allowed for the calculation of an overall relative intensity, shown in Figure S 50. This data shows the last sequence set (16 – 20) better fits all the data compared to the other three, so the next step was to determine the correct order of the sequences.

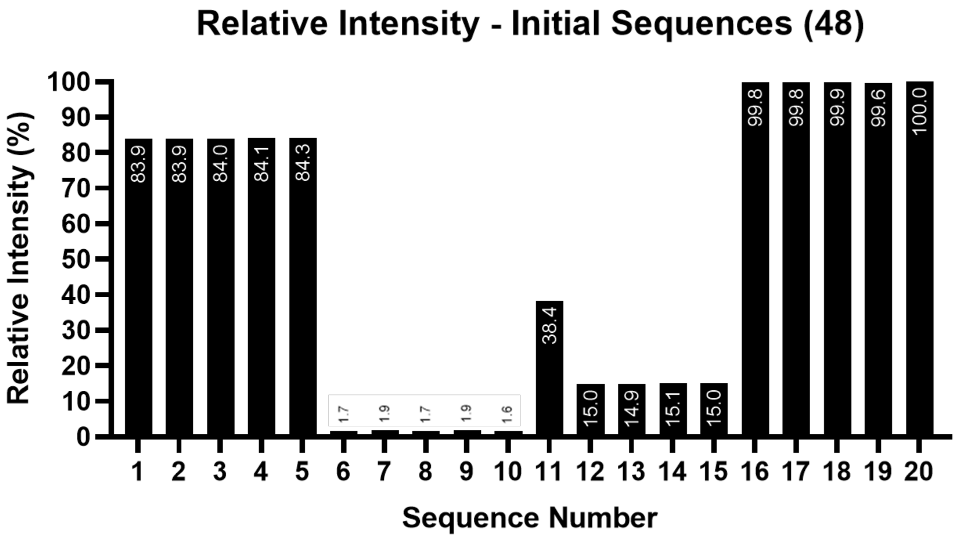

**Figure S 50.** Relative intensity comparisons of initial sequences for **O1**.

The sequence with the highest relative intensity contained one **C12** monomer, one **DAN** monomer and three **HEG** monomers. There were 20 combination possibilities using a 1-1-3 combination of monomers, and these are laid out in Table S 9, with 16 – 20 being the sequences already analysed.

**Table S 9.** Possible sequences for known composition of **O1**.

| Sequence Number | Sequence (OH → P) |
|---|---|
| **16** | HEG-HEG-HEG-DAN-C12 |
| **17** | HEG-DAN-HEG-C12-HEG |
| **18** | C12-HEG-HEG-HEG-DAN |
| **19** | DAN-HEG-HEG-C12-HEG |
| **20** | C12-DAN-HEG-HEG-HEG |
| **21** | C12-HEG-DAN-HEG-HEG |
| **22** | C12-HEG-HEG-DAN-HEG |
| **23** | DAN-HEG-HEG-HEG-C12 |
| **24** | DAN-C12-HEG-HEG-HEG |
| **25** | DAN-HEG-C12-HEG-HEG |
| **26** | HEG-C12-HEG-HEG-DAN |
| **27** | HEG-C12-DAN-HEG-HEG |
| **28** | HEG-DAN-C12-HEG-HEG |
| **29** | HEG-C12-HEG-DAN-HEG |
| **30** | HEG-DAN-HEG-HEG-C12 |
| **31** | HEG-HEG-C12-DAN-HEG |
| **32** | HEG-HEG-C12-HEG-DAN |
| **33** | HEG-HEG-DAN-C12-HEG |
| **34** | HEG-HEG-DAN-HEG-C12 |
| **35** | HEG-HEG-HEG-C12-DAN |

Comparing the different combinations in Table S 9, the initial sequences with the highest intensities were 20, 18, and 16 respectively. These three sequences all contained three **HEG** units consecutively compared to 17 and 19 which had the **HEG** units separated, which could suggest this is important in the sequence order. When analysing the different sequence orders they all had a relative intensity above 99%, but sequence 20 had the best fit of all the data (Figure S 51**Error! Reference source not found.**) and so was designated as the working sequence. Its structure is given in

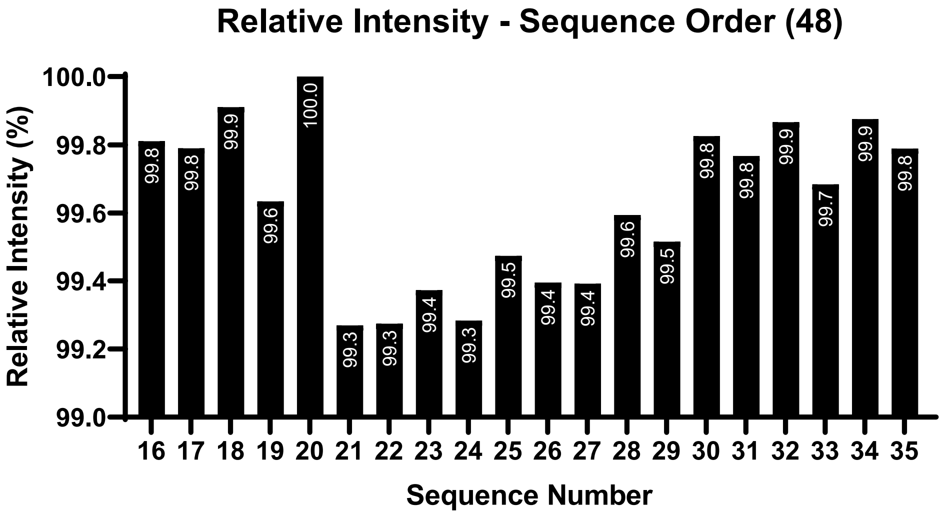

**Figure S 51.** Relative intensities of possible sequence variations for **O1**.

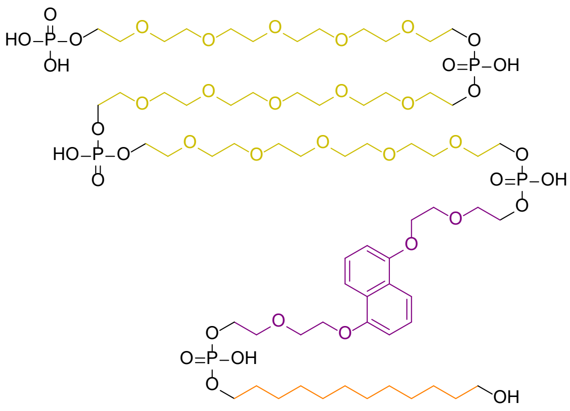

**O1**: HO-C12-DAN-HEG-HEG-HEG-P

**Figure S 52.** Best fit chemical structure of **O1**.

The RoboOligo data from this sequence fits both the $[M-H]^-$ data and the $[M-2H]^{2-}$ data (Figure S 53) and gave an overall relative intensity of 100%. Whilst other sequences did show a relative intensity close to this (above 99%) due to limitations in chemicals and time only this sequence was chosen to analyse further as it was the best fit of both data.

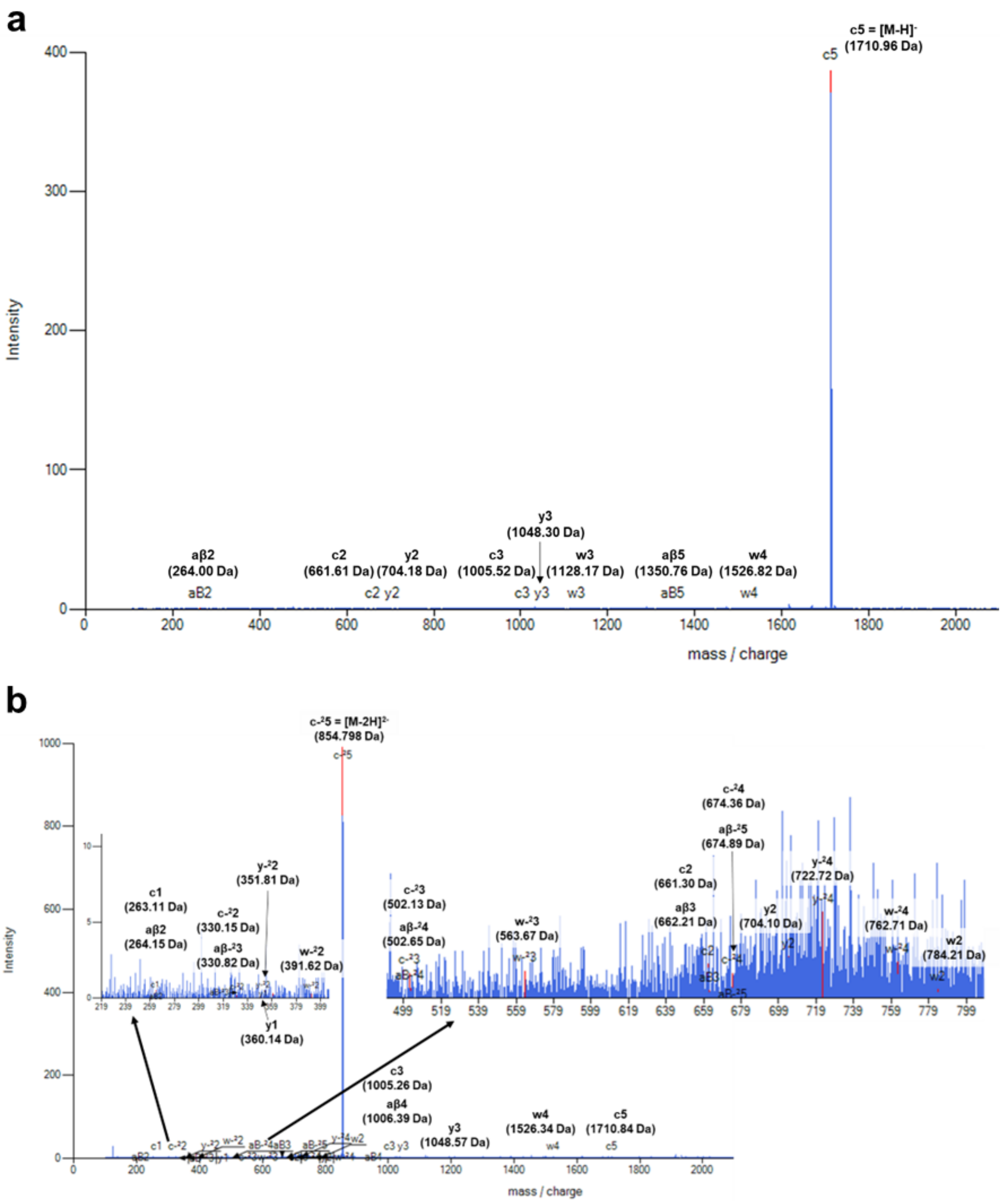

**Figure S 53.** (a) RoboOligo analysis of **O1** using 1710.963 $[M-H]^-$ as a precursor. (b) RoboOligo analysis of **O2** using 854.978 $[M-2H]^{2-}$ as a precursor.

##### Oligomer 2

The MS1 data revealed the existence of an oligomer with a double charge state at 992.314 Da (Figure S 54), meaning the overall monoisotopic mass would be 1986.628 Da.

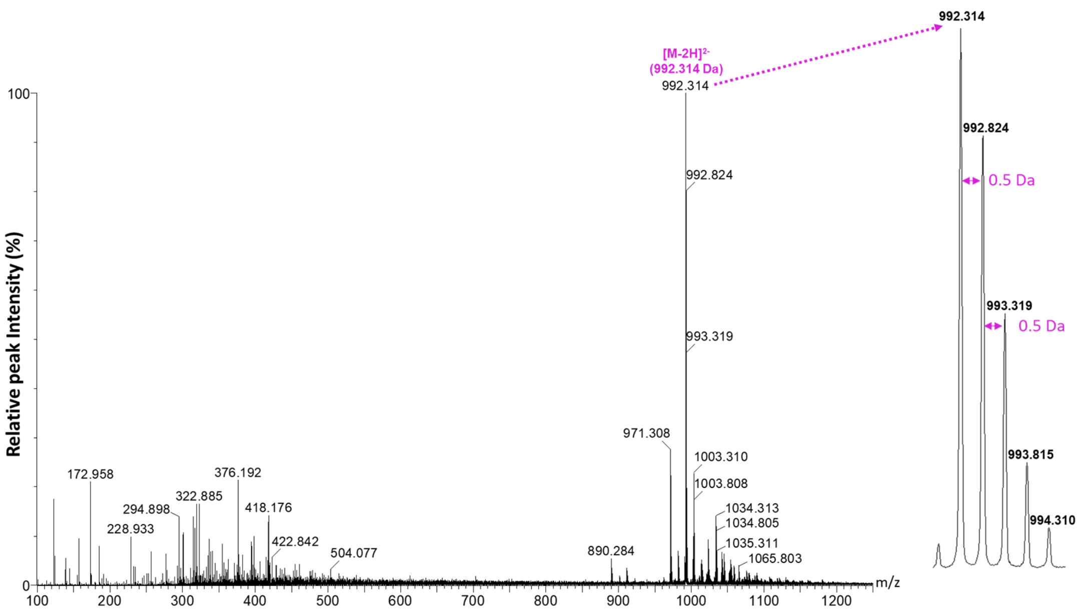

**Figure S 54.** MS1 data of **O2** from the chromatogram at 35 minutes, with $[M-2H]^{2-}$ charge state highlighted.

**Table S 10.** Sequence possibilities within the mass range of **O2**

| **Mass** | | **Monomer Combinations (OH → P)** | | | | | | |
|---|---|---|---|---|---|---|---|---|
| **1986.628** | **1** | BPA | BPA | C12 | C12 | cSS | cSS | cYY |
| | **2** | BPA | BPA | cSS | cSS | cSS | cSS | NDI |
| | **3** | BPA | C12 | C12 | C12 | C12 | cSS | cYY |
| | **4** | BPA | C12 | C12 | cSS | cSS | cSS | NDI |
| | **5** | C12 | C12 | HEG | cSS | cSS | cSS | cYY |
| | **6** | HEG | cSS | cSS | cSS | cSS | cSS | NDI |
| | **7** | BPA | BPA | C12 | cSS | NDI | NDI | - |
| | **8** | BPA | C12 | C12 | C12 | NDI | NDI | - |
| | **9** | C12 | C12 | C12 | HEG | cYY | NDI | - |
| | **10** | C12 | HEG | cSS | cSS | NDI | NDI | - |
| | **11** | DAN | DAN | DAN | cYY | cYY | - | - |

**Table S 11.** Initial sequences selected for RoboOligo analysis for **O2**.

| Sequence Number | Sequence (OH → P) | Sequence Number | Sequence (OH → P) |
|---|---|---|---|
| **1** | HEG-cSS-cSS-NDI-cSS-cSS-cSS | **21** | NDI-NDI-cSS-C12-BPA-BPA |
| **2** | NDI-BPA-C12-C12-cSS-cSS-cSS | **22** | C12-NDI-BPA-cSS-BPA-NDI |
| **3** | BPA-C12-cSS-cSS-cSS-C12-NDI | **23** | C12-BPA-NDI-cSS-NDI-BPA |
| **4** | C12-cSS-cSS-cSS-BPA-NDI-C12 | **24** | BPA-C12-BPA-cSS-NDI-NDI |
| **5** | C12-C12-cSS-cSS-cSS-HEG-cYY | **25** | BPA-BPA-cSS-NDI-C12-NDI |
| **6** | C12-cSS-C12-cSS-cSS-HEG-cYY | **26** | BPA-BPA-cSS-NDI-NDI-C12 |
| **7** | C12-HEG-cSS-C12-cSS-cSS-cYY | **27** | NDI-cSS-BPA-BPA-C12-NDI |
| **8** | C12-HEG-cSS-cSS-C12-cSS-cYY | **28** | NDI-BPA-cSS-C12-BPA-NDI |
| **9** | HEG-cSS-cSS-C12-cSS-C12-cYY | **29** | NDI-BPA-C12-C12-C12-NDI |
| **10** | HEG-cSS-cSS-cSS-cSS-cSS-NDI | **30** | NDI-BPA-NDI-C12-C12-C12 |
| **11** | HEG-cSS-cSS-cSS-cSS-NDI-cSS | **31** | BPA-NDI-C12-C12-C12-NDI |
| **12** | HEG-NDI-cSS-cSS-cSS-cSS-cSS | **32** | NDI-C12-C12-C12-HEG-cYY |
| **13** | NDI-HEG-cSS-cSS-cSS-cSS-cSS | **33** | C12-C12-NDI-C12-cYY-HEG |
| **14** | NDI-BPA-BPA-cSS-cSS-cSS-cSS | **34** | NDI-C12-C12-C12-cYY-HEG |
| **15** | BPA-NDI-BPA-cSS-cSS-cSS-cSS | **35** | C12-HEG-cSS-cSS-NDI-NDI |
| **16** | C12-BPA-BPA-cSS-C12-cSS-cYY | **36** | NDI-HEG-cSS-cSS-C12-NDI |
| **17** | BPA-BPA-cSS-cSS-cYY-C12-C12 | **37** | HEG-cSS-cSS-C12-NDI-NDI |
| **18** | cYY-cSS-cSS-C12-C12-BPA-BPA | **38** | DAN-cYY-cYY-DAN-DAN |
| **19** | cSS-cYY-BPA-C12-cSS-C12-BPA | **39** | cYY-cYY-DAN-DAN-DAN |
| **20** | cSS-cYY-BPA-C12-cSS-BPA-C12 | **40** | DAN-DAN-cYY-DAN-cYY |

Sequences 29 and 31 were identified as the most likely to fit within the data (Figure S 55) as they were the only ones above 95% (Figure S 55). The sequence would have had to contain one **BPA** monomer, two **NDI** monomers and three **C12** monomers.

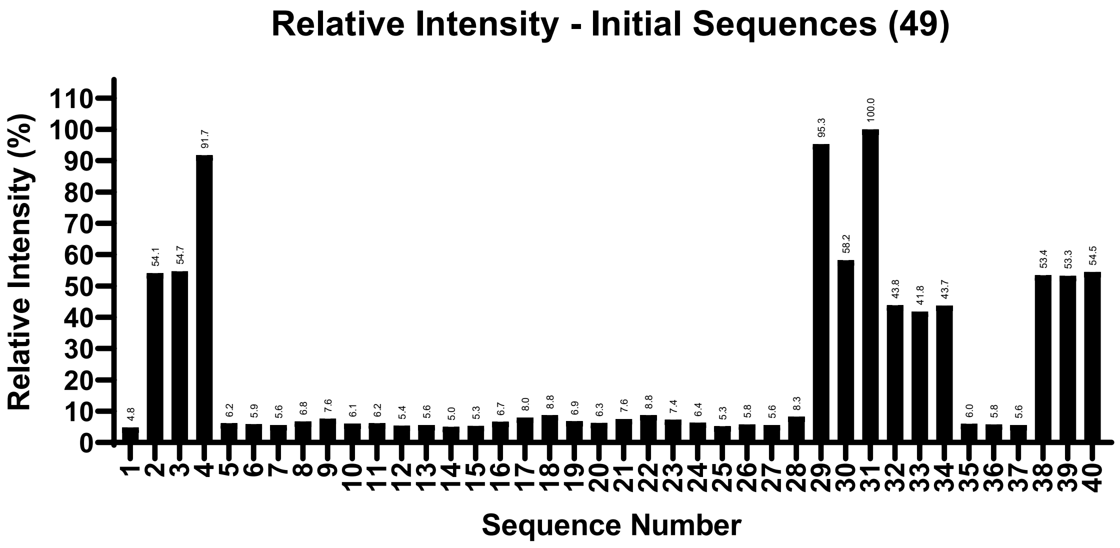

**Figure S 55.** Relative intensity comparisons of initial sequences for **O2**.

**Table S 12.** Possible sequences to fit monomer composition for **O2**. Continued on next page.

| **Sequence Number** | **Sequence (OH → P)** | **Sequence Number** | **Sequence (OH → P)** |
|---|---|---|---|
| **29** | NDI-BPA-C12-C12-C12-NDI | **68** | C12-BPA-NDI-NDI-C12-C12 |
| **30** | NDI-BPA-NDI-C12-C12-C12 | **69** | C12-BPA-NDI-C12-NDI-C12 |
| **31** | BPA-NDI-C12-C12-C12-NDI | **70** | C12-BPA-NDI-C12-C12-NDI |
| **41** | BPA-NDI-NDI-C12-C12-C12 | **71** | C12-BPA-C12-NDI-NDI-C12 |
| **42** | BPA-NDI-C12-NDI-C12-C12 | **72** | C12-BPA-C12-NDI-C12-NDI |
| **43** | BPA-NDI-C12-C12-NDI-C12 | **73** | C12-BPA-C12-C12-NDI-NDI |
| **44** | BPA-C12-NDI-NDI-C12-C12 | **74** | C12-NDI-BPA-NDI-C12-C12 |
| **45** | BPA-C12-NDI-C12-NDI-C12 | **75** | C12-NDI-BPA-C12-NDI-C12 |
| **46** | BPA-C12-NDI-C12-C12-NDI | **76** | C12-NDI-BPA-C12-C12-NDI |
| **47** | BPA-C12-C12-NDI-NDI-C12 | **77** | C12-NDI-NDI-BPA-C12-C12 |
| **48** | BPA-C12-C12-NDI-C12-NDI | **78** | C12-NDI-NDI-C12-BPA-C12 |
| **49** | BPA-C12-C12-C12-NDI-NDI | **79** | C12-NDI-NDI-C12-C12-BPA |
| **50** | NDI-BPA-C12-NDI-C12-C12 | **80** | C12-NDI-C12-BPA-NDI-C12 |
| **51** | NDI-BPA-C12-C12-NDI-C12 | **81** | C12-NDI-C12-BPA-C12-NDI |
| **52** | NDI-NDI-BPA-C12-C12-C12 | **82** | C12-NDI-C12-NDI-BPA-C12 |
| **53** | NDI-NDI-C12-BPA-C12-C12 | **83** | C12-NDI-C12-NDI-C12-BPA |

| | | | |
|---|---|---|---|
| **54** | NDI-NDI-C12-C12-BPA-C12 | **84** | C12-NDI-C12-C12-BPA-NDI |
| **55** | NDI-NDI-C12-C12-C12-BPA | **85** | C12-NDI-C12-C12-NDI-BPA |
| **56** | NDI-C12-BPA-NDI-C12-C12 | **86** | C12-C12-BPA-NDI-NDI-C12 |
| **57** | NDI-C12-BPA-C12-NDI-C12 | **87** | C12-C12-BPA-NDI-C12-NDI |
| **58** | NDI-C12-BPA-C12-C12-NDI | **88** | C12-C12-BPA-C12-NDI-NDI |
| **59** | NDI-C12-NDI-BPA-C12-C12 | **89** | C12-C12-NDI-BPA-NDI-C12 |
| **60** | NDI-C12-NDI-C12-BPA-C12 | **90** | C12-C12-NDI-BPA-C12-NDI |
| **61** | NDI-C12-NDI-C12-C12-BPA | **91** | C12-C12-NDI-NDI-BPA-C12 |
| **62** | NDI-C12-C12-BPA-NDI-C12 | **92** | C12-C12-NDI-NDI-C12-BPA |
| **63** | NDI-C12-C12-BPA-C12-NDI | **93** | C12-C12-NDI-C12-BPA-NDI |
| **64** | NDI-C12-C12-NDI-BPA-C12 | **94** | C12-C12-NDI-C12-NDI-BPA |
| **65** | NDI-C12-C12-NDI-C12-BPA | **95** | C12-C12-C12-BPA-NDI-NDI |
| **66** | NDI-C12-C12-C12-BPA-NDI | **96** | C12-C12-C12-NDI-BPA-NDI |
| **67** | NDI-C12-C12-C12-NDI-BPA | **97** | C12-C12-C12-NDI-NDI-BPA |

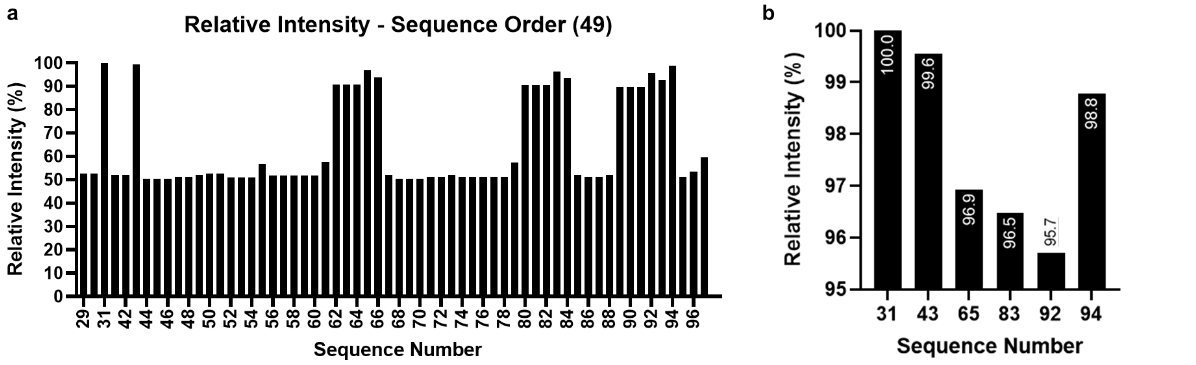

**Figure S 56.** (a) Relative intensities of possible sequences for **O2**. (b) Relative intensities of sequences above 95%.

The only distinct feature is comparing the top two sequences (31 and 43, Figure S 56) as these are almost identical, with the last two monomers in the sequence being swapped around being the only difference. This slight change did affect the RoboOligo analysis, as 31 had both c- and y- ions present which was not seen in 43 (Figure S 57), therefore reducing the cumulative intensity across the spectra and resulting in a lower relative intensity.

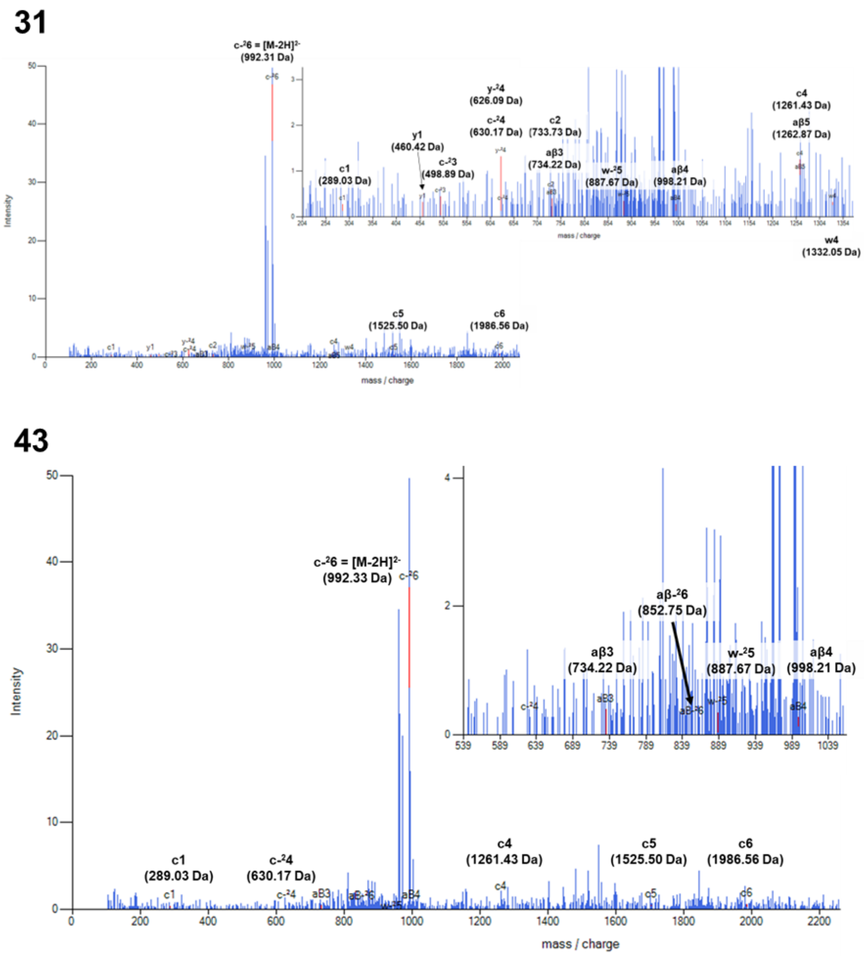

**Figure S 57.** RoboOligo data comparing sequence 31 (top) and sequence 43 (bottom) for **O2**.

Ultimately oligomer **49** was determined to be sequence 31 as this had the relative intensity of 100%, the structure of which is shown in Figure S 58.

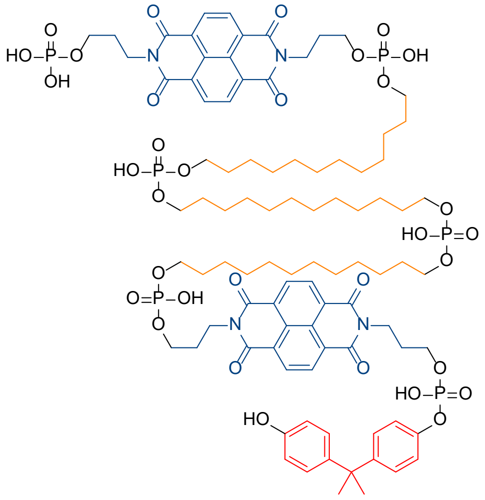

**O2**: HO-BPA-NDI-C12-C12-C12-NDI-P

**Figure S 58.** Best fit chemical structure of **O2**.

#### Oligomer 3

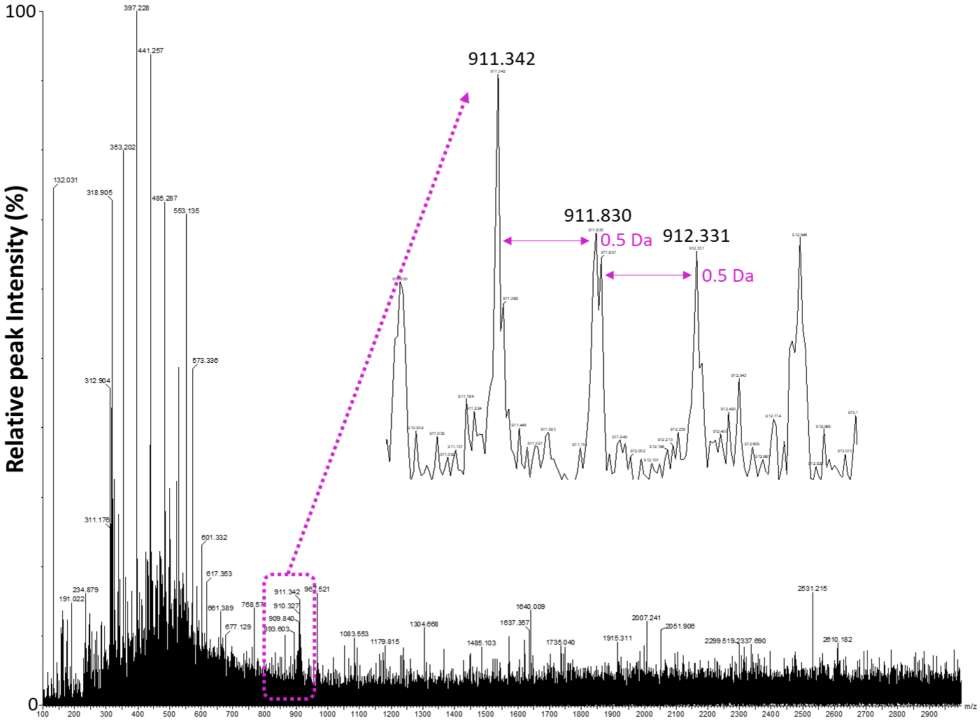

**Figure S 59.** *MS1 data from **O3** with the [M-2H]$^{2-}$ charge state highlighted.*

**Table S 13.** Sequence possibilities with the mass range of **O3**.

| **Mass** | **Monomer Combinations (OH → P)** | | | | | |
|---|---|---|---|---|---|---|
| **1824.684** | 1 | BPA | DAN | HEG | cYY | cYY |
| | 2 | DAN | DD | cSS | cYY | cYY |
| | 3 | DAN | HEG | HEG | cYY | cYY |

**Table S 14.** Initial sequences selected for RoboOligo analysis for **O3**.

| Sequence Number | Sequence (OH → P) |
|---|---|
| **1** | BPA-DAN-HEG-cYY-cYY |
| **2** | DAN-cYY-BPA-cYY-HEG |
| **3** | HEG-BPA-cYY-cYY-DAN |
| **4** | cYY-DAN-cYY-HEG-BPA |
| **5** | cYY-cYY-HEG-DAN-BPA |
| **6** | DAN-DAN-cSS-cYY-cYY |
| **7** | cSS-cYY-DAN-cYY-DAN |
| **8** | cYY-DAN-DAN-cSS-cYY |
| **9** | DAN-cYY-cYY-DAN-cSS |
| **10** | cYY-cYY-cSS-DAN-DAN |
| **11** | HEG-HEG-HEG-cYY-cYY |
| **12** | HEG-cYY-HEG-cYY-HEG |
| **13** | cYY-HEG-HEG-HEG-cYY |
| **14** | cYY-HEG-cYY-HEG-HEG |
| **15** | cYY-cYY-HEG-HEG-HEG |

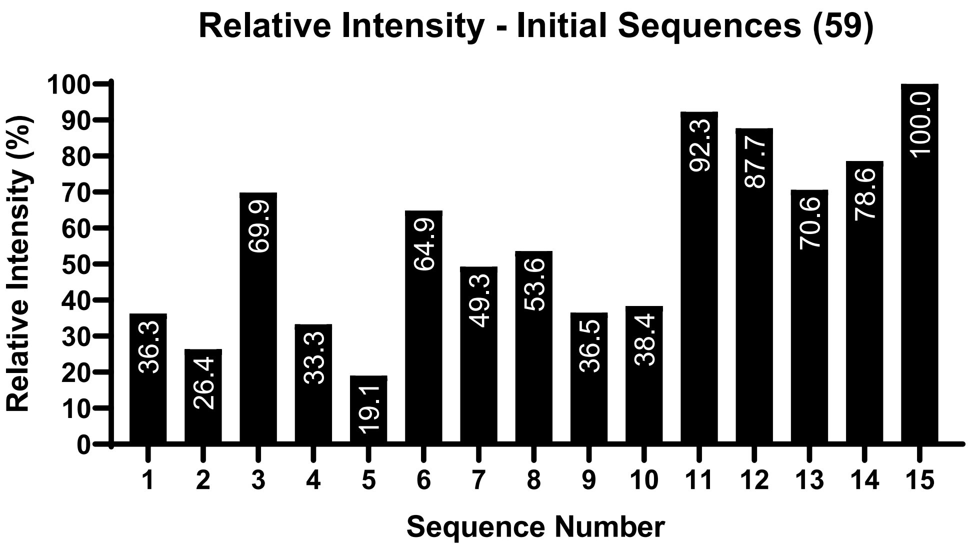

**Figure S 60.** Relative intensity comparisons of initial sequences for **O3**.

**Table S 15.** Possible sequences to fit monomer composition for **O3.**

| Sequence Number | Sequence (OH → P) |
|---|---|
| **11** | HEG-HEG-HEG-cYY-cYY |
| **12** | HEG-cYY-HEG-cYY-HEG |
| **13** | cYY-HEG-HEG-HEG-cYY |
| **14** | cYY-HEG-cYY-HEG-HEG |
| **15** | cYY-cYY-HEG-HEG-HEG |
| **16** | HEG-HEG-cYY-HEG-cYY |
| **17** | HEG-HEG-cYY-cYY-HEG |
| **18** | HEG-cYY-HEG-HEG-cYY |
| **19** | HEG-cYY-cYY-HEG-HEG |
| **20** | cYY-HEG-HEG-cYY-HEG |

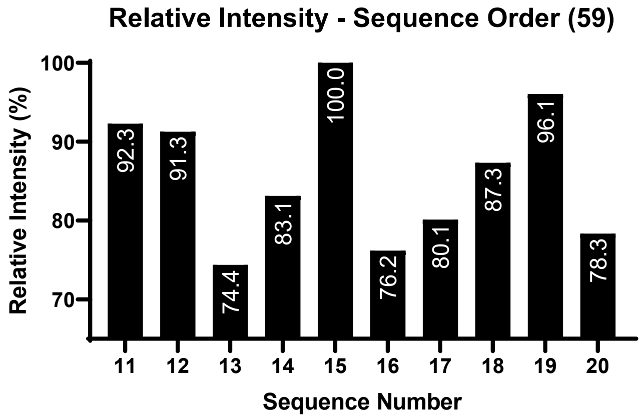

**Figure S 61.** Relative intensities of different sequences for **O3**.

The RoboOligo analysis (Figure S 62) identified a number of peaks consistent with the top ranked sequence, 15.

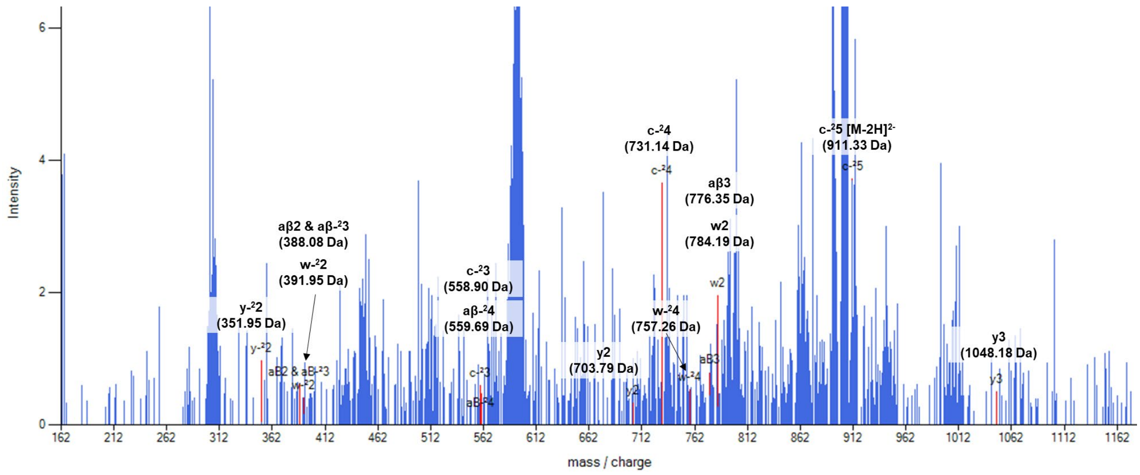

**Figure S 62.** RoboOligo data from the top sequence of **O3**.

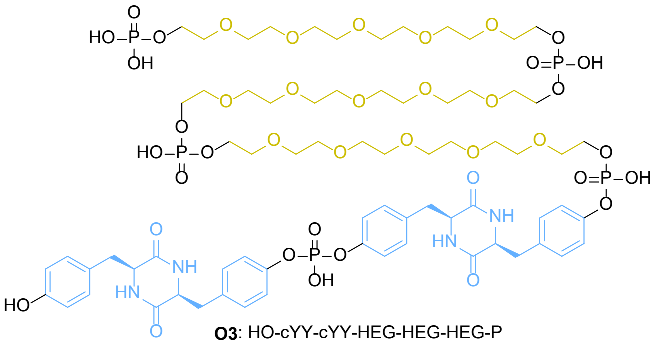

**Figure S 63.** Best fit chemical structure for **O3**.

#### Oligomer 4

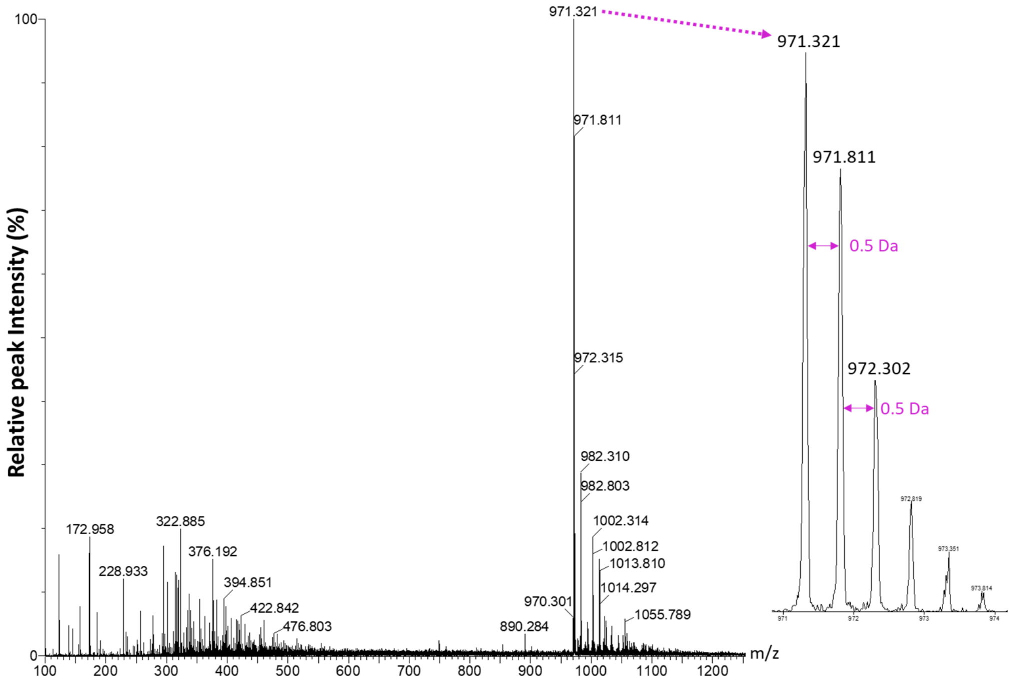

**Figure S 64.** MS1 data of **O4** from the chromatogram at 31 minutes, with $[M-2H]^{2-}$ charge state highlighted.

**Table S 16.** Sequence possibilities with the mass range of **O4**.

| **Mass** | | **Monomer Combinations (OH → P)** | | | | | | |
|---|---|---|---|---|---|---|---|---|
| **1944.616** | **1** | BPA | BPA | BPA | C12 | C12 | C12 | C12 |
| | **2** | BPA | C12 | C12 | C12 | C12 | HEG | cSS |
| | **3** | C12 | C12 | C12 | C12 | C12 | C12 | HEG |
| | **4** | C12 | C12 | C12 | C12 | DAN | cSS | cSS |
| | **5** | BPA | cSS | cSS | cYY | cYY | cYY | - |
| | **6** | C12 | C12 | cSS | cYY | cYY | cYY | - |
| | **7** | cSS | cSS | cSS | cYY | cYY | NDI | - |
| | **8** | BPA | DAN | DAN | DAN | NDI | - | - |
| | **9** | C12 | cYY | cYY | NDI | NDI | - | - |
| | **10** | DAN | DAN | DAN | HEG | cYY | - | - |
| | **11** | DAN | DAN | HEG | HEG | NDI | - | - |

**Table S 17.** Initial sequences selected for RoboOligo analysis for **O4**.

| Sequence Number | Sequence (OH → P) | Sequence Number | Sequence (OH → P) |
|---|---|---|---|
| 1 | BPA-BPA-BPA-C12-C12-C12-C12 | **23** | cYY-C12-cYY-C12-cYY-cSS |
| 2 | C12-BPA-C12-BPA-C12-BPA-C12 | **24** | cYY-cYY-cYY-cSS-C12-C12 |
| 3 | C12-C12-BPA-BPA-C12-C12-BPA | **25** | cSS-cSS-cSS-cYY-cYY-NDI |
| 4 | C12-C12-C12-C12-BPA-BPA-BPA | **26** | cYY-cSS-cYY-cSS-NDI-cSS |
| 5 | BPA-C12-C12-C12-C12-HEG-cSS | **27** | cYY-cSS-NDI-cSS-cSS-cYY |
| 6 | HEG-C12-C12-BPA-cSS-C12-C12 | **28** | NDI-cYY-cYY-cSS-cSS-cSS |
| 7 | C12-BPA-C12-cSS-C12-C12-HEG | **29** | BPA-DAN-DAN-DAN-NDI |
| 8 | cSS-HEG-C12-C12-C12-C12-BPA | **30** | DAN-BPA-DAN-NDI-DAN |
| 9 | C12-C12-C12-C12-C12-C12-HEG | **31** | DAN-NDI-BPA-DAN-DAN |
| 10 | C12-C12-C12-HEG-C12-C12-C12 | **32** | NDI-DAN-DAN-DAN-BPA |
| 11 | C12-C12-HEG-C12-C12-C12-C12 | **33** | C12-cYY-cYY-NDI-NDI |
| 12 | HEG-C12-C12-C12-C12-C12-C12 | **34** | cYY-NDI-C12-NDI-cYY |
| 13 | C12-C12-C12-C12-DAN-cSS-cSS | **35** | NDI-C12-cYY-cYY-NDI |
| 14 | DAN-C12-cSS-C12-cSS-C12-C12 | **36** | NDI-NDI-cYY-cYY-C12 |
| 15 | cSS-C12-C12-cSS-C12-C12-DAN | **37** | DAN-DAN-DAN-HEG-cYY |
| 16 | cSS-cSS-DAN-C12-C12-C12-C12 | **38** | DAN-DAN-cYY-DAN-HEG |
| 17 | BPA-cSS-cSS-cYY-cYY-cYY | **39** | HEG-DAN-DAN-DAN-cYY |
| 18 | cYY-cSS-cYY-BPA-cYY-cSS | **40** | cYY-HEG-DAN-DAN-DAN |
| 19 | cSS-cYY-BPA-cYY-cSS-cYY | **41** | DAN-DAN-HEG-HEG-NDI |
| 20 | cYY-cYY-cYY-cSS-cSS-BPA | **42** | HEG-DAN-NDI-DAN-HEG |
| 21 | C12-C12-cSS-cYY-cYY-cYY | **43** | NDI-HEG-DAN-HEG-DAN |
| 22 | C12-cYY-cSS-cYY-cYY-C12 | **44** | NDI-HEG-HEG-DAN-DAN |

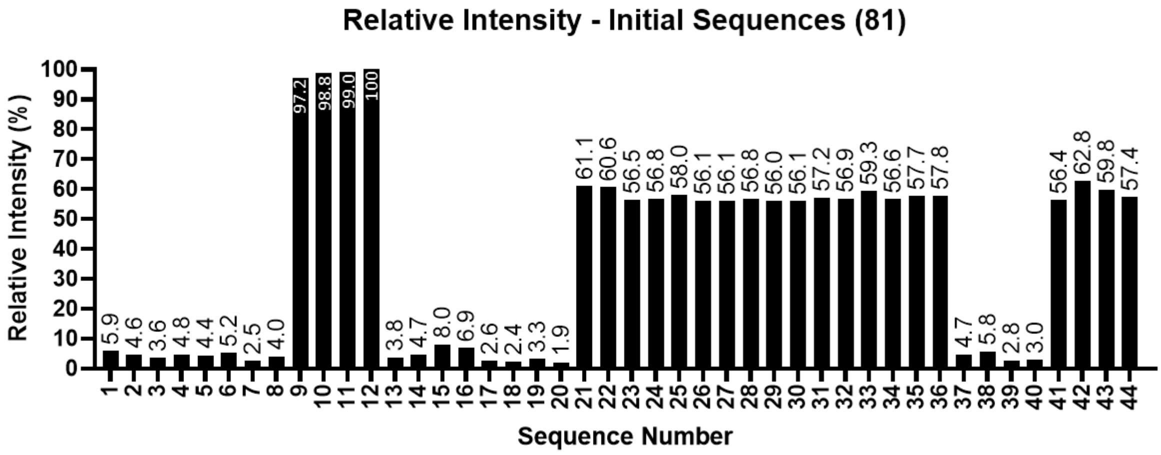

**Figure S 65.** Relative intensity comparisons of initial sequences for **O4**

**Table S 18.** Possible sequences to fit monomer composition for **O4**.

| Sequence Number | Sequence (OH – P) |
|---|---|
| **9** | C12-C12-C12-C12-C12-C12-HEG |
| **10** | C12-C12-C12-HEG-C12-C12-C12 |
| **11** | C12-C12-HEG-C12-C12-C12-C12 |
| **12** | HEG-C12-C12-C12-C12-C12-C12 |
| **45** | C12-HEG-C12-C12-C12-C12-C12 |
| **46** | C12-C12-C12-C12-HEG-C12-C12 |
| **47** | C12-C12-C12-C12-C12-HEG-C12 |

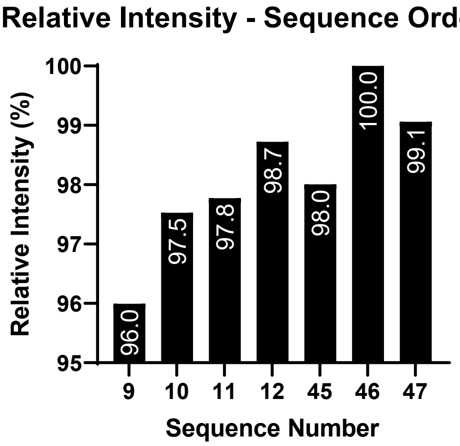

**Figure S 66.** Relative intensities of different sequences to match best fit composition of **O4**.

Examining the RoboOligo data, the change in HEG placement would affect the overall y- ion cumulative intensity and sequence 46 contains more y- ions compared to the other sequences (Figure S 67).

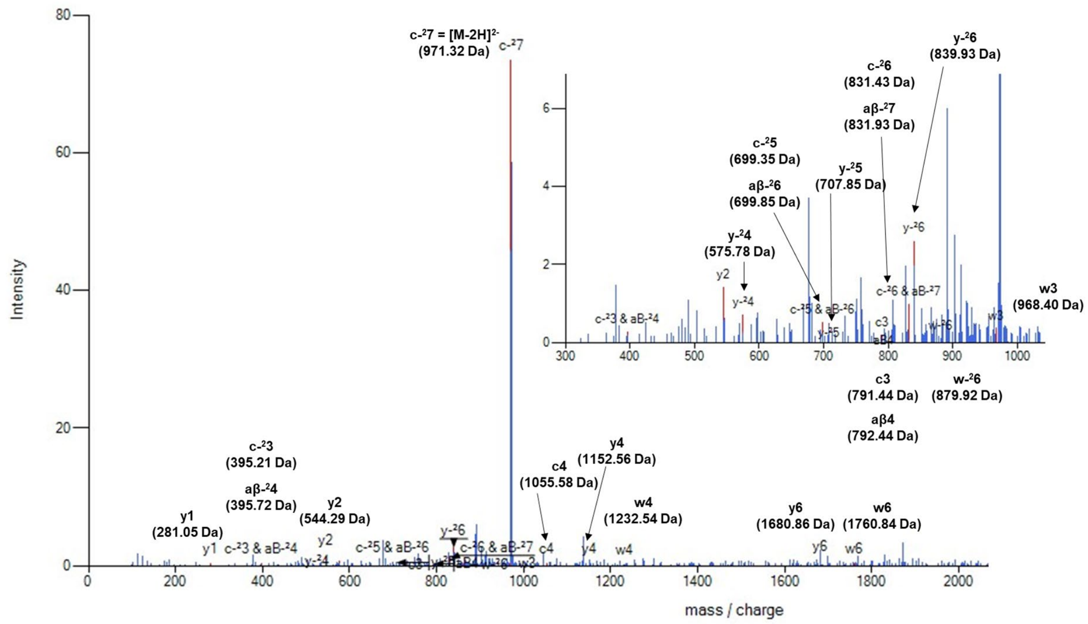

**Figure S 67.** RoboOligo data from candidate sequence 46 of oligomer **O4**.

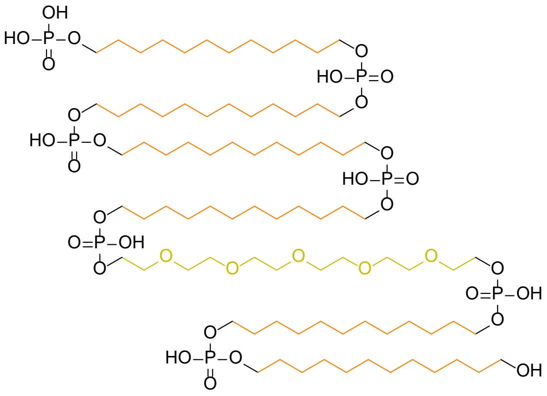

**O4**: HO-C12-C12-C12-C12-HEG-C12-C12-P

**Figure S 68.** Best fit chemical structure for **O4**.

**Oligomer 5**

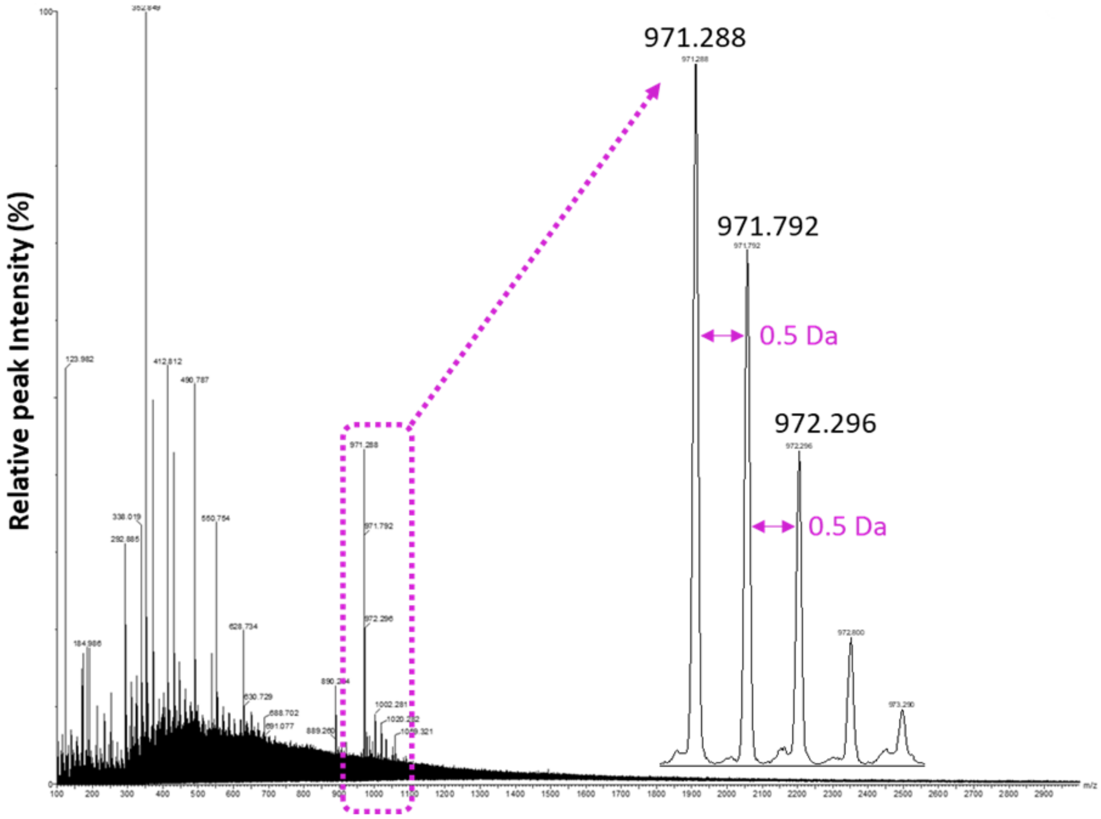

**Figure S 69.** MS1 data from **O5**, with the $[M-2H]^{2-}$ charge state highlighted.

**Table S 19.** Initial sequences selected for RoboOligo analysis for **O5**. Table continued on following page.

| Sequence Number | Sequence (OH → P) | Sequence Number | Sequence (OH → P) |
|---|---|---|---|
| **1** | C12-C12-HEG-C12-C12-C12-C12 | **29** | C12-cYY-cYY-C12-cYY-cSS |
| **2** | C12-C12-C12-C12-C12-C12-HEG | **30** | cYY-cYY-cYY-cSS-C12-C12 |
| **3** | C12-HEG-C12-C12-C12-C12-C12 | **31** | cSS-cSS-cSS-cYY-cYY-NDI |
| **4** | C12-C12-C12-C12-HEG-C12-C12 | **32** | NDI-cYY-cSS-cYY-cSS-cSS |
| **5** | HEG-C12-C12-C12-C12-C12-C12 | **33** | cSS-NDI-cSS-cYY-cSS-cYY |
| **6** | BPA-BPA-BPA-C12-C12-C12-C12 | **34** | cYY-cSS-cSS-cSS-cYY-NDI |
| **7** | C12-BPA-C12-BPA-C12-BPA-C12 | **35** | NDI-cYY-cYY-cSS-cSS-cSS |
| **8** | C12-C12-BPA-BPA-BPA-C12-C12 | **36** | BPA-DAN-DAN-DAN-NDI |
| **9** | BPA-C12-C12-BPA-C12-C12-BPA | **37** | DAN-BPA-DAN-NDI-DAN |
| **10** | C12-C12-C12-C12-BPA-BPA-BPA | **38** | NDI-DAN-DAN-BPA-DAN |
| **11** | BPA-C12-C12-C12-C12-HEG-cSS | **39** | BPA-DAN-NDI-DAN-DAN |

| | | | |
|---|---|---|---|
| **12** | C12-cSS-BPA-HEG-C12-C12-C12 | **40** | NDI-DAN-DAN-DAN-BPA |
| **13** | HEG-C12-C12-cSS-C12-C12-BPA | **41** | C12-cYY-cYY-NDI-NDI |
| **14** | BPA-HEG-C12-C12-cSS-C12-C12 | **42** | NDI-cYY-C12-cYY-NDI |
| **15** | cSS-HEG-C12-C12-C12-C12-BPA | **43** | cYY-cYY-C12-NDI-NDI |
| **16** | C12-C12-C12-C12-DAN-cSS-cSS | **44** | cYY-C12-NDI-NDI-cYY |
| **17** | C12-C12-cSS-C12-C12-cSS-DAN | **45** | NDI-NDI-cYY-cYY-C12 |
| **18** | DAN-cSS-C12-cSS-C12-C12-C12 | **46** | DAN-DAN-DAN-HEG-cYY |
| **19** | cSS-C12-C12-C12-C12-cSS-DAN | **47** | DAN-HEG-DAN-cYY-DAN |
| **20** | cSS-cSS-DAN-C12-C12-C12-C12 | **48** | HEG-DAN-cYY-DAN-DAN |
| **21** | BPA-cSS-cSS-cYY-cYY-cYY | **49** | cYY-DAN-DAN-DAN-HEG |
| **22** | BPA-cYY-cSS-cYY-cSS-cYY | **50** | cYY-HEG-DAN-DAN-DAN |
| **23** | cSS-cYY-cYY-BPA-cYY-cSS | **51** | DAN-DD-HEG-HEG-NDI |
| **24** | cYY-BPA-cYY-cSS-cYY-cSS | **52** | DAN-HEG-NDI-HEG-DAN |
| **25** | cYY-cYY-cYY-cSS-cSS-BPA | **53** | HEG-DAN-DAN-NDI-HEG |
| **26** | C12-C12-cSS-cYY-cYY-cYY | **54** | NDI-DAN-HEG-DAN-HEG |
| **27** | cSS-cYY-C12-cYY-C12-cYY | **55** | NDI-HEG-HEG-DAN-DAN |
| **28** | cYY-C12-C12-cYY-cYY-cSS | | |

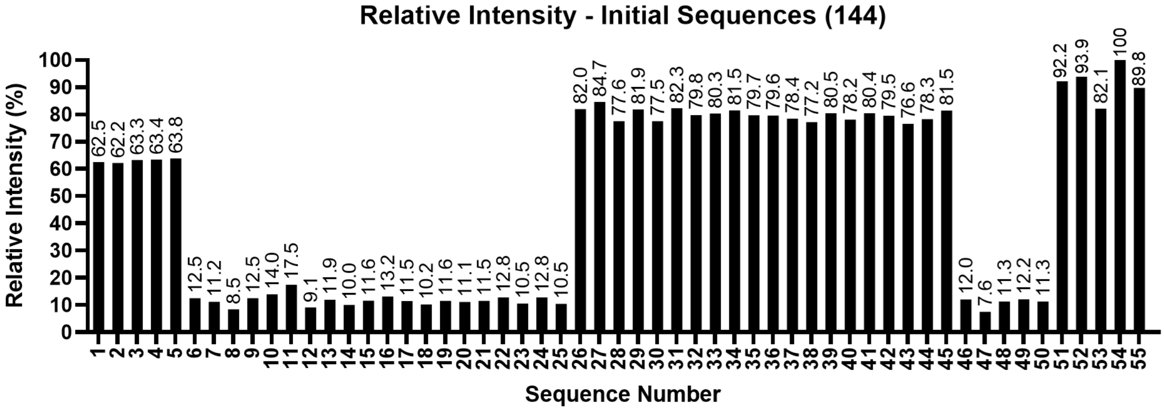

**Figure S 70.** Relative intensity comparisons of initial sequences for **O5**.

**Table S 20.** Possible sequences to fit monomer composition for **O5**.

| Sequence Number | Sequence (OH → P) | Sequence Number | Sequence (OH → P) |
|---|---|---|---|
| **51** | DAN-DAN-HEG-HEG-NDI | **66** | HEG-DAN-NDI-HEG-DAN |
| **52** | DAN-HEG-NDI-HEG-DAN | **67** | HEG-DAN-NDI-DAN-HEG |
| **53** | HEG-DAN-DAN-NDI-HEG | **68** | HEG-DAN-HEG-NDI-DAN |
| **54** | NDI-DAN-HEG-DAN-HEG | **69** | HEG-DAN-HEG-DAN-NDI |
| **55** | NDI-HEG-HEG-DAN-DAN | **70** | HEG-DAN-DAN-HEG-NDI |
| **56** | NDI-HEG-DAN-HEG-DAN | **71** | DAN-NDI-HEG-HEG-DAN |
| **57** | NDI-HEG-DAN-DAN-HEG | **72** | DAN-NDI-HEG-DAN-HEG |
| **58** | NDI-DAN-HEG-HEG-DAN | **73** | DAN-NDI-DAN-HEG-HEG |
| **59** | NDI-DAN-DAN-HEG-HEG | **74** | DAN-HEG-NDI-DAN-HEG |
| **60** | HEG-NDI-HEG-DAN-DAN | **75** | DAN-HEG-HEG-NDI-DAN |
| **61** | HEG-NDI-DAN-HEG-DAN | **76** | DAN-HEG-HEG-DAN-NDI |
| **62** | HEG-NDI-DAN-DAN-HEG | **77** | DAN-HEG-DAN-NDI-HEG |
| **63** | HEG-HEG-NDI-DAN-DAN | **78** | DAN-HEG-DAN-HEG-NDI |
| **64** | HEG-HEG-DAN-NDI-DAN | **79** | DAN-DAN-NDI-HEG-HEG |
| **65** | HEG-HEG-DAN-DAN-NDI | **80** | DAN-DAN-HEG-NDI-HEG |
| **66** | HEG-DAN-NDI-HEG-DAN | | |

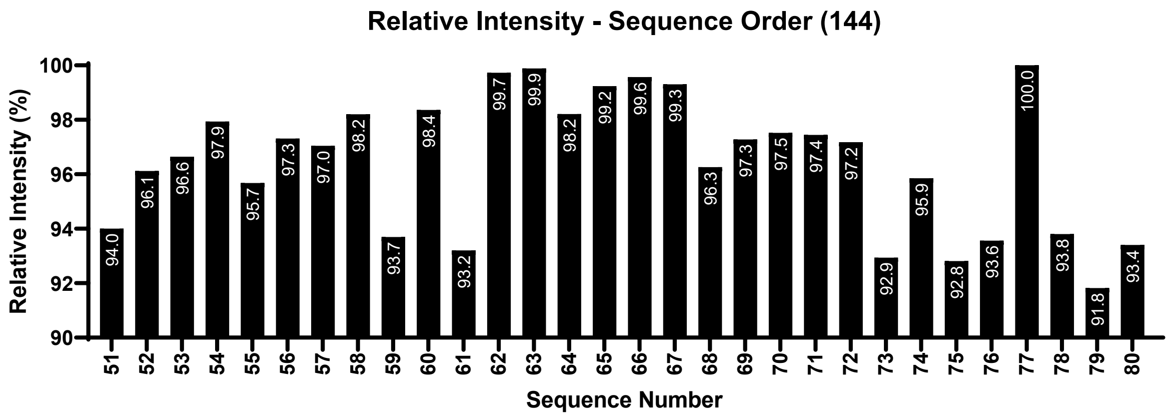

**Figure S 71.** Relative intensities of different sequences to match best fit composition of **O5**.

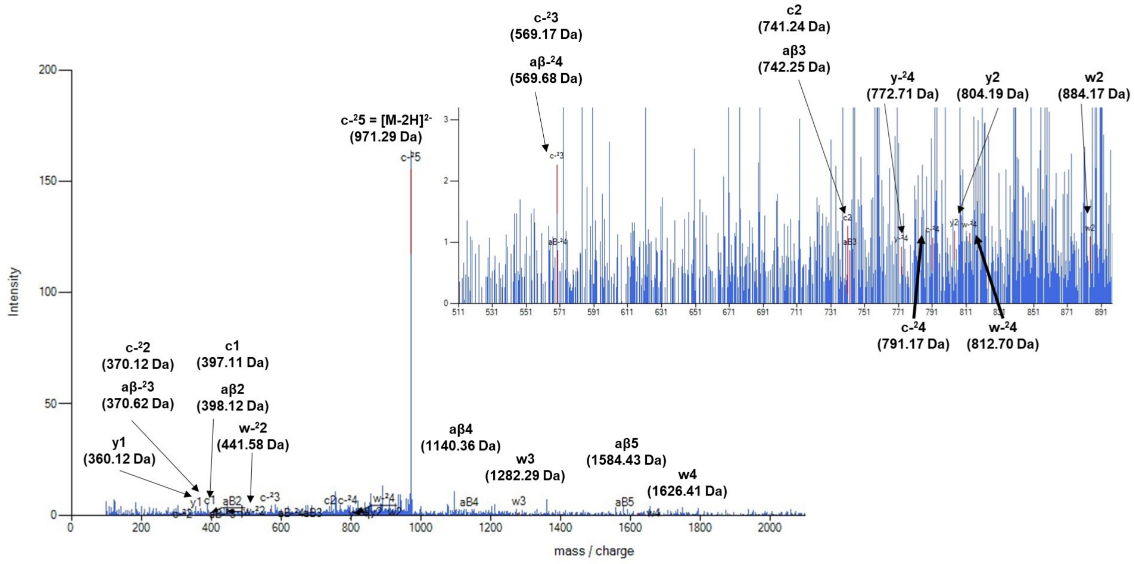

**Figure S 72.** RoboOligo data from sequence 77 of **O5**.

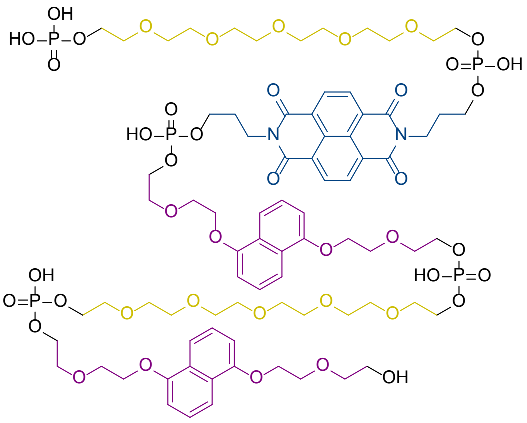

**O5**: HO-DAN-HEG-DAN-NDI-HEG-P

**Figure S 73.** Best fit chemical structure for **O5**.

**Oligomer 6**

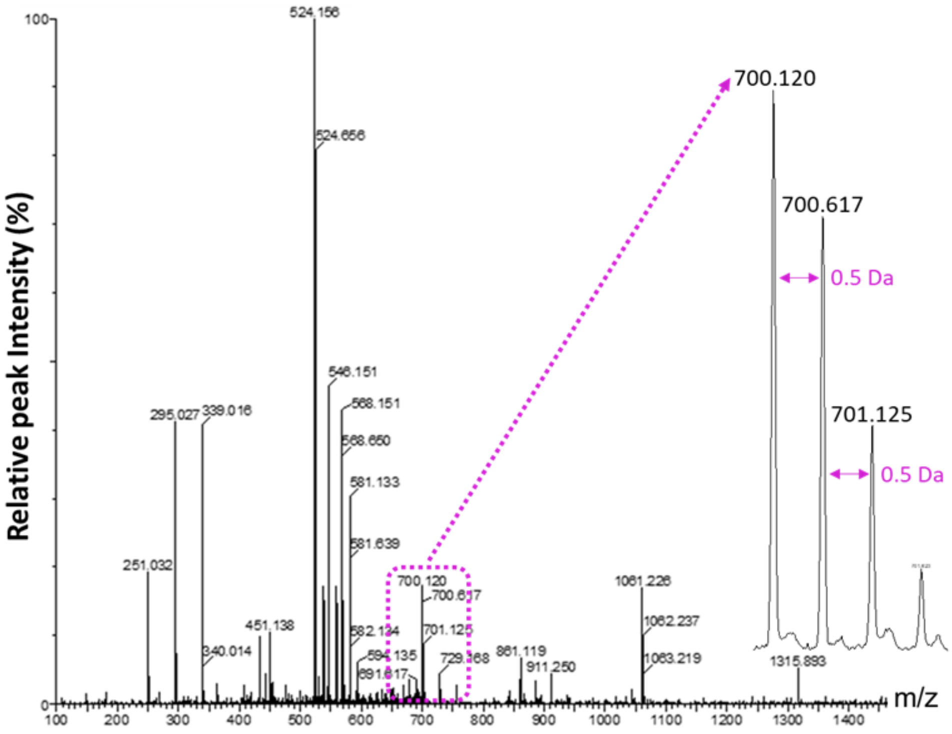

**Figure S 74.** MS1 data from oligomer **O6**, with the $[M-2H]^{2-}$ charge state highlighted.

**Table S 21.** Monomer composition possibilities within the mass range of **O6**.

| **Mass** | | **Monomer Combinations** | | | | |
|---|---|---|---|---|---|---|
| **1402.24** | **1** | BPA | cSS | cSS | cSS | cYY |
| | **2** | C12 | HEG | cYY | cYY | - |
| | **3** | BPA | C12 | cYY | NDI | - |

**Table S 22.** Initial sequences selected for RoboOligo analysis for **O6**.

| Sequence Number | Sequence (OH → P) |
|---|---|
| 1 | BPA-cSS-cSS-cSS-cYY |
| 2 | cSS-cYY-cSS-BPA-cSS |
| 3 | cYY-cSS-BPA-cSS-cSS |
| 4 | cSS-CPA-cYY-cSS-cSS |
| 5 | cYY-cSS-cSS-cSS-BPA |
| 6 | C12-HEG-cYY-cYY |
| 7 | cYY-HEG-cYY-C12 |
| 8 | HEG-cYY-cYY-C12 |
| 9 | C12-cYY-cYY-HEG |
| 10 | cYY-cYY-HEG-C12 |
| 11 | BPA-C12-cYY-NDI |
| 12 | cYY-BPA-NDI-C12 |
| 13 | cYY-C12-NDI-BPA |
| 14 | NDI-BPA-cYY-C12 |
| 15 | NDI-cYY-C12-BPA |

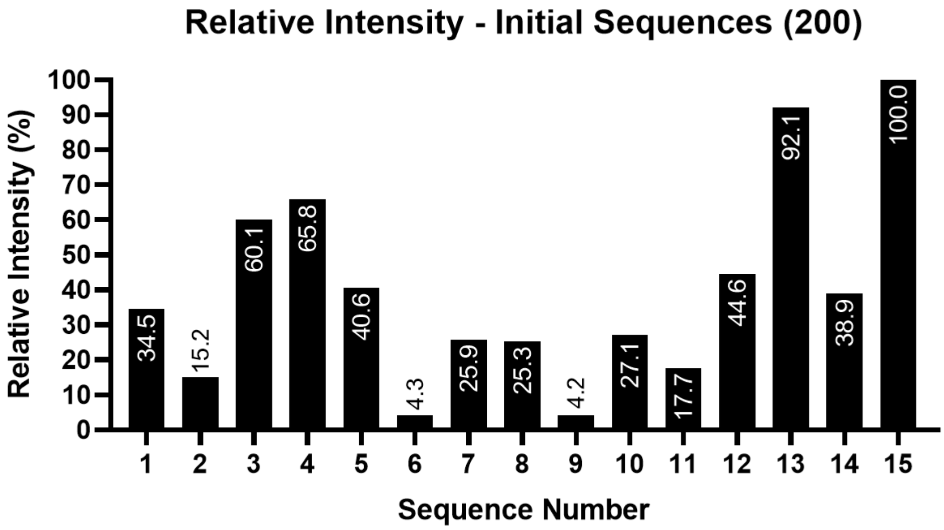

**Figure S 75.** Relative intensity comparisons of initial sequences for **O6**.

**Table S 23.** Possible sequences to fit monomer composition for **O6**.

| Sequence Number | Sequence (OH – P) | Sequence Number | Sequence (OH – P) |
|---|---|---|---|
| **11** | BPA-C12-cYY-NDI | **23** | C12-cYY-BPA-NDI |
| **12** | cYY-BPA-NDI-C12 | **24** | C12-cYY-NDI-BPA |
| **13** | cYY-C12-NDI-BPA | **25** | C12-NDI-cYY-BPA |
| **14** | NDI-BPA-cYY-C12 | **26** | cYY-BPA-C12-NDI |
| **15** | NDI-cYY-C12-BPA | **27** | C12-NDI-BPA-cYY |
| **16** | BPA-C12-NDI-cYY | **28** | cYY-C12-BPA-NDI |
| **17** | BPA-cYY-C12-NDI | **29** | cYY-NDI-BPA-C12 |
| **18** | BPA-cYY-NDI-C12 | **30** | cYY-NDI-C12-BPA |
| **19** | BPA-NDI-C12-cYY | **31** | NDI-BPA-C12-cYY |
| **20** | BPA-NDI-cYY-C12 | **32** | NDI-C12-BPA-cYY |
| **21** | C12-BPA-cYY-NDI | **33** | NDI-C12-cYY-BPA |
| **22** | C12-BPA-NDI-cYY | **34** | NDI-cYY-BPA-C12 |

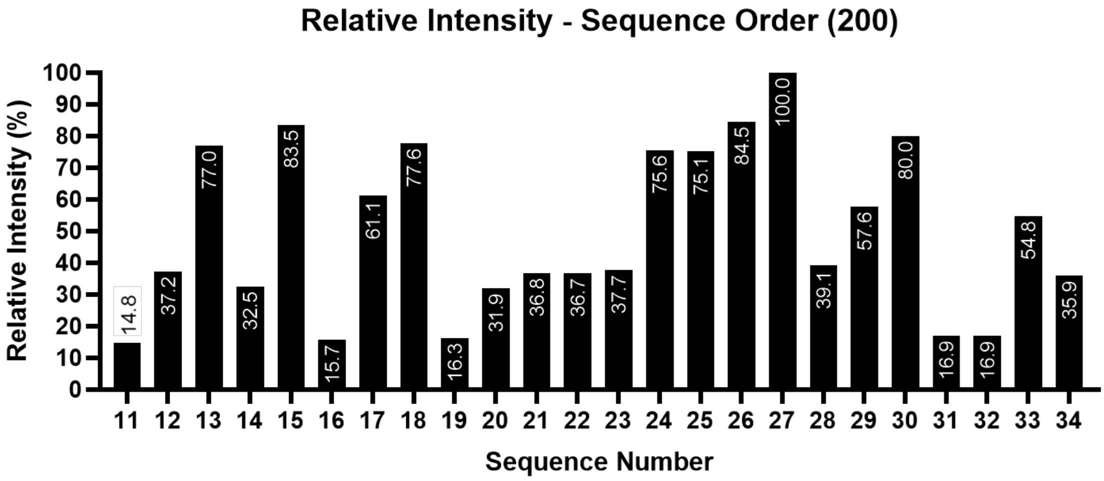

**Figure S 76.** Relative intensities of different sequences to match best fit composition of **O6**.

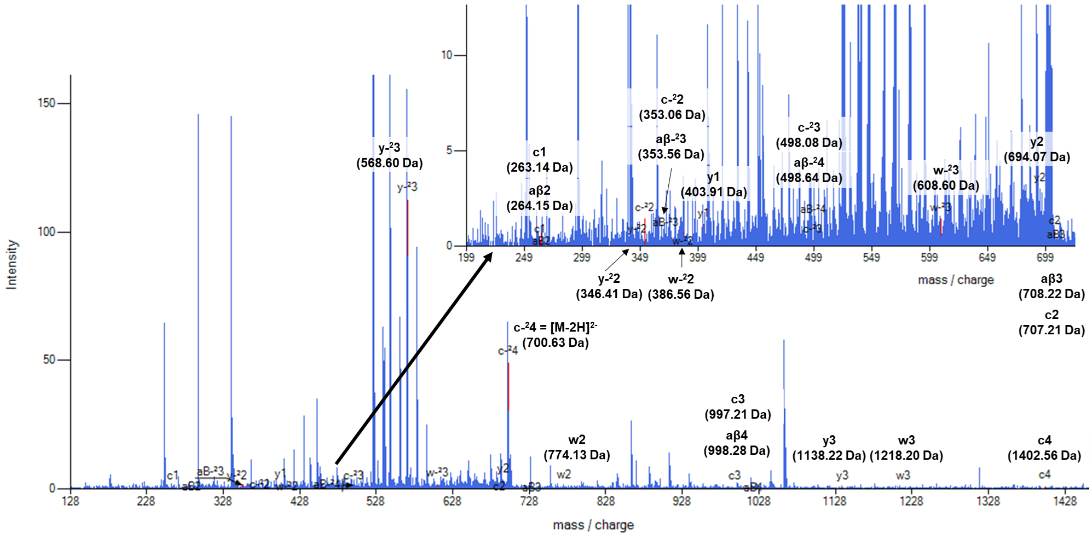

**Figure S 77.** RoboOligo data from sequence 27 of **O6**.

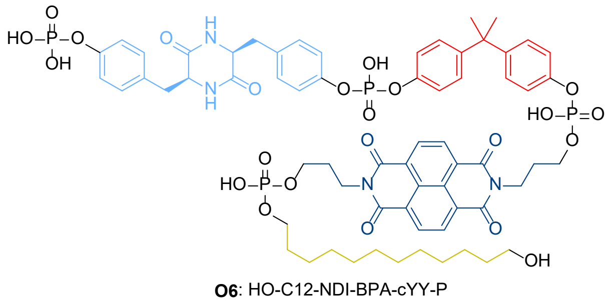

**Figure S 78.** Best fit chemical structure for **O6**.

### 6 Validation assays

**General procedure**

To the 96 well plate, 100 µL was added to each well containing 2 µg/mL $KRAS^{G12D}$-GMPPnP or $KRAS^{WT}$-GMPPnP in coating buffer (see buffers, page 3). The plate was shaken on a plate rotator at 300 rpm and left to incubate at 4 °C overnight. After incubation, the $KRAS^{G12D/WT}$ solution was removed, and the plate was then washed with 100 µL per well of blocking buffer twice. 100 µL of blocking buffer was then added per well and the plate was incubated at 37 °C for one hour. After incubation the blocking buffer was removed and the plate was then washed with 100 µL per well of PBS twice, then 2 µg/mL RAF1-GFP (100 µL) (made up in PBS Buffer) and required oligomers (100 µL – variable concentrations) (made up in PBS Buffer) were added to their respective wells in the plate. The plate was then shaken on a plate rotator at 300 rpm at room temperature for 15 minutes before incubation at 37 °C for one hour. The well-plate was then washed with 100 µL per well PBS twice, with any excess liquid removed from the well-plate. The emission spectrum of GFP (490 – 560 nm) was measured on the plate reader. GraphPad Prism (9.5.1) was used to calculate the area under the curve (AUC) of the emission spectra between 490 and 540 nm. $IC_{50}$ values were calculated on GraphPad Prism, using the non-linear regression fit (Sigmoidal, 4PL, X is concentration) analysis.

**KRAS binding assay**

To the well plate, 100 µL was added to each well containing 2 µg/mL of the 4:1 conjugate of $KRAS^{G12D}$-GMPPnP-streptavidin fluorescein in coating buffer. The plate was then shaken on a plate rotator at 300 rpm and left to incubate at 4 °C overnight. After incubation the $KRAS^{G12D}$ solution was removed, and the plate was then washed with 100 µL per well of blocking buffer twice. 100 µL of blocking buffer was then added per well and the plate was incubated at 37 °C for one hour. The well plate was then washed with 100 µL per well of PBS twice, shaken on a plate rotator at 300 rpm at room temperature for 15 minutes before incubation at 37 °C for one hour. The plate was then washed with 100 µL per well PBS twice, with any excess liquid removed from the plate. The emission spectrum of fluorescein (500 – 600 nm) was measured using the plate reader. GraphPad Prism (9.5.1) was used to calculate the area under the curve (AUC) of the fluorescein emission spectra.

**Positive control assays**

Assays were run following the General Assay Protocol. Positive control **2** was diluted to the following concentrations in PBS Buffer: 15, 10, 7.5, 5, 2.5, and 1.25 µM. Each assay also contained a well with 0 µM sample (only $KRAS^{G12D}$ and RAF1-GFP), a well with only $KRAS^{G12D}$-GMPPnP, and a well containing only **2** (at 15 µM). Each assay condition was run in triplicate wells and the assay was performed to achieve three biological repeats (n = 3).

**Initial oligomer assays to determine concentration ranges.**

Using the DMT cleavage protocol to determine the oligomer concentrations, in 1 mL of solution the concentrations were as shown in Table S 24.

**Table S 24.** Determined oligomer concentrations.

| Oligomer | O1 | O2 | O3 | O4 | O5 | O6 |
|---|---|---|---|---|---|---|
| **Concentration (µM)** | 1.971 | 1.234 | 1.003 | 4.362 | 1.544 | 2.960 |

The oligomers were left to dry and resuspended in 2 mL of KRAS buffer. These were used as stock solutions for the oligomers.

Starting with the highest concentration, a 0.5 x serial dilution across 12 concentration points (including a 0 µM concentration point) was performed, resulting in the following concentrations:

**O1**: 0.985, 0.493, 0.246, 0.123, 0.062, 0.031, 0.015, 0.008, 0.004, 0.002, 0.001 and 0 µM

**O2**: 0.617, 0.309, 0.154, 0.077, 0.039, 0.019, 0.010, 0.005, 0.002, 0.001, 0.0006 and 0 µM

**O3**: 1.003, 0.502, 0.251, 0.125, 0.063, 0.031, 0.016, 0.008, 0.004, 0.002, 0.001 and 0 µM

**O4**: 2.181, 1.091, 0.545, 0.273, 0.136, 0.068, 0.034, 0.017, 0.009, 0.004, 0.002 and 0 µM

**O5**: 0.772, 0.386, 0.193, 0.097, 0.048, 0.024, 0.012, 0.006, 0.003, 0.0015, 0.00075 and 0 µM

**O6**: 1.480, 0.740, 0.370, 0.185, 0.093, 0.046, 0.023, 0.012, 0.0058, 0.0029, 0.0014 and 0 µM

Using these concentrations, the assay was performed using the General Assay Procedure.

**$KRAS^{G12D}$-GMPPnP assays**

Assays were run following the General Assay Protocol, with oligomers diluted within various concentration ranges. Each assay also contained a well with only $KRAS^{G12D}$-GMPPnP, and a well with $KRAS^{G12D}$-GMPPnP, RAF1-GFP and **2** at 15 µM as negative controls to ensure that readings within the assay represented actual results. Each assay was run in triplicate wells and the assay was performed to achieve three biological repeats (n = 3). The assay for oligomer **O4** was performed in quadruplicate (n = 4) due to a large variation in data.

**$KRAS^{WT}$-GMPPnP assays**

Assays were run following the General Assay Protocol (5.2.3.9). Compound **Ch-3** was diluted to 60, 10 and 1 µM in PBS Buffer and run in triplicate wells per assay with 3 biological repeats of the assay (n = 3). The oligomers were diluted to three different concentrations for analysis. Each assay also contained a well with only $KRAS^{WT}$-GMPPnP, and a well with $KRAS^{WT}$-GMPPnP, RAF1-GFP and Ch-3 at 15 µM to ensure the assay was working correctly. Assay conditions for **O1**, **O2** and **O3** were run in triplicate wells and the assay was performed to achieve three biological repeats (n = 3). **O5** had 2 repeats (n = 2), but after seeing a change in signal between 15 and 150 nM a full data series was run as an alternative. **O4** and **O6** were run in triplicate once (n = 1). Statistical analysis (Dunnett's multiple comparisons test) was used to compare 0 nM to all other concentrations using GraphPad Prism (9.5.1).

**$KRAS^{WT}$-GMPPnP assay – O5 full data series.**

**O5** was diluted to the following concentrations in PBS Buffer: 250, 200, 150, 100, 50, 25 and 15 µM. The assays were run using the General Assay Protocol. Each assay also contained a well at 0 µM of sample, a well with only $KRAS^{WT}$-GMPPnP, and a well with $KRAS^{WT}$-GMPPnP, RAF1-GFP and **Ch-3** at 15 µM to ensure the assay was working correctly. Each assay condition was run in triplicate and the assay was performed to achieve three biological repeats (n = 3).

### 7 References

1 Pierce™ C18 Spin Tips & Columns, https://www.thermofisher.com/order/catalog/product/84850, (accessed 30 May 2023).

2 P. J. Sample, K. W. Gaston, J. D. Alfonzo and P. A. Limbach, RoboOligo: software for mass spectrometry data to support manual and de novo sequencing of post-transcriptionally modified ribonucleic acids, *Nucleic Acids Res.*, 2015, **43**, e64.

3 D. Kessner, M. Chambers, R. Burke, D. Agus and P. Mallick, ProteoWizard: open source software for rapid proteomics tools development, *Bioinformatics*, 2008, **24**, 2534–2536.

4 M. C. Chambers, B. Maclean, R. Burke, D. Amodei, D. L. Ruderman, S. Neumann, L. Gatto, B. Fischer, B. Pratt, J. Egertson, K. Hoff, D. Kessner, N. Tasman, N. Shulman, B. Frewen, T. A. Baker, M.-Y. Brusniak, C. Paulse, D. Creasy, L. Flashner, K. Kani, C. Moulding, S. L. Seymour, L. M. Nuwaysir, B. Lefebvre, F. Kuhlmann, J. Roark, P. Rainer, S. Detlev, T. Hemenway, A. Huhmer, J. Langridge, B. Connolly, T. Chadick, K. Holly, J. Eckels, E. W. Deutsch, R. L. Moritz, J. E. Katz, D. B. Agus, M. MacCoss, D. L. Tabb and P. Mallick, A cross-platform toolkit for mass spectrometry and proteomics, *Nat. Biotechnol.*, 2012, **30**, 918–920.
